# Cell wall arabinogalactan is responsible for Fungitell® cross reactivity in nocardiosis

**DOI:** 10.1101/2024.08.22.608962

**Authors:** Christophe Mariller, Pascal Letowski, Wei-Ting Chang, Todd L. Lowary, Marc Ulrich, Karine Faure, Séverine Loridant, Boualem Sendid, Frédéric Wallet, Daniel Poulain, Marc Hazzan, Marie Frimat, Yann Guerardel, Marie Titecat

**Affiliations:** Université de Lille, CNRS, UMR 8576 – UGSF - Unité de Glycobiologie Structurale et Fonctionnelle, F-59000 Lille, France; Institute of Biological Chemistry, Academia Sinica, Nangang, Taipei, 11529, Taiwan; Institute of Biochemical Sciences, National Taiwan University, Taipei, 10617, Taiwan; CHU Lille, Nephrology Department, F-59000, Lille, France; CHU Lille, Infectious Diseases Department, F-59000 Lille, France; CHU Lille, Institute of Microbiology, F-59000 Lille, France; Univ. Lille, CNRS UMR 8576, UGSF, Inserm U1285, CHU Lille, Laboratoire de Parasitologie-Mycologie, Lille, France; Univ. Lille, Inserm, Institut Pasteur de Lille, U1167 - RID-AGE, F-59000 Lille, France; Institute for Glyco-core Research (iGCORE), Gifu University, Gifu, Japan; Université de Lille, INSERM, CHU Lille, U1286-INFINITE-Institute for Translational Research in Inflammation, F-59000 Lille, France

**Keywords:** nocardiosis, cross reactivity, serum (1, 3)-β-D-Glucan, *Nocardia nova*

## Abstract

Nocardiosis is a serious infection in immunosuppressed patients, especially transplant recipients. The slow-growing phenotype of the bacterium and the variety of symptoms complicate diagnosis and delay antimicrobial therapy, which results in high mortality rates despite effective treatments. Incidentally, some nocardiosis patients test positive in fungal diagnostics that detect (1,3) β−D-glucan (the Fungitell® assay), but the basis for this cross-reactivity remains unknown. We demonstrate that nocardial cell wall arabinogalactan is a cryptic antigen responsible for cross reactivity in the Fungitell® assay and that this antigen is revealed *in vivo* following bacterial cell lysis. We further show that the reactivity results from β-glucose substitution of the galactan domain, a modification specific to Nocardia, and identify the optimal antigen as a tetramer of the trisaccharide repeating unit. By providing structural evidence for Fungitell® cross-reactivity during nocardiosis, this work paves the way for developing specific diagnostic tools that are presently lacking.

## MAIN

Nocardiosis is a life-threatening, opportunistic infection caused by *Nocardia* spp, a ubiquitous group of Gram-positive environmental bacteria. These organisms are related to mycobacteria through a common structural feature: an arabinogalactan polysaccharide domain that covalently connects a peptidoglycan layer to mycolic acid (lipid) chains ^1^. Among approximately 100 species, *N. nova* complex frequently causes disease in humans ^2–6^. Nocardiosis displays various presentations from localized (*i.e.* skin, lung) to disseminated (*i.e*. bacteremia, brain abscesses) forms and predominantly occurs in immunocompromised patients, especially those with cellular immune disorders with poor clinical outcome ^3,7^. The non-specific clinical features of nocardiosis (^8,9^ combined with the slow growing bacterial phenotype can dramatically delay diagnosis. This, in turn, delays the initiation of appropriate antimicrobial therapy – the use of imipenem plus amikacin ^10,11^ – an uncommon antimicrobial combination for the management of severe Gram-positive infections.

In addition to nocardiosis, the immunocompromised population is also at risk of developing opportunistic fungal infections with potential serious outcomes. Thus, in case of fever, these patients benefit from fungal antigen monitoring to predict and treat invasive disease without delay ^12,13^. Among these antigens, elevations of the β-(1,3)-D-glucan (BDG), a glucose polymer found in the cell wall of many fungal organisms (*i.e. Candida* spp., *Aspergillus* spp., *Pneumocystis* spp.) is indicative of fungal disease. Interestingly, positive BDG rates have been reported in various cases of evolutive nocardiosis ^14–18^. However, the mechanisms underlying how a bacterial species that does not produce BDG can give rise to a positive result in diagnostics that detect this antigen, have never been reported.

Starting from the observation of an apparent BDG elevation in a *N. nova*-infected patient, we identified the arabinogalactan (AG) fraction of the nocardial cell wall as the antigen resulting in BDG test positivity. These results were enhanced by the chemical synthesis of the pattern responsible for the cross reactivity, which both confirmed the structure and provide tools that may help in developing new diagnostic approaches for nocardiosis.

## RESULTS

### Case report

A 67-year-old kidney transplant patient was admitted with fever and a month-long ulceration on the right hand associated with an axillary lymph node following gardening (Fig. 1A). Immunosuppressive therapy included tacrolimus, mycophenolate mofetil, and steroids. Standard cutaneous and blood cultures yielded negative results, as did polymerase chain reaction (PCR) assays for *Bartonella* spp. and *Mycobacterium tuberculosis*. A Fungitell® serum test exhibited apparent BDG titers of 2000 pg/mL. Skin biopsy analyses showed nonspecific inflammation, without fungal filaments, and 18S PCR assay detected no fungal DNA. Fungitell positive samples tested negative for circulating trehalose (MS-DS), excluding invasive candidiasis ^19^. First-line therapy with amoxicillin/clavulanate and azithromycin failed to improve symptoms, with apparent BDG levels rising to 2326 pg/ml. Prolonged culture of the surgically removed axillary lymph node showed dry white colonies of branching gram-positive rods identified as *Nocardia nova* (reliability score of 1.95). Brain MRI revealed a cerebral abcess (Fig.1B). Appropriate antimicrobial therapy consisting imipenem (4 weeks) and oral trimethoprim-sulfamethoxazole (12 months), along with adjustments to the immunosuppressive regimen, led to symptom resolution (Fig. 1a, 1b) and decrease of the serum BDG concentrations. Patient features detailed in Supplementary Table 1.

**Figure 1.**
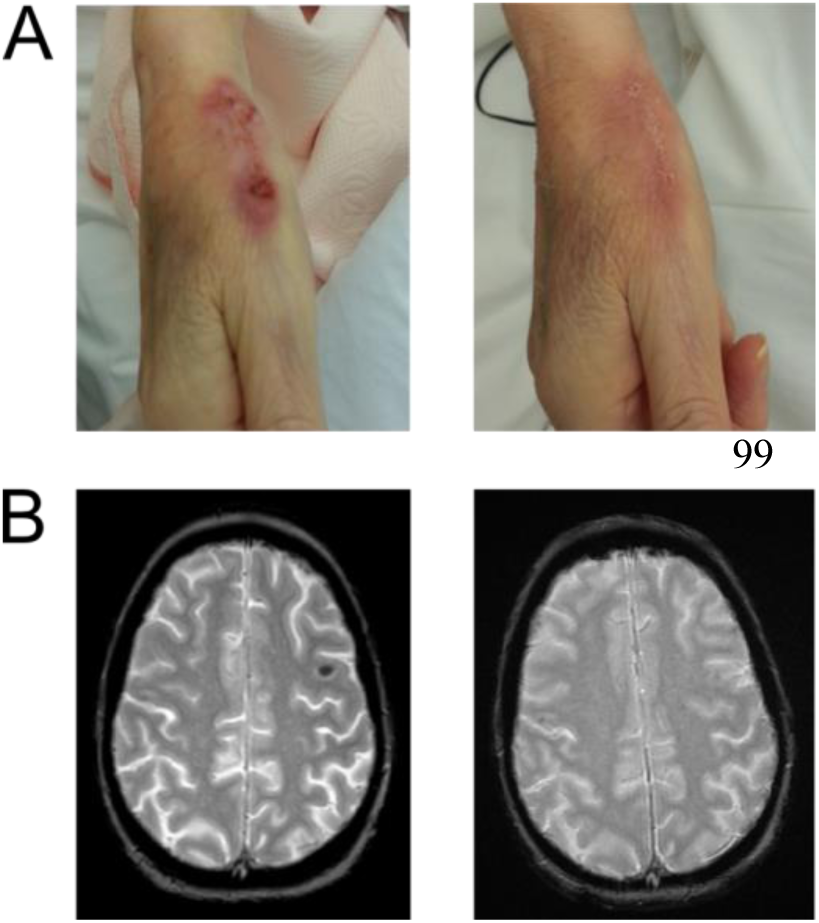
Patient clinical outcome under optimal antimicrobial therapy. The evolution of skin (A) and brain abscess (B) lesions in the patient were observed over 15 days and 1 month, respectively.

### The positive Fungitell® result is not recovered by direct Nocardia cross-reactivity testing

Because the apparent BDG elevation was not associated to any evolutive fungal disease, the *N. nova* clinical strain was directly tested for the possibility of cross reactivity with the Fungitell® assay (Fungitell®, Cape Cod) according to the Koncan *et al.* procedure ^14^. Bacterial cells were washed seven times and the inoculum adjusted to 6 × 10^7^ CFU/mL in glucan free water. Application of the Fungitell® test to the suspensions provided negative results even after pellet grinding. A second set of experiments consisting of bacterial growth in glucan-free serum of donor secondarily tested with the Fungitell® assay remained negative. These findings invalidated the possibility that the interaction between serum proteins and *N. nova* cell wall elements led to a false positive Fungitell® result. Further unsuccessful experiments comparing different culture conditions were also conducted: solid *versus* liquid media, “young” versus “old” bacterial suspensions, CO_2_ enriched atmosphere or not. None of these conditions led to reproducing the Fungitell® result arising from the patient’s sera (data not shown).

### Nocardia nova cell wall extraction is required to identify the positive cell wall pattern

Because the lack of Fungitell® reactivity of intact and ground bacterial pellets apparently contradicted the presence of a cell-surface cross-reactive component, we hypothesized that the component may be unmasked during the infection process. Thus, to identify potential cryptic Fungitell®-reactive motifs, the cell wall components were sequentially extracted from intact bacteria and individually tested. The *N. nova* cell wall was fractionated following the procedure of Besra and coworkers ^20^ from a 1196 mg sample of lyophilised bacteria cultivated in Sauton broth (Fig. 2A). Fungitell® reactivities of each fraction in pg of β-glucan per mL equivalent are reported in Figure 2B.

**Figure 2.**
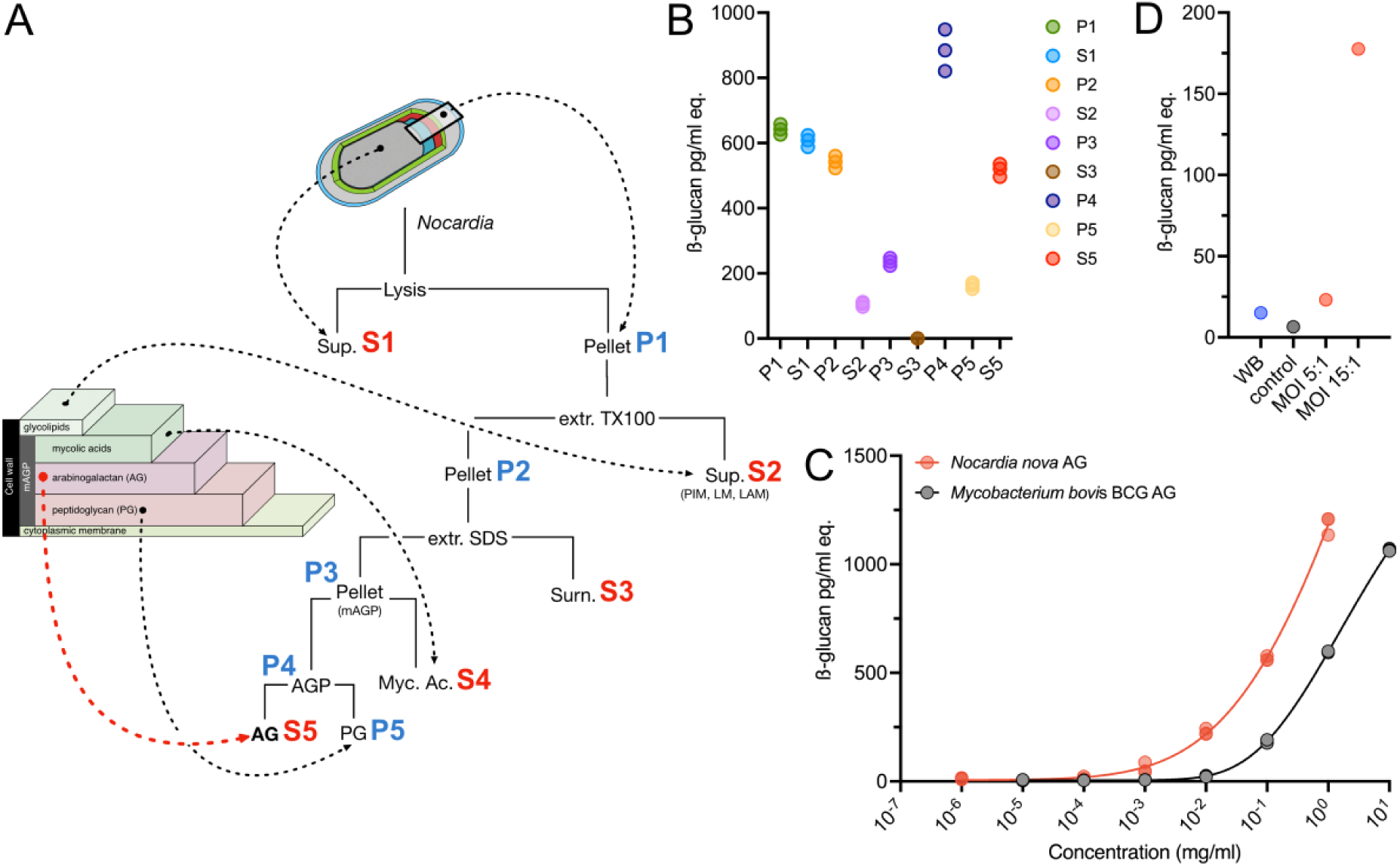
Subcellular fractionation of N. nova and BDG assays. (A) Cell wall components were separated and purified from bacteria through sequential extractions using Triton X100, hot SDS and 2% KOH and centrifugation. (B) Individual fractions were tested using the Fungitell® assay and apparent BDG concentrations are expressed in pg/mL eq. Purified AGP (P4) and AG (S5) show high reactivity to the Fungitell® assay (C) AG isolated from *N. nova* shows much higher Fungitell® reactivity than AG isolated from *M. bovis* BCG. (D) Following infection by *N. nova* (MOI 1:5 and 1:15) and processing by macrophage-like THP-1, the culture medium is highly reactive to the Fungitell® assay compared with non-infected THP-1 (control) and intact, whole bacteria *N. nova* (WB).

Mechanical lysis of the bacteria in PBS and subsequent centrifugation generated, on the one hand, a supernatant fraction (S1) mainly composed of soluble proteins and a complex mixture of surface components and, on the other hand, a pellet P1. Both S1 and P1 exhibited significant reactivity to the Fungitell® assay. At this stage, the S1 fraction was disregarded to avoid possible contamination from surface-associated molecules to focus on bacteria-associated molecules. This was deemed a valid approach as the bacteria themselves were unreactive (above), suggesting a surface-associated molecule was not the cross-reactive species. The reactive P1 fraction was thus further treated with 2% Triton X-100 to extract amphiphilic glycolipids including phospho-inositol mannosides (PIMs), lipomannan (LM) and lipoarabinomannan (LAM) in the S2 fraction that exhibited low reactivity to the Fungitell® assay. In contrast, the resulting pellet fraction (P2) showed very high Fungitell® reactivity, comparable to P1, demonstrating that the cross-reactive motif present in P1 was retained in this fraction. Successive extractions of cell-wall associated proteins with hot SDS (P3 and S3) and saponification of mycolic acids using 2% KOH generated an arabinogalactan-peptidoglycan (AGP) fraction (P4) and a lipid fraction that contained mycolic acids (S4). Purified AGP devoid of mycolic acids showed the highest reactivity to Fungitell® of all fractions. Finally, dissociation of peptidoglycan (PG) from AGP fraction (P4) led to the isolation of a highly purified arabinogalactan (AG) fraction (S5) that could be structurally defined. S5 has retained a high value for apparent BDG (535 pg/mL) albeit slightly lower than P4. In comparison, the PG fraction showed much lower reactivity.

Considering the high reactivity of the AG fraction, we assayed AG that was purified from *Mycobacterium bovis* BCG through the same procedure. At equivalent concentration, the AG extracted from *M. bovis* BCG demonstrated significantly reduced reactivity, with a concentration shift of over one logarithmic unit compared to the one purified from *M. nova* (Fig 2C). Taken together, these experiments demonstrated that the reactivity of *N. nova* is triggered exclusively after bacterial lysis and that the cell-wall AG is the most likely component responsible for the Fungitell® reactivity.

AG is normally inaccessible when buried in the bacterial cell wall. It is therefore reasonable to hypothesize that the otherwise cryptic polysaccharide epitopes are exposed after lysis of *N. nova* by immune cells *in vivo*. To test this hypothesis *in vitro*, THP-1 macrophages were infected with labeled *N. nova* at MOIs of 5:1 and 15:1. After six hours of infection followed by a twelve hours macrophage incubation, the proportion of phagocytes observed reached 30% with a maximum of two bacteria per cell. At the end of the incubation period, the reactivity of the resulting medium toward β-glucans was evaluated with the Fungitell® assay, and compared to medium from control experiments with THP-1 cells alone and *N. nova* alone. As shown in Figure 2D, the media collected after infection of THP-1 with *N. nova* showed a much higher reactivity toward β-glucans, in particular at a MOI of 15:1, strongly suggesting that *N. nova* antigens must be processed by macrophage-induced cell lysis to reveal the cross-reactive antigen, consistent with the lack of reactivity of intact bacteria.

### The arabinogalactan isolated from Nocardia nova is substituted with β-D-glucopyranose residues

To understand the mechanism that underpins the unexpected and specific cross-reactivity of the AG containing fraction isolated from *N. nova* (NnAG) in the Fungitell® test, the structure of the reactive fraction (S5) was investigated in detail. AG is the central component of the complex cell wall of Corynebacteriales, including the *Mycobacteriaceae*, *Corynebacteriaceae* and *Norcardiaceae*. Although the structure and biosynthetic pathways of AG have been thoroughly studied in a number of mycobacterial species, information about its composition and fine structure in nocardia is scarce. Earlier preliminary studies by Daffe *et al.* suggest that *Nocardia asteroides* and *Nocardia brasiliensis* do synthesize AG with structures that differ significantly from those produced by mycobacteria ^1^. Monosaccharide composition analysis of fraction S5 isolated from *N. nova* showed that NnAG was mostly composed of arabinose (Ara) and galactose (Gal), similar to the AG isolated from *Mycobacterium bovis BCG* (MbAG) as shown in Table 1A ^21^. Furthermore, it contained a small amount of rhamnose (Rha) and *N*-acetyl-glucosamine (GlcNAc), strongly suggesting that NnAG is anchored to peptidoglycan through the Rha*p*-(1,4)-Glc*p*NAc-P disaccharide as in mycobacteria ^22^. However, unlike mycobacterial AG, NnAG also contained a significant amount of glucose (Glc, 8.5%). Linkage analysis of the monosaccharides established that NnAG contains terminal, 2-, 5-, 3,5- and 2,5-linked Ara in furanose forms (Ara*f*), indicating that the arabinan domain of NnAG shows strong similarities with that of MbAG (Table 1B). The Gal residues were observed as 5Gal*f* and 5,6Gal*f*, which demonstrates that the galactan domain of NnAG differs from MbAG, which is composed of a linear chain of alternating 6Gal*f* and 5Gal*f* substituted in 6 position by the arabibnan chain. Finally, all Glc residues were identified as unsubstituted in the pyranose form (Glc*p*) revealing that they substitute AG in terminal positions.

**Table 1.**
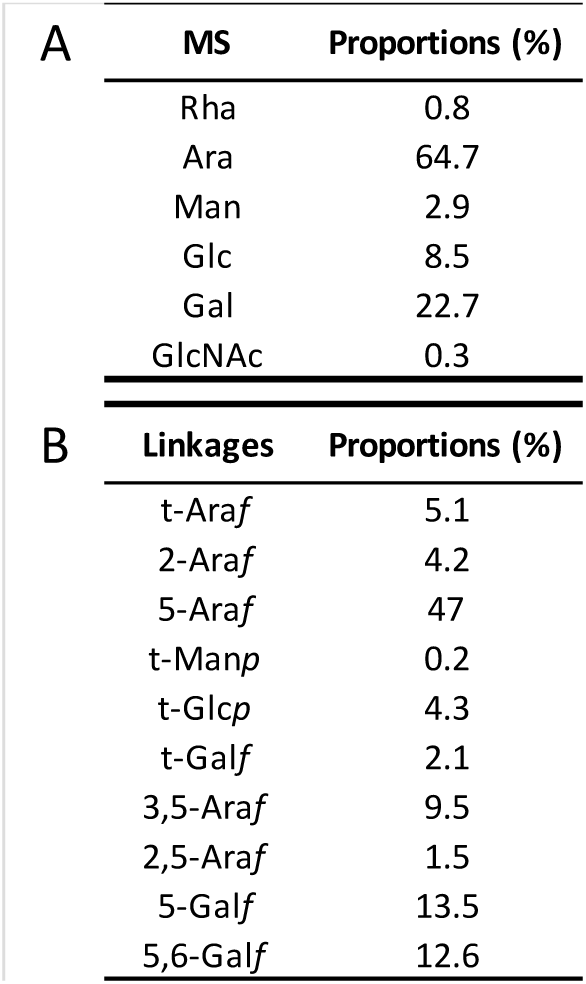
Monosaccharide composition analysis of arabinogalactan (AG) isolated from N. nova (NnAG). (A) GC-FID identification and relative quantification of reduced-peracetylated monosaccharides (MS) showed that NnAG was composed primarily of arabinofuranose (Ara*f*), galactofuranose (Gal*f*) and glucopyranose (Glc*p*). (B) GC-MS identification and relative quantification of partially methylated and acetylated reduced derivatives of major monosaccharides, showed that NnAG shared most of the structural features of AG isolated from mycobacteria, except for the presence of Gal*f* disubstituted at both the 5 and 6 positions in the same proportion than 5Gal*f*.

NnAG was further characterized by high field NMR spectroscopy. Spin systems and chemical shifts obtained through a combination of homonuclear ^1^H–^1^H experiments (COSY, TOCSY) and heteronuclear ^1^H–^13^C experiments (HSQC, HSQC–TOCSY and HMBC) permitted the identification of seven types of monosaccharides in accordance with the composition and linkage analyses as shown in the detail of ^1^H–^13^C HSQC experiments (Fig. 3) and reported in Table 2. The polysaccharide was compared to the AG isolated from *M. bovis* BCG, which is known to be identical to AG from *M. tuberculosis* and all mycobacterial species studies reported to date ^21^. When compared with MbAG, NnAG exhibited marked differences in both the arabinan and galactan domains.

**Figure 3.**
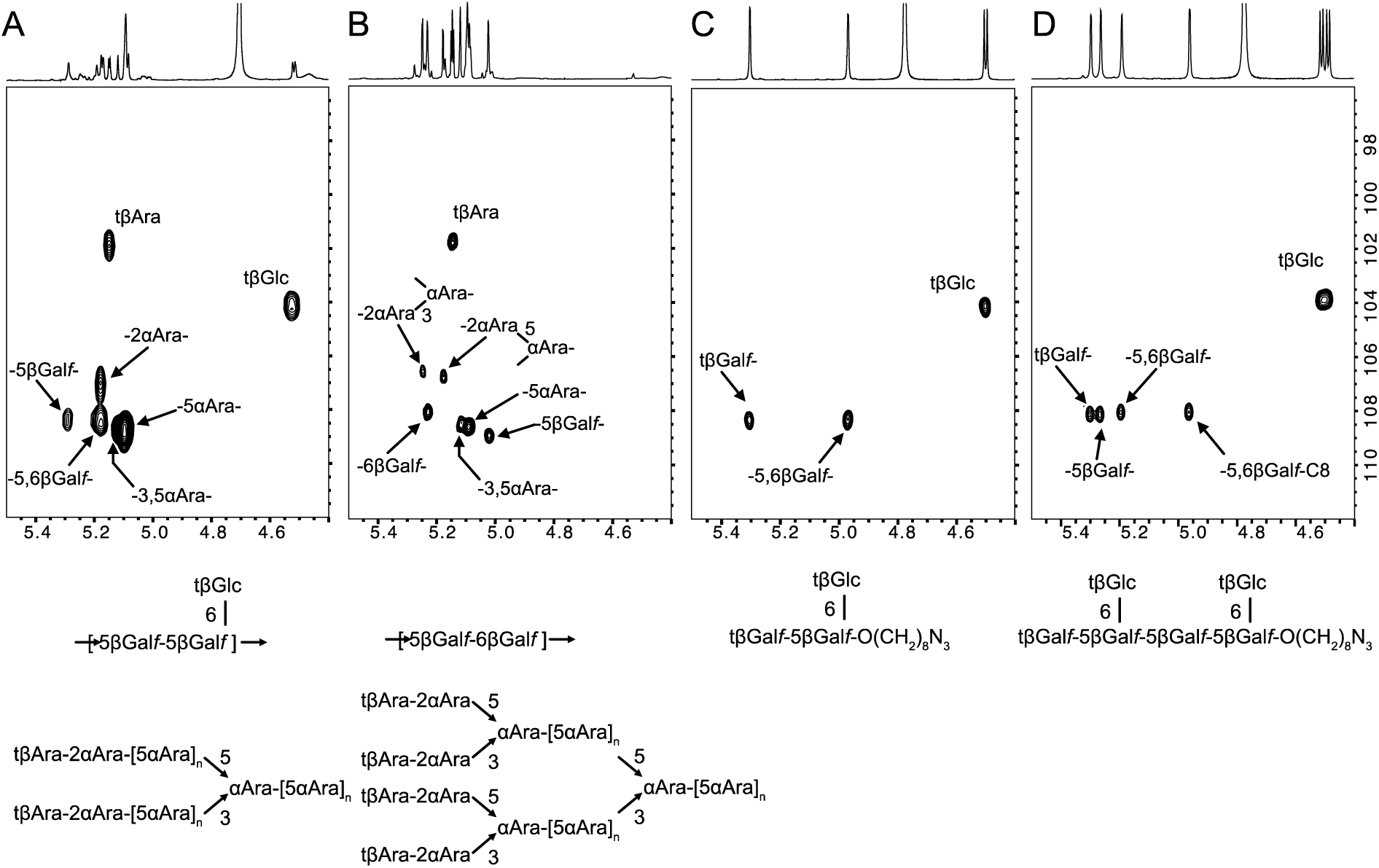
Comparison of the main structural features of NnAG and MbAG. Anomeric regions of the ^1^H– ^13^C HSQC NMR spectra of NnAG (A top panel) and MbAG (B top panel) demonstrate that AG isolated from *N. nova* and *M. bovis* differ in the structure of both the arabinan and galactan moieties (A and B lower panels). In particular, the presence of an additional βGlc residue substituting a 5,6-αGal*f* residue in NnAG was established by comparison of ^1^H–^13^C HSQC and ^1^H–^13^C HMBC (Fig. 4 and SupFig. 1) spectra. The exclusive presence of βGal*f*-5(βGlc*p*-6)βGal*f* epitope on NnAG was confirmed by comparison with the NMR spectra of synthetic compounds βGal*f*-5(βGlc*p*-6)βGal*f*-O(CH_2_)_8_N_3_ (**NnAG1**) and [βGal*f*-5(βGlc*p*-6)βGal*f*]_2_-O(CH_2_)_8_N_3_ (**NnAG2**), the synthesis of which is described below.

**Table 2.**
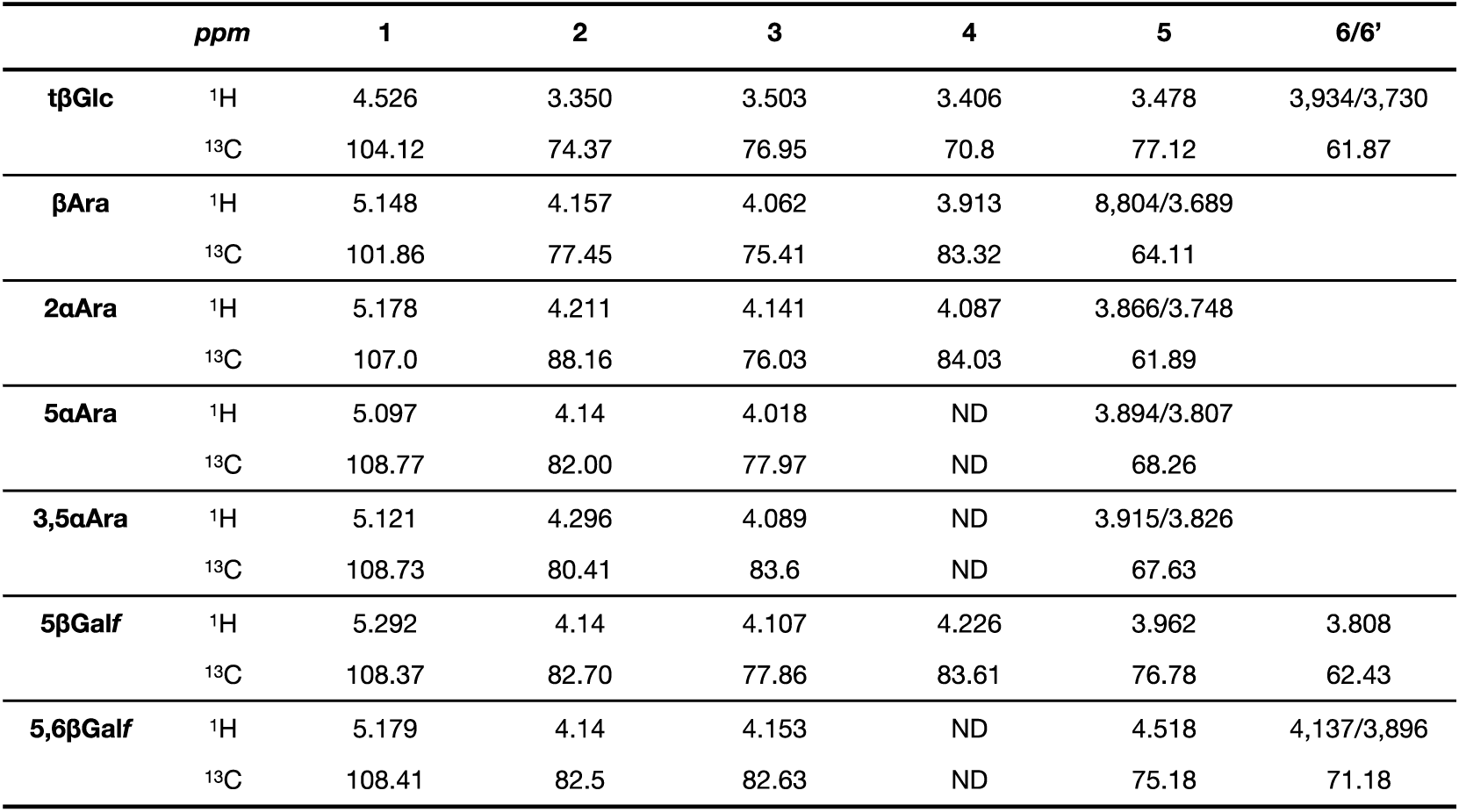
^1^H and ^13^C chemical shifts of main monosaccharides of NnAG derived from ^1^H–^1^H TOCSY, ^1^H–^13^C HSQC, ^1^H–^13^C HSQC–TOCSY and ^1^H–^13^C HMBC NMR experiments.

Notably, although NnAG and MbAG share common features such as tβAra, 2αAra, 3αAra, 5αAra and 3,5αAra, NnAG contains a single signal associated with 2αAra whereas MbAG contains two. The ^1^H–^13^C HMBC spectrum of NnAG showed an intense ^3^*J*_H,C_ connection between 2αAra-H1 at 5.178 ppm and 5αAra-C5 at 68.26 ppm whereas (Fig. 4) The ^1^H–^13^C HMBC spectrum of MbAG showed that the two 2αAra-H1 had differential ^3^*J*_H,C_ connections with 3,5αAra-C3 and 3,5αAra-C5 (data not shown), demonstrating that NnAG lacks the tβAra-2αAra-3αAra branch that is present in the arabinose capping of MbAG as depicted in Fig 3, making the arabinan domain simpler. NMR experiments confirmed the presence of 5βGal*f* at δ ^1^H/^13^C 5.292/108.37 and 5,6βGal*f* at δ ^1^H/^13^C 5.179/108.41 in NnAG instead of the alternating 5βGal*f* and 6βGal*f* observed in MbAG. The ^1^H–^13^C HMBC spectrum established that 5βGal*f* and 5,6βGal*f* were connected to each other through (1,5) linkages based on the strong 5βGal*f-*H1/5,6βGal*f*-C5 and 5,6βGal*f-*H1/5βGal*f*-C5 ^3^*J*_H,C_ cross signals (Fig. 4).

**Figure 4.**
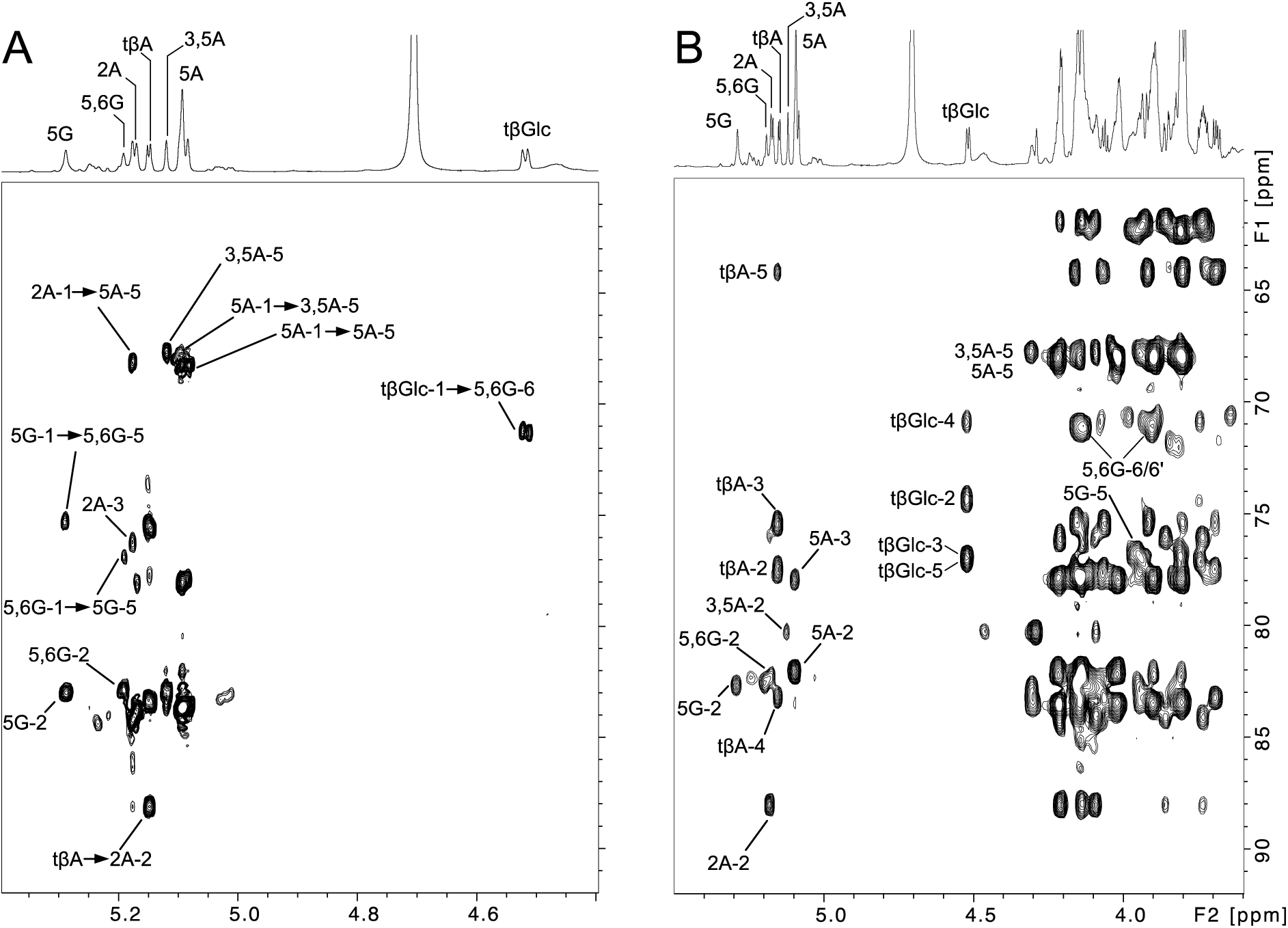
Identification of the linkage pattern of arabinan and galactan moieties of NnAG. Comparison of the ^1^H–^13^C HMBC (A) and ^1^H–^13^C HSQC–TOCSY (B) spectra of NnAG demonstrated that the tβGlc residue is linked to the C6 position of a 5,6-βGal*f* (5,6G) residue. This analysis also demonstrated that the 2αAra residue exclusively substitutes the C5 position of the arabinan chain contrary to MbAG in which Ara residues can be di-substituted at the C3 and C5 positions.

In addition, the ^1^H–^13^C HSQC spectrum of NnAG shows an intense signal at δ ^1^H/^13^C 4.526/104.12 that was further identified as a tβGlc*p* residue owing to its characteristic spin system (Table 3). A strong ^3^*J*_H,C_ cross signal between tβGlc*p*-H1 and 5,6βGal*f*-C6 (Fig. 4) unambiguously demonstrated that the βGlc residue substitutes the galactan moiety at the O6 position to generate a repetitive –[5βGal*f*-5(βGlc*p*-6)βGal*f*]_n_– motif. To confirm the structural attribution of the new Gal*f* backbone observed in NnAG, a series of oligosaccharide analogues including the trisaccharide Gal*f*-5(βGlc*p*-6)βGal*f*-O(CH_2_)_8_N_3_and the hexasaccharide [Gal*f*-5(βGlc*p*-6)βGal*f*]_2_-O(CH_2_)_8_N_3_ were synthesized (details below) and analyzed by NMR spectroscopy as shown in SupFig1 and SupFig2. Of particular interest, the tβGlc*p* residues in both compounds showed identical NMR parameters to NnAG (Fig. 3) confirming its structure. The presence of a backbone composed of alternating 5βGal*f* and 5,6βGal*f* residues in NnAG was also confirmed by identical spin systems and chemical shifts compared to the internal 5βGal*f* and 5,6βGal*f* residues of the synthetic hexasaccharide (Figure 3D).

Altogether, detailed structural analysis of the *N. nova* Fungitell®-reactive fraction demonstrated that NnAG differs from mycobacterial AG in both arabinan and galactan domains. Of particular interest, the galactan domain is characterized by a β-glucose containing –[5βGal*f*-5(βGlc*p*-6)βGal*f*]_n_– repeating unit instead of a –[5βGal*f*-6βGal*f*]_n_– motif. Because the Fungitell® assay detects BDG, which is composed of β-glucose residues, we hypothesize that this structural difference, in particular the β-Glc*p* side chains, was key to understanding the cross reactivity in the Fungitell® test, which was further assessed as described below.

### Synthesis of repeating units of the NnAG galactan confirms the structure of the native polysaccharide and provides compounds for additional evaluation

To confirm the structure of the *N. nova* AG galactan as determined by NMR spectroscopy (see above) and to provide a series of structurally-defined compounds that could be evaluated in the Fungitell® assay, six fragments of the polysaccharide (**NnAG1**–**NnAG6**, Figure 5) were synthesized. The targets consist of 1**–**6 NnAG galactan repeating units, each functionalized with an azidooctyl aglycone to facilitate their future coupling to other species, *e.g.,* proteins for the generation of monoclonal antibodies. We envisioned that **NnAG1**–**NnAG6** could be obtained from three monosaccharide building blocks: thioglycoside **1** and glycosyl fluorides **2** and **3** (Figure 5).

**Figure 5.**
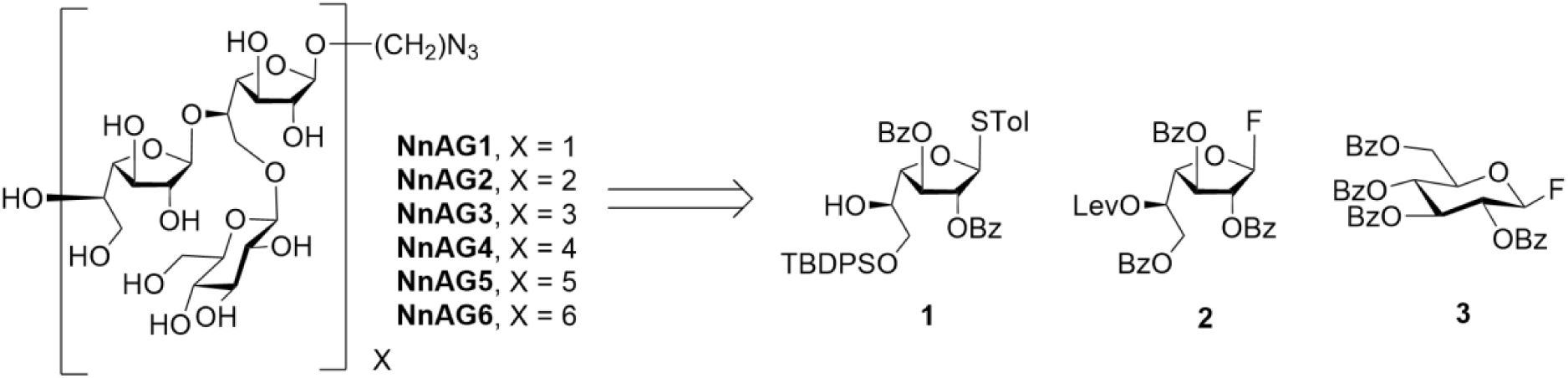
Synthetic NnAG galactan fragments **NnAG1**–**NnAG6** and the three monosaccharides (**1**–**3**) used for their preparation.

The synthesis of **1**–**3** is shown in Scheme 1. Access to **1** started with the known thioglycoside **4** ^23^ (Scheme 1A). Benzoylation of the diol in **4** and acid hydrolysis of the isopropylidene ketal provided an 83% yield of **5**, which was then treated with a limiting amount of *t*-butylchlorodiphenylsilane (TBDPSCl) in pyridine to generate **1** in 77% yield. Preparing **2** (Scheme 1B) started from **6** ^24^, which was first treated with *p*-toluenesulfonic acid (*p*-TsOH) in methanol and dichloromethane, triggering removal of the trityl either and subsequent migration of the O-5 benzoyl group to O-6. The intermediate alcohol formed in this process underwent Steglich esterification with levulinic acid affording an 83% yield of **7** over the two steps. Reaction of this thioglycoside with *N*,*N*-diethylaminosulfurtrifluoride (DAST) and *N*-bromosuccinimide (NBS) provided glycosyl fluoride **2** in 97% yield. Finally, the synthesis of **3** (Scheme 1C) was carried out starting from the perbenzoylated glucose derivative **8** ^25^ in two steps: conversion to thioglycoside **9** and then to the glycosyl fluoride under standard conditions to obtain the product in 63% overall yield.

**Scheme 1.**
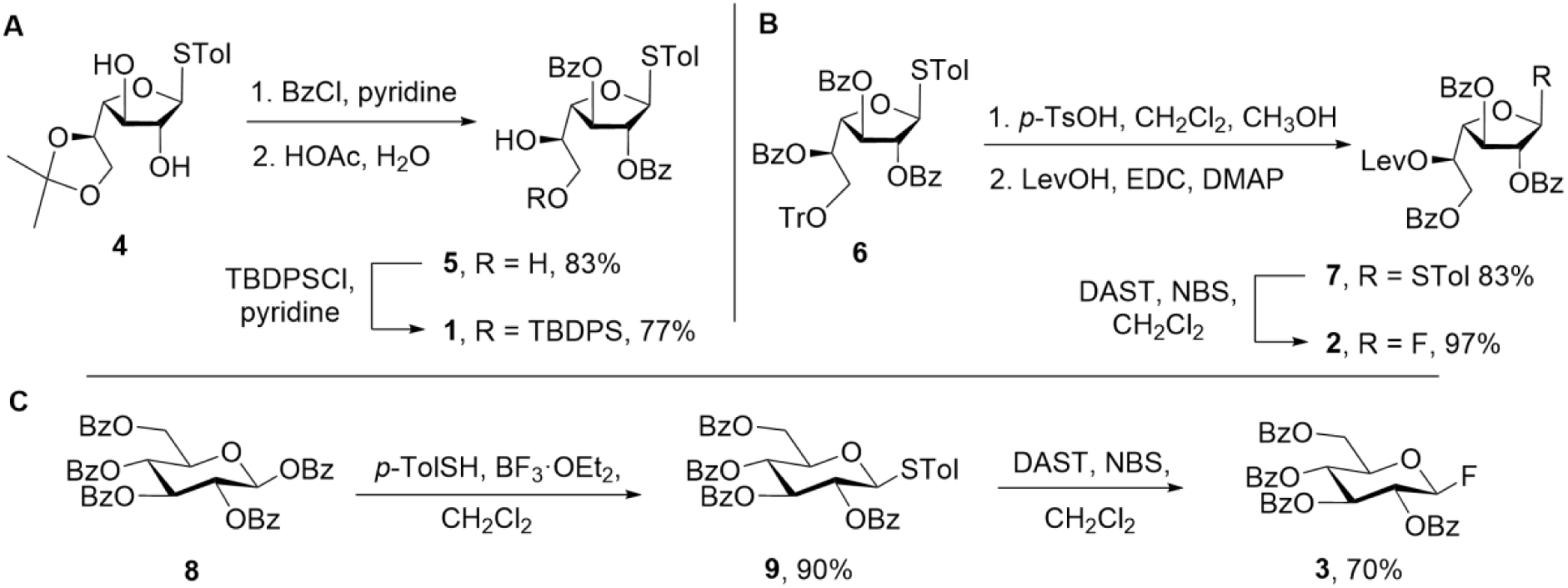
Synthesis of building blocks **1**–**3**.

These three monosaccharides were then used to generate a trisaccharide derivative (Scheme 2) that could be employed in the assembly of the target polysaccharide fragments. Initially, thioglycoside alcohol **1** and glycosyl fluoride **2** were coupled upon treatment with zirconocene dichloride (Cp_2_ZrCl_2_) and silver triflate (AgOTf) giving disaccharide **10** in 92% yield. Removal of the silyl ether protecting group afforded an 82% yield of disaccharide alcohol **11**. Upon glycosylation of **11** with glycosyl fluoride **3**, again promoted by Cp_2_ZrCl_2_ and AgOTf, an 87% yield of trisaccharide **12** was obtained. With this compound in hand, we explored its ability to act as a glycosyl donor via coupling with 8-azidooctanol using *N*-iodosuccinimde (NIS) and AgOTf as the promoters. This reaction cleanly provided the expected trisaccharide, **13** (99% yield). This result suggested that **12** could be efficiently used to access the larger fragments through an iterative process of selective cleavage of the levulinyl (Lev) ester and glycosylation.

**Scheme 2.**
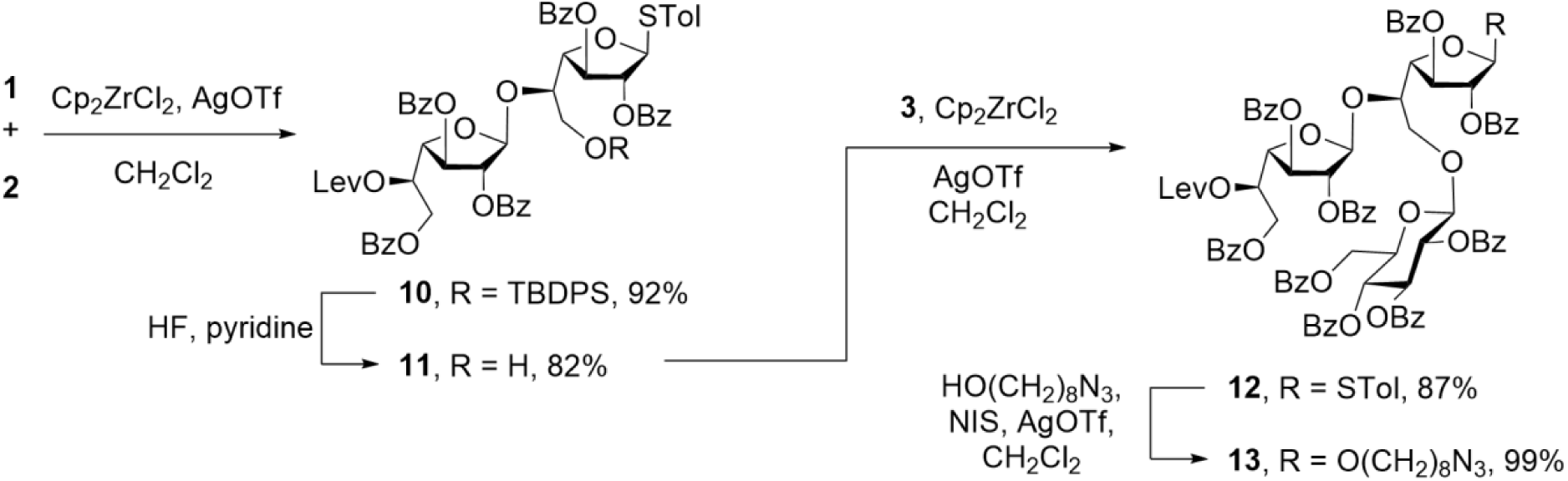
Synthesis of trisaccharide **13**.

The application of the iterative process described above to the synthesis of **NnAG1**–**NnAG6** is shown in Scheme 3. Treatment of **13** with hydrazine acetate (NH_2_NH_2_·HOAc) selectively cleaved the Lev ester leading to an 82% yield of trisaccharide **14**. Next, NIS/AgOTf-promoted glycosylation of **14** with **12** provided the expected hexasaccharide **15** in 73% yield. This cycle was applied one more time (**15**→**16**→**17**) in comparable yields; however, attempted glycosylation of nonasaccharide **18** with **12** failed to give the expected dodecasaccharide **19**. Fortunately, the use of glycosyl fluoride **24** (Scheme 3, obtained in 99% yield by treatment of **12** with NIS and DAST) as the glycosyl donor gave **19** in 85% yield from **18**. Further iteration of the cycle using **24** yielded the desired targets with 15 and 18 monosaccharide residues; **21** and **23**, respectively. Access to **NnAG1**–**NnAG6** was achieved in 72–92% yields by reaction of **13**, **15**, **17**, **19**, **21** or **23** with sodium methoxide in methanol.

**Scheme 3.**
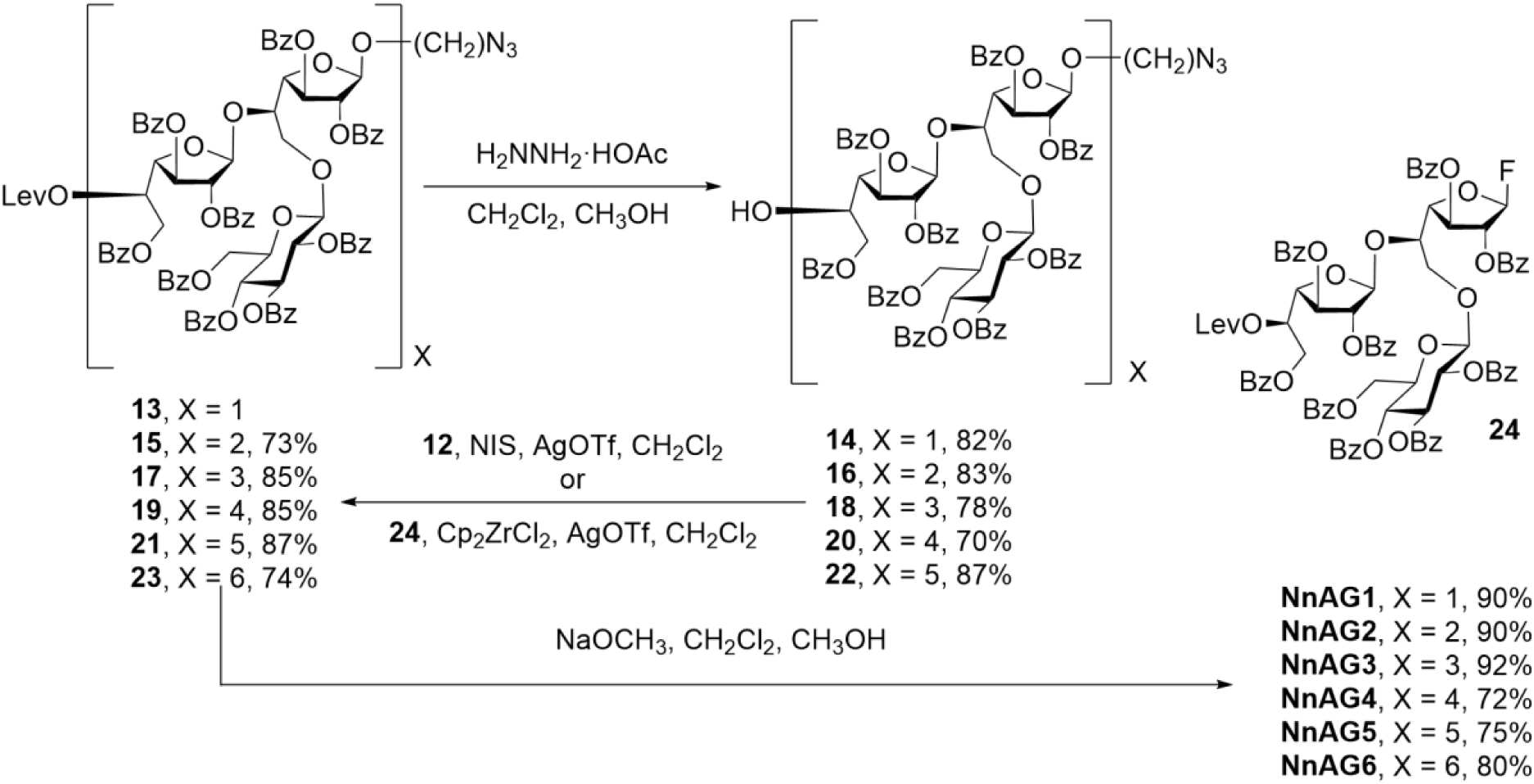
Synthesis of **NnAG1**–**NnAG6** via an iterative process of Lev ester cleavage and glycosylation followed by deacylation.

As outlined above, the NMR data obtained from trisaccharide **NnAG1** and dodecasaccharide **NnAG2** agreed well with those obtained on the NnAG galactan domain, thus supporting the NMR structural assignments (Fig.3).

### The β-glucose residues substituting the N. nova arabinogalactan are responsible for Fungitell*®* cross-reactivity

To confirm that the –[5βGal*f*-5(βGlc*p*-6)βGal*f*]_n_– motif observed in NnAG is indeed responsible the cross reactivity, the responsiveness of synthetic analogues **NnAG1**–**NnAG6** to Fungitell® were evaluated in comparison with purified NnAG. Screening at two concentrations showed that all synthetic compounds were reactive at the highest concentration, but only **NnAG4** showed significant reactivity equivalent to 1 ng of β-glucan/ml eq. at the lowest concentration and a very robust reactivity at the highest concentration, which are very similar to natural AG purified from *N. nova* (Fig. 6A). This was confirmed by comparing the dose-dependent reactivity of **NnAG4** and natural *N. nova* AG, which showed very similar patterns (Fig. 6B).

**Figure 6.**
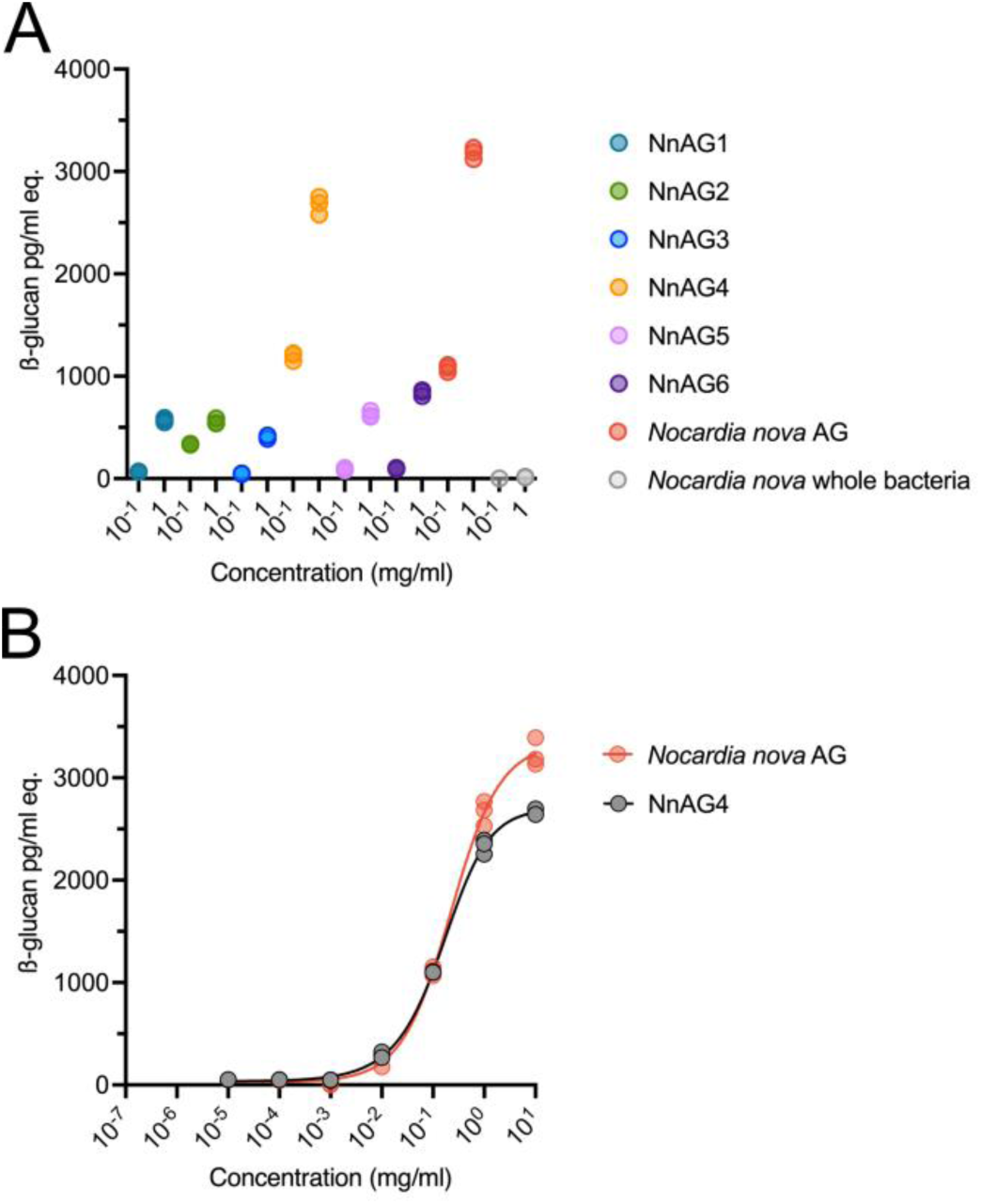
Comparison of BDG reactivities of NnAG and synthetic oligosaccharides −5[(βGlcp-6)βGalf-5βGalf]-_n_. (A) Out of the six synthetic analogues of *N. nova* AG fragments, the tetramer oligosaccharide (**NnAG4**) showed the highest reactivity to the BDG assay, equivalent to the natural AG at two different concentrations. (B) Concentration dependent assays showed a very similar trend for **NnAG4** and natural *N. nova* AG (NnAG).

In conclusion, the synthetic dodecasaccharide βGal*f*-5(βGlc*p*-6)βGal*f*-[5βGal*f*-5(βGlc*p*-6)βGal*f*]_3_-(CH_2_)_8_N_3_ (**NnAG4**), mimicking the galactan domain of NnAG, shows a reactivity level in the Fungitell® assay comparable to intact NnAG. This demonstrates that the substitution of four βGlc*p* residues along the galactan backbone is sufficient to trigger a positive result in the assay.

## DISCUSSION

The mortality rate of nocardiosis is high, estimated at 16.2% in European organ transplant recipients ^26^ and 17.5% in hematopoietic stem cell transplant cohorts ^27^. As such, there is an urgent need to identify methods for faster diagnosis and earlier treatment initiation in these vulnerable populations. These patients are also at risk to develop fungal infections justifying BDG testing in case of fever. Because various studies report positive BDG titers during *Nocardia* spp. infection ^14,15,17,28^, discrepant experimental results may question this cross reactivity ^14,15,28^ and the reliability of this marker in evolutive cases of nocardiosis ^29^.

Detection of BDG in patient serum relies on its ability to activate the horseshoe crab coagulation cascade, which, in turn, cleaves an artificial substrate enabling colorimetric or turbidimetric detection ^30^. Several other BDG assays have been developed that differ in their threshold of detection, horseshoe crab species and detection method ^31^. Regardless of the assay, direct BDG dosage performed on whole bacteria cells were negative both in the present *N. nova* infection case using the Fungitell® assay and in a reported *N. farcinica* infection case using the Wako assay (Wako; Wako Pure Chemical Industries, Ltd., Tokyo, Japan) ^15^. We demonstrate here that a mechanical lysis step is required to detect a positive signal (641 pg/mL) and that a chemical cell wall extraction procedure made it possible to localize at this level (560 pg/mL). The first step was necessary to uncover the reactive structure buried in the bacterial cell wall but was insufficient to reach high levels in patient sera (2600 pg/mL).

The genus Nocardia belongs to the Actinomycetes family, and is structurally related to Mycobacteria ^1^. Its cell wall structure is characterized by a peptidoglycan layer covalently connected *via* an AG polysaccharide to a lipid layer of mycolic acids. Owing to the similarity between *Nocardia* spp. and *Mycobacteria* spp., we followed a known cell wall fractionation protocol ^20^ to characterize precisely the structure responsible for the Fungitell® cross reactivity. The *N. nova* AG was identified as the key bacterial fraction responsible for the high apparent BDG rates. As previously described by Daffé et *al.*, *N. asteroides* and *N. brasiliensis*, AG structures do not have β-glucan architecture but rather possess a linear chain of galactofuranose with side-chain β-glucopyranose appendages. Comparison of the GC/FID analysis of glycosyl linkage for NnAG, compared to that published for *N. asteroides* and *N. brasiliensis*, showed a similar proportion of glucopyranose: 8.5% (NnAG) versus 10.2% (*N. asteroides* AG) and 10% (*N. brasiliensis* AG). Consistent with Daffé et al., NMR experiments performed on *N. nova* AG assigned a β-D-configuration to the glucosyl residue and its terminal location on the AG chain. By testing purified extracts of AG in the Fungitell® assay, NnAG proved to be more reactive than MbAG with a one-log shift in dose-response curves. The structure of NnAG galactan domain was confirmed by the chemical synthesis of structurally defined fragments, up to six repeating units (18 monosaccharide residues) in length, whose NMR spectra matched those of the native polysaccharide.

To confirm that NnAG is indeed the endogenous molecule responsible for the BDG reactivity, the synthetic analogues (**NnAG1–NnAG6**) of the NnAG galactan, the purity and integrity of which were determined by high resolution NMR spectroscopy, were evaluated in the Fungitell® assay. Remarkably, the synthetic analogue **NnAG4**, a dodecasaccharide consisting of four repeating units exhibited a strong cross reactivity with BDG that was comparable to native NnAG, strongly suggesting that the substitution of the galactan domain by βGlc*p* residues was indeed responsible for the observed *in vivo* reactivity. These results provide, for the first time, direct biochemical proof that a purified cell wall component of *N. nova* cross reacts in assays designed to detect BDG. Surprisingly, although it was not negligible, the reactivity of longer polymers (**NnAG5** and **NnAG6**) was lower that of **NnAG4**. This finding demonstrates that the Fungitell® assay has a very restricted specificity toward NnAG, possibly resulting from the in-solution conformation of the βGlc epitopes along the galactan backbone.

*Nocardia* spp. strains are known to interact with host macrophages during infection and, in some cases, to evade the phagocytosis process ^32,33^. In an attempt to explain AG release leading to a BDG positive signal in patients, we performed macrophage infection experiments. High levels of apparent BDG were recorded after 12 hours of incubation with values increasing according to the initial bacterial load (C2 > C1). These observations explain the high levels observed in the patient described here with an uncontrolled infection site (brain abscess). In our experiments, *in vitro* rates decreased after 24 h of incubation probably because of macrophage necrosis and bacterial degradation. These results may suggest a transitory phenomenon that can also be balanced by an unknown AG half-life in the host blood stream, and consequently explain the few Fungitell® positive reported cases to date.

We demonstrate here that apparent BDG positivity in *N. nova* infection is not an incidental cross-reaction but relies on specific bacterial cell wall structures. Structural analysis of *N. nova* AG combined with macrophage infection experiments supported our hypothesis that Fungitell® assay positivity in *N. nova* infections arises as a result of cross-reactivity with NnAG. Thus, an isolated apparent BDG elevation, without evidence of any fungal infection, should be considered as a potential marker of nocardiosis for clinicians and microbiologists. The availability of synthetic NnAG chemical analogues (**NnAG1**–**NaAG6**) paves the way for the development of specific diagnosis tools (*e.g.,* monoclonal antibodies), that could be used in clinical settings.

## METHODS

### Patient data and bacterial strain

The patient was diagnosed with a *Nocardia nova* infection at University Lille Hospital’s Nephrology and Transplantation Department. Prior informed consent was obtained for the use of clinical data. The infective strain was isolated from positive blood and lymph node culture. Bacterial identification was performed by MALDI TOF mass spectrometry (Bruker Daltonics, Wissembourg, France). An accurate identification to the species level was given by a score value ≥1.9. The bacterial strain was sent to the French Referent Center for nocardiosis to confirm identification and perform antimicrobial susceptibility testing. For further experiments, the *N. nova* strain was stored at –80 °C in Müeller–Hinton medium supplemented with 10% glycerol until use.

### Fungitell assay procedures

The Fungitell® assay (Associates of Cape Cod, E. Falmouth, MA, USA) was performed according to the manufacturer’s recommendations using patient sera; a cutoff value of 80 pg/mL apparent BDG was used as a positive result. In cases in which apparent BDG levels exceeded 500 pg/mL, samples were diluted and retested. For bacterial experiments, *N. nova* strain was suspended in glucan-free sterile saline, pelleted and washed according to the Koncan *et al*. procedure ^14^. Serum culture assays were performed in BDG-negative serum from a healthy donor of the Etablissement Français du Sang (EFS). For dose response experiments involving arabinogalactan extracts of *N. nova, Mycobacterium bovis* BCG and synthetic fragments, concentrations of the different fractions were standardized to 1 mg/ml and ranged from 0.01 µg/mL to 10 mg/mL to determine dose response relationship between AG concentration and Fungitell® positivity. The interpretation of the Fungitell® assay required adjustment for non-BDG samples. To address saturation issues, kinetic data, comprising both standards and samples, were analyzed to determine the slope of absorbance change at 405 nm over time (SupFig. 3, 5 7 and 9). Using BDG standards, a calibration curve was generated to depict the relationship between absorbance change per second and BDG concentration (abs405 nm vs. pg/ml). Subsequently, this calibration curve was employed to quantify the reactivity of samples in terms of BDG equivalents (pg/ml) (SupFig. 4, 6, 8 and 10).

### Diagnosis of invasive fungal infection by mass spectrometry

As previously described, 1 μL of purified serum was spotted onto a MALDI plate followed by 1 μL of ionic liquid matrix preparation ^34^. Analysis was performed using an AB 4800 MALDI-TOF/TOF analyser (Applied Biosystems/MDS Sciex, Concord, ON, Canada) at fixed laser intensity for 1000 shots/spectrum. Signals were registered between m/z 300 and 800.

### Fractionation of nocardial and mycobacterial cell walls

The *N. nova* patient’s strain was cultivated in Sauton broth and the cell wall was fractionated essentially according to a procedure reported by Besra and coworkers ^20^ from a 1196 mg sample of lyophilised bacteria (Figure 2A). Briefly, the bacterial pellet was rinsed with cold PBS and suspended in buffer PBS containing 2 mM NaCl, 2 mM MgCl_2_, and protease inhibitors (Halt Protease Inhibitor Cocktail w/o EDTA x100, Thermo Scientific) (1.0 g bacteria/mL) before lysis by French press (SLM Aminico French Pressure Cell Press FA-078) at a pressure of 1650 psi. The total lysate was centrifuged at 34,000 g (JA-20 rotor, Beckman) at 4 °C for 30 minutes. The pellet (P1) was washed twice with PBS and freeze-dried. The supernatant (S1) was stored at –20 °C. P1 was suspended in a 2% (v/v) solution of Triton X-100 by vortexing and then centrifuged in a glass tube at 24,000 g (rotor JA-20, Beckman) at 4 °C for 30 minutes to obtain S2 and P2. P2 was washed twice with PBS and freeze-dried and then suspended in 20 ml of 2% SDS in PBS and heated to 95 °C for 1 h with vortexing every 15 minutes. After cooling, the suspension was centrifuged at 24,000 g (JA-20 rotor, Beckman) at 4 °C for 30 minutes to obtain P3 and S3. This procedure was repeated twice. P3 was washed with ultrapure water, then with a mixture of water and acetone (20:80), followed by acetone, before being suspended in 20 mL of a 2% KOH solution in a mixture of toluene and methanol (50:50 vol/vol), and heated at reflux at 70 °C for 16 hours. The solution was then cooled and centrifuged at 24,000 g (JA-20 rotor, Beckman) at 20 °C for 20 minutes to yield P4 and S4. P4 was dried under nitrogen and weighed. Next, 25 ml of 2 M sodium hydroxide were added to P4 and the solution was heated at reflux at 80 °C for 16 hours. After cooling, the solution was neutralised by the dropwise addition of 1 M hydrochloric acid centrifuged at 34,000 g (JA-20 rotor, Beckman) at 20 °C for 30 minutes to yield P5 and S5. Fraction P5 was washed several times with ultrapure water to remove salts and lyophilised. Fraction S5 was filtered (0.45 μm, Millex® syringe filter units, disposable, Durapore® PVDF) and freeze-dried. The arabinogalactan contained in S5 was dissolved in 10 mL of ultrapure water and dialysed (spectra/Por 6 standard regenerated cellulose dialysis membrane, Spectrum Laboratories Inc) against ultrapure water. The obtained arabinogalactan was then lyophilised, weighed and stored at –20 °C. A similar procedure was adapted to purify arabinogalactan from *Mycobacterium bovis* BCG, which served as standard during the study.

### Composition analysis of isolated polysaccharides

Monosaccharides were identified and quantified as reduced and acetylated derivatives (alditol acetates) relative to internal *myo*-inositol standards. Polysaccharide samples (0.2 to 1 mg) were hydrolyzed with 4 M trifluoroacetic acid (TFA) at 110 °C for 3 h and then, after cooling dried. Samples were then placed in water and a few drops of 0.1 M NH_4_OH (pH 9) were added before the samples were reduced with sodium borohydride (NaBH_4_). Excess NaBH_4_ was removed by addition of 10% AcOH in methanol (MeOH), and the solution was dried under a stream of nitrogen. The drying process was repeated twice after addition of 10% AcOH in MeOH (1 mL) and a further two times after addition of MeOH (1 mL). The residue was acetylated with 0.4 mL of acetic anhydride and 0.4 mL of pyridine at 100 °C for 1 h and was then dried under a stream of nitrogen with addition of toluene (1 mL) before being analyzed by GC-MS.

### Linkage analysis of isolated polysaccharides

Methylation was performed using the Ciucanu and Kerek procedure ^35^. The arabinogalactan (0.5 to 1 mg) was dissolved in 1 mL of dimethyl sulfoxide. Freshly powdered NaOH (about 50 mg) was added and the mixture was stirred for 15 min. Methyl iodide (0.2 mL) was then added and the mixture stirred for another 2 h. The reaction was stopped by addition of 3 mL of 10% aqueous sodium thiosulfate (Na_2_S_2_O_3_). The permethylated product was extracted twice with CHCl_3_ (2 mL). The organic phase was washed five times with water (4 mL), filtered through a cotton-plugged Pasteur pipette and the filtrate was evaporated. The product was hydrolyzed with 4 M TFA (110 °C, 3 h), cooled to room temperature, dried, reduced with sodium borodeuteride (NaBD_4_), and then peractylated by incubation in acetic anhydride before being analyzed by GC-MS (see below). Methylated derivatives were identified using the Complex Carbohydrate Research Center partially methylated alditol acetates database (www.ccrc.uga.edu/specdb/ms/pmaa/pframe.html) and by comparison with authentic standards of methylation analysis of arabinogalactan from mycobacteria ^36^.

### Gas chromatography of methylated alditol acetates

Gas chromatography (GC) was performed on a Trace GC Ultra system (Thermo Scientific) equipped with a NMTR-5MS capillary column (30 m x 0.25 mm) and a flame ionization detection (FID) unit using a temperature gradient from 170 °C to 250 °C at 5 °C·min^−1^. GC-MS was performed using a Trace GC Ultra system TSQ quantum GC detector (Thermo Scientific), equipped with a SILGEL1MS capillary column (30 m x 0.25 mm), and a temperature gradient from 170 °C to 230 °C at 3 °C min^−1^ then to 270 °C at 10 °C min^−1^.

### Nuclear Magnetic Resonance spectroscopy of isolated polysaccharides

Samples were solubilized in highly enriched deuterated water (D_2_O, 99.96% deuterium; EurisoTop, St-Aubin, France) and lyophilized. This process was repeated twice. Data were recorded on a 9.4-T spectrometer and an 18.8-T spectrometer (Infrastruture de Recherche-Très Hauts Champs-Résonance Magnétique Nucléaire, CNRS); at these field strengths, ^1^H resonate at 400 and 800 MHz, and ^13^C resonate at 100 and 201 MHz, respectively. As an internal standard, 1 μl of a solution of 2.5 μl of acetone in 10 mL of D_2_O was added to each sample. All pulse sequences were taken from the Bruker library of pulse programs and then optimized for each sample. Spectral widths were 12 and 200 ppm for the ^1^H and ^13^C observations, respectively. TOCSY was performed with various mixing times of 40 to 120 ms, and spectra were recorded with a mixing time of 300 ms. Edited ^1^H–^13^C HSQC and HMBC spectra were recorded with 1,536 data points for detection and 256 data points for indirect direction.

### Macrophage infection experiments

The THP-1 human pro-monocytic cell line (European Collection of Authenticated Cell Cultures, ECACC no. 88081201) was cultured at a density of 3 × 10^5 cells/ml in RPMI-1640 medium (Roswell Park Memorial Institute Medium, Gibco), supplemented with 10% (v/v) decomplemented fetal calf serum (Dutcher), 2 mM L-glutamine and 20 μM β-mercaptoethanol. Cultures were kept under a humidified atmosphere containing 5% CO2 at 37° C. To induce differentiation into a macrophage phenotype, THP-1 cells were treated with 20 nM phorbol-12-myristate-13-acetate (Sigma-Aldrich) for 72 hours in sterile 24-well culture plates (*Nunclon Delta surface*, Thermoscientific). PMA-activated THP-1 macrophages were then infected with two different concentrations of *Nocardia nova* (Multiplicity of Infection, MOI, 5:1 and 15:1) in RPMI-1640 medium devoid of serum for 6 hours. The supernatants were then discarded, and the macrophages were cultured for an additional 12 hours in serum-free RPMI-1640 medium supplemented with 20 μg/ml amikacin (amikacin sulfate, Sigma). Following incubation, the media were collected by centrifugation and subjected to Fungitell® assay at a 1/1000 dilution. Control experiments included incubation of either bacteria alone or uninfected macrophages under identical conditions.

### Synthesis of NnAG1–NnAG6

General methods, detailed procedures and spectroscopic data can be found in the Online Methods section.

## ACKNOWLEDGMENTS

This work was supported by grants from INSERM and intramural funding from Academia Sinica (TLL). We are grateful to Nadine François and Rachid Aiidjou for their support with the experimentation and to Fanny Vuotto for insightful advice. We thank Michael Howsam for helpful advice about this work and editorial writing assistance. We are grateful to the PAGes core facilty (Plateformes Lilloises en Biologie et Santé (PLBS) - UAR 2014 - US 41) for providing the scientific and technical environment conducive to achieving this work. We acknowledge the Academia Sinica High-Field NMR Center (HFNMRC) and the Medicinal Chemistry and Analytical Core Facility, funded by Academia Sinica Core Facility and Innovative Instrument Project (AS-NBRPCF-111-201 and AS-CFII-108-112), for technical support in acquiring NMR data. Optical rotation data was obtained in the Biophysical Instrumentation Laboratory at the Academia Sinica Institute of Biological Chemistry. Mass spectrometry analyses were performed by Mass Spectrometry Facility of the Academia Sinica Institute of Chemistry, Academia Sinica, Taiwan.

## AUTHOR CONTRIBUTIONS

Research idea and study design: MT and MF; clinical information provided by: MU, EB, AL, KF, PR, MH and MF; clinical data acquisition: MF; microbiological data analysis: MT, SL, BS, FW and DP; performed research: MT, PL, YG and CM; design and synthesis of oligosaccharides **NnAG1**– **NnAG6**: WTC and TLL. All authors discussed the data and approved the submission.

## Extended data

**SupFig 1.**
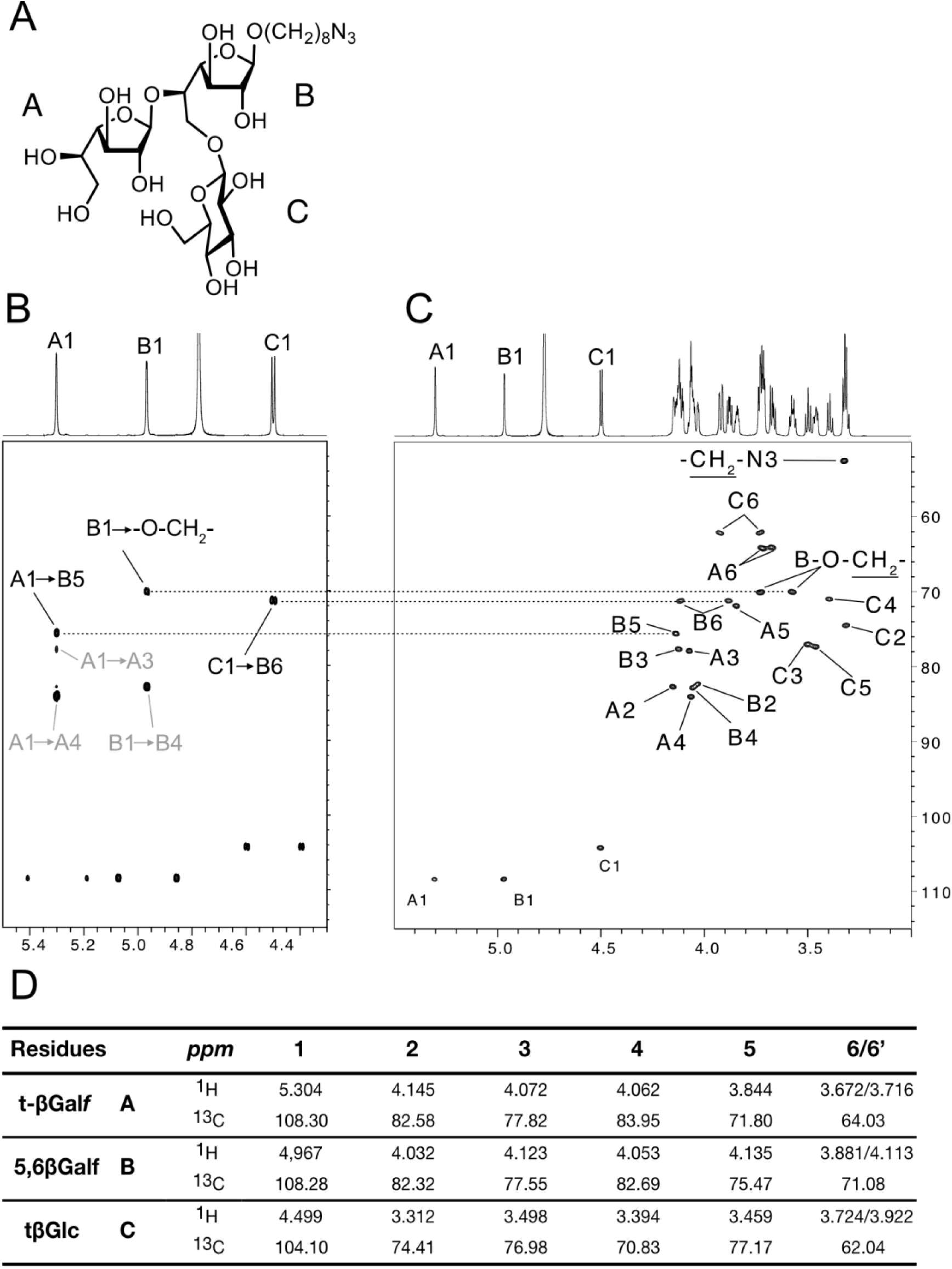
Characterisation of βGalf-5(βGlcp-6)βGalf epitope from synthetic NnAG1. (A) structure of the trisaccharide, (B) ^1^H–^13^C HMBC and (C) ^1^H–^13^C HSQC spectra showing the substitution of 5,6βGalf residue by a tβGlc residue in C6 position. (D) Full spin system of the trisaccharide deduced from ^1^H–^1^H COSY, ^1^H–^1^H TOCSY, ^1^H–^13^C HSQC and ^1^H–^13^C HSQC–TOCSY spectra. In (B), black labels represent inter-residues connections that demonstrates the sequence of the trisaccharide as represented in (A). Grey labels represent intraresidues connections.

**SupFig 2.**
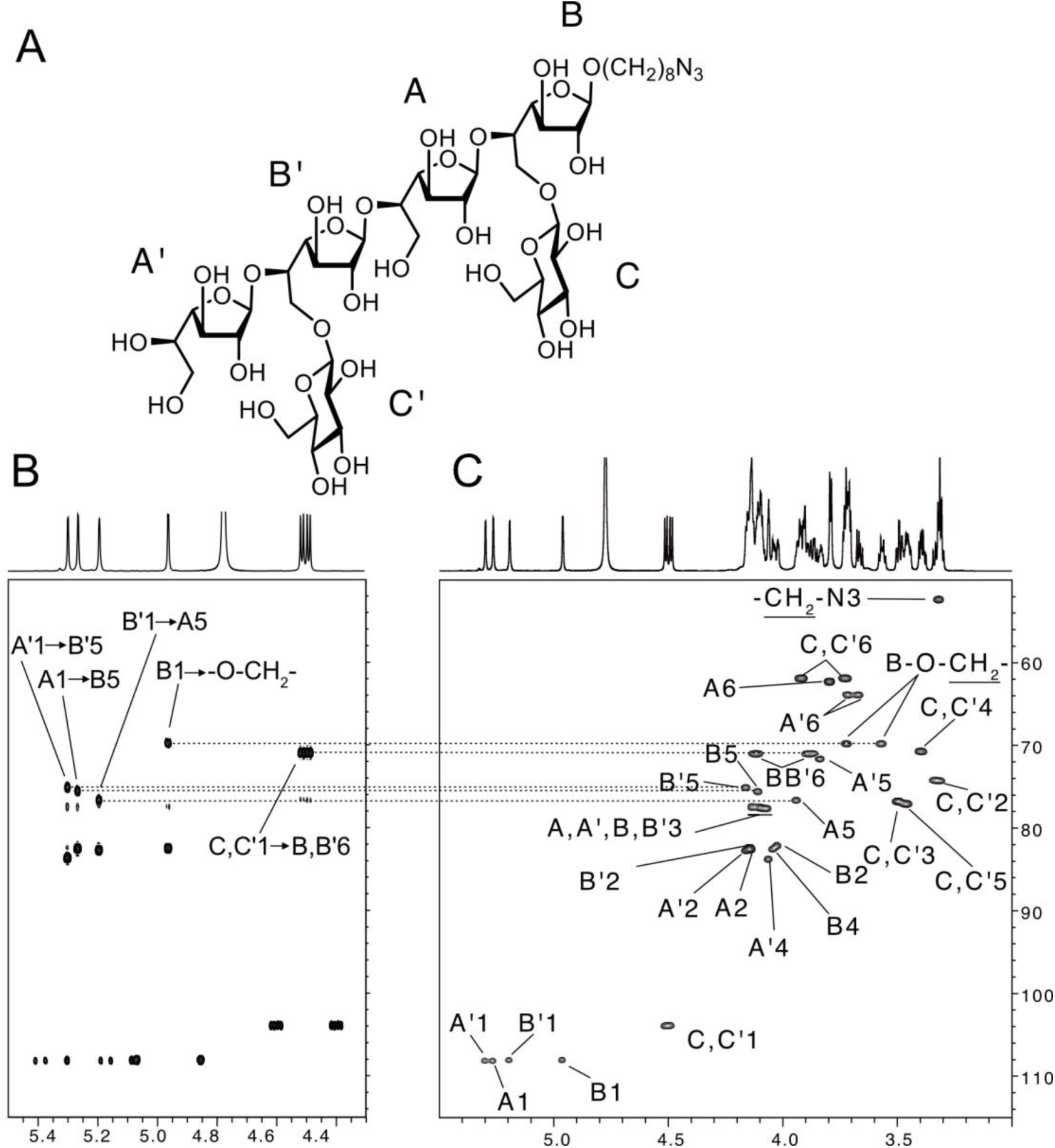
Characterisation of βGalf-5(βGlcp-6)βGalf internal and external epitopes from synthetic NnAG2. (A) Structure of the hexasaccharide, (B) ^1^H–^13^C HMBC and (C) ^1^H–^13^C HSQC spectra showing the substitution of 5,6βGalf residues B and B’ by tβGlc residues C and C’ in C6 position.

**SupTable 1.**
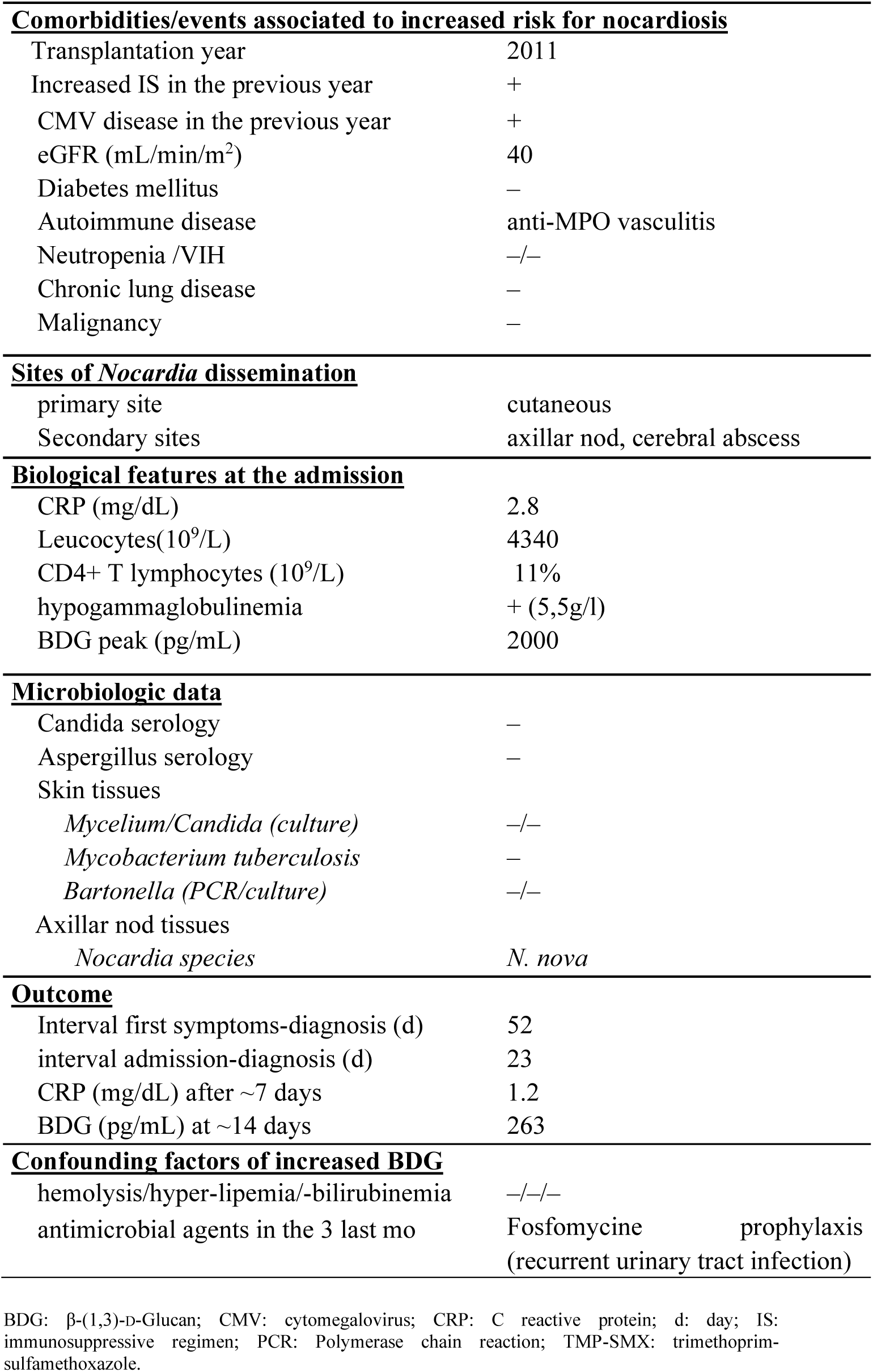
Patient characteristics.

**SupFig 3.**
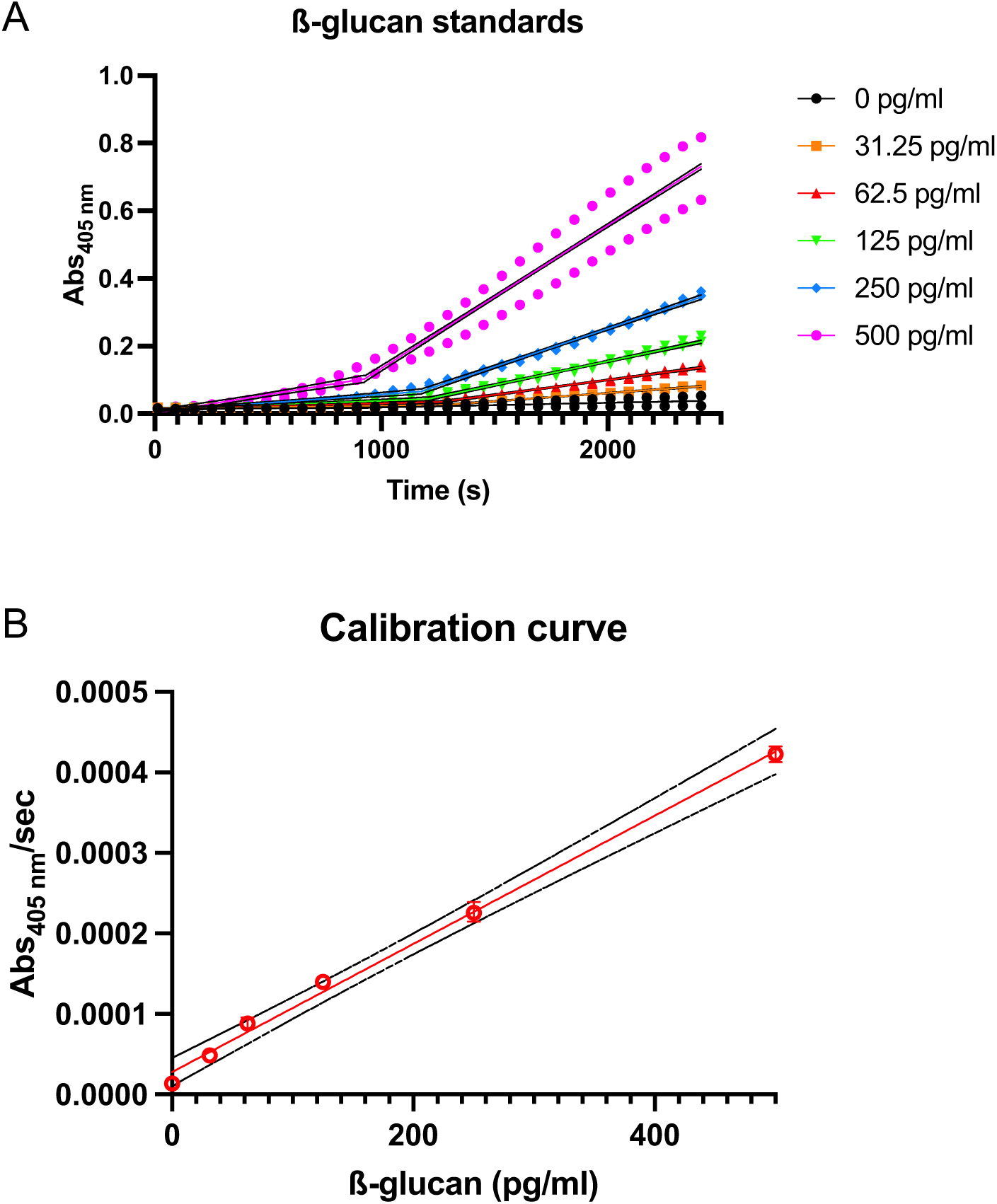
Development of standard curve for Fungitell assay for samples shown in Fig.2B. (A) kinetic data obtained with BDG standards were extracted to determine the slope of absorbance at 405 nm over time, (B) calibration curve obtained used to quantify the reactivity of samples in terms of BDG equivalents (pg/ml).

**SupFig 4.**
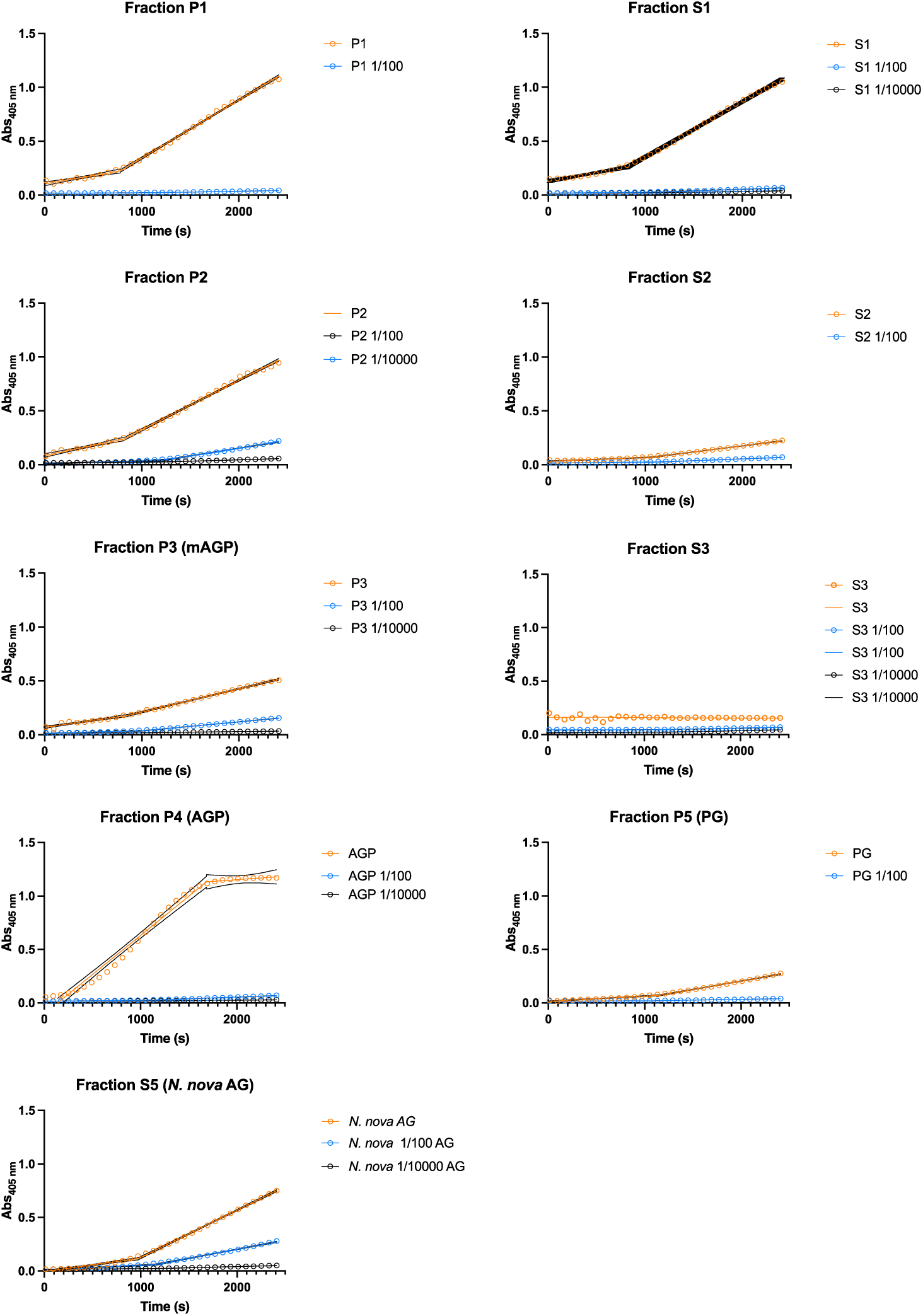
Fungitell assay for non-BDG samples (samples for Fig.2B). The slope of absorbance change at 405 nm over time is used with the calibration curve (SupFig. 3B) to obtain the reactivity of the sample in BDG equivalent.

**SupFig 5.**
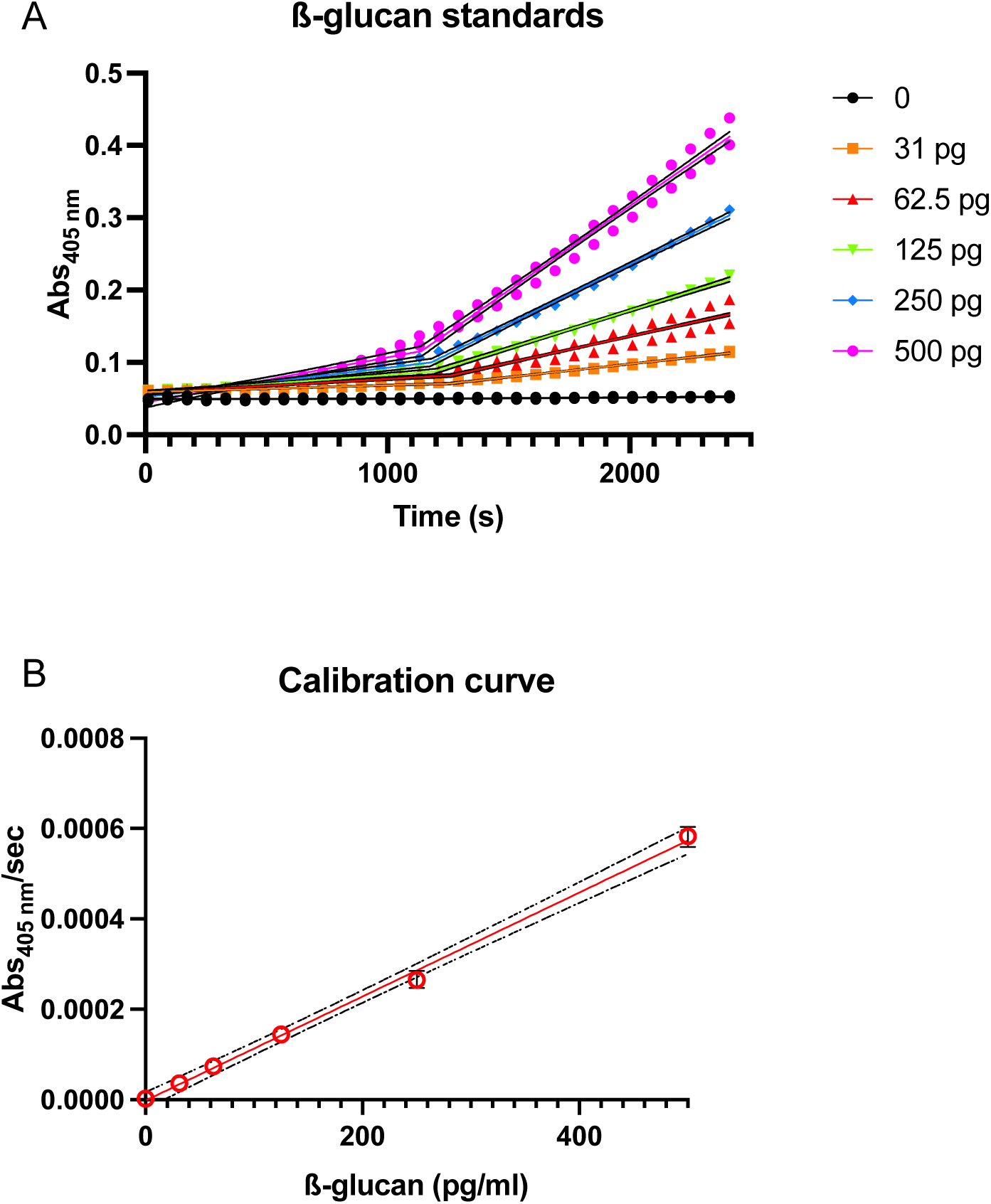
Development of standard curve for Fungitell assay for samples in Fig.2C. (A) kinetic data obtained with BDG standards were extracted to determine the slope of absorbance at 405 nm over time, (B) calibration curve obtained used to quantify the reactivity of samples in terms of BDG equivalents (pg/ml).

**SupFig 6.**
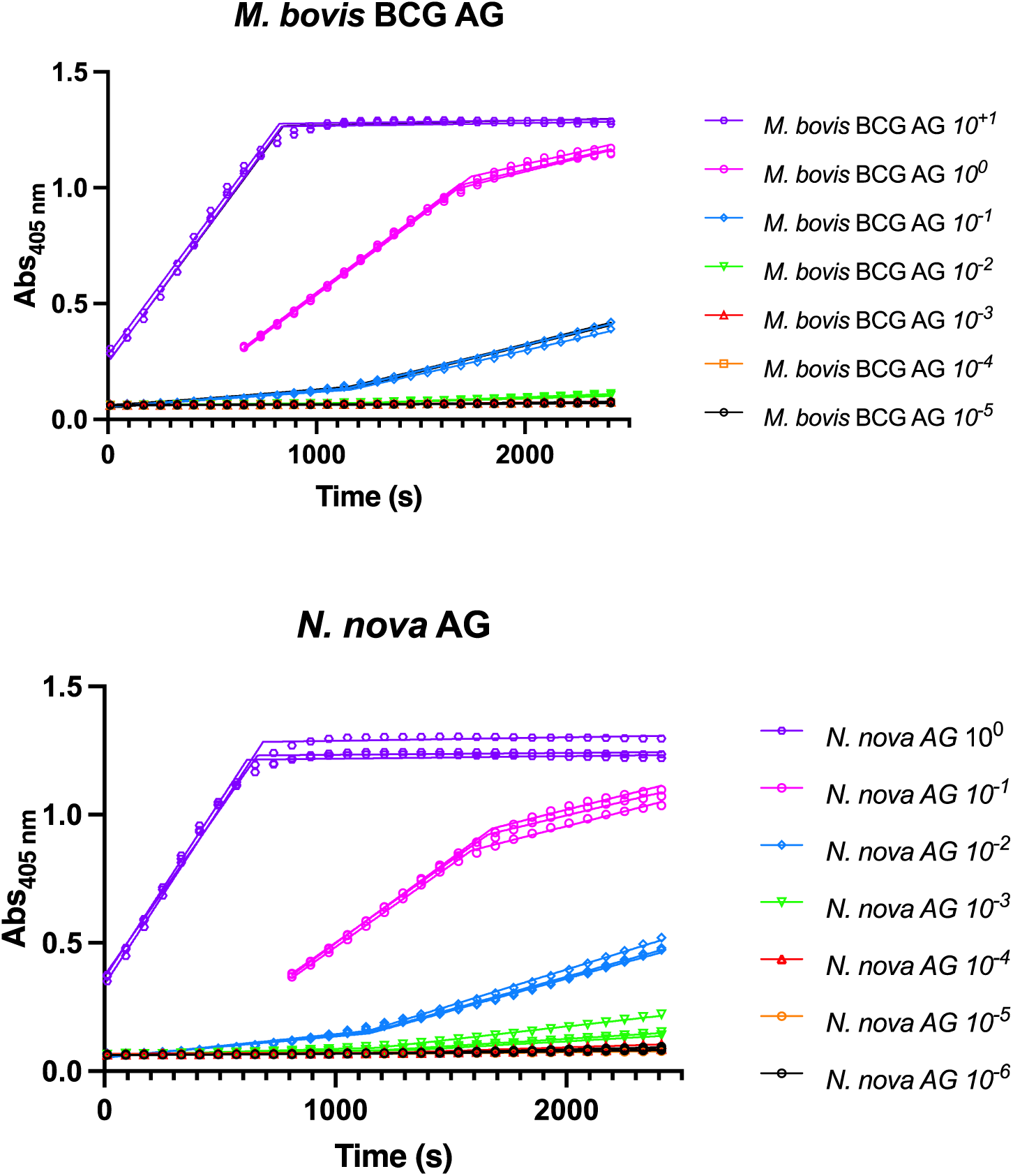
Fungitell assay for non-BDG samples (samples for Fig.2C). The slope of absorbance change at 405 nm over time is used with the calibration curve (SupFig. 5B) to obtain the reactivity of the sample in BDG equivalent.

**SupFig 7.**
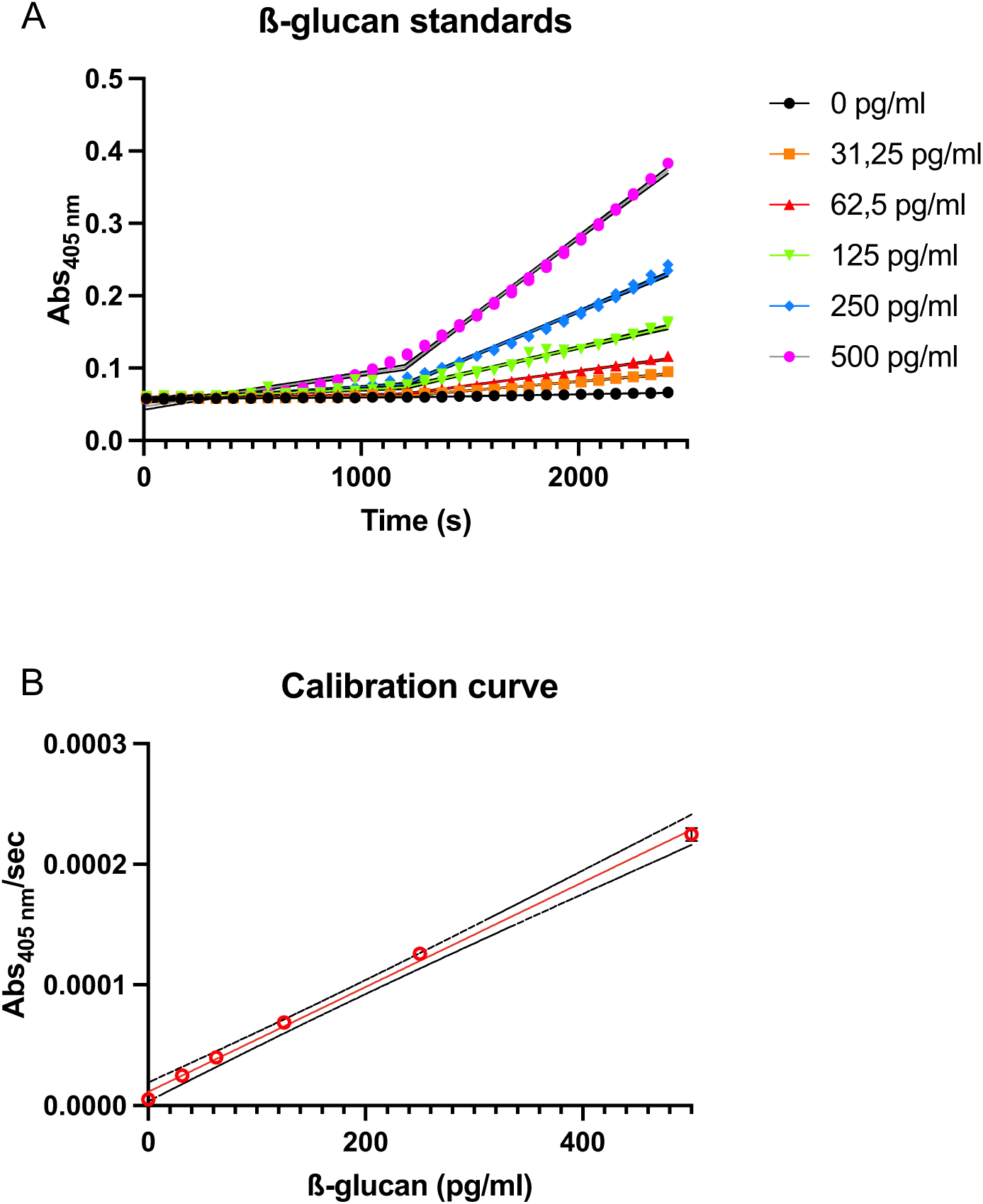
Development of standard curve for Fungitell assay for samples in Fig. 6A. (A) kinetic data obtained with BDG standards were extracted to determine the slope of absorbance at 405 nm over time, (B) calibration curve obtained used to quantify the reactivity of samples in terms of BDG equivalents (pg/ml).

**SupFig 8.**
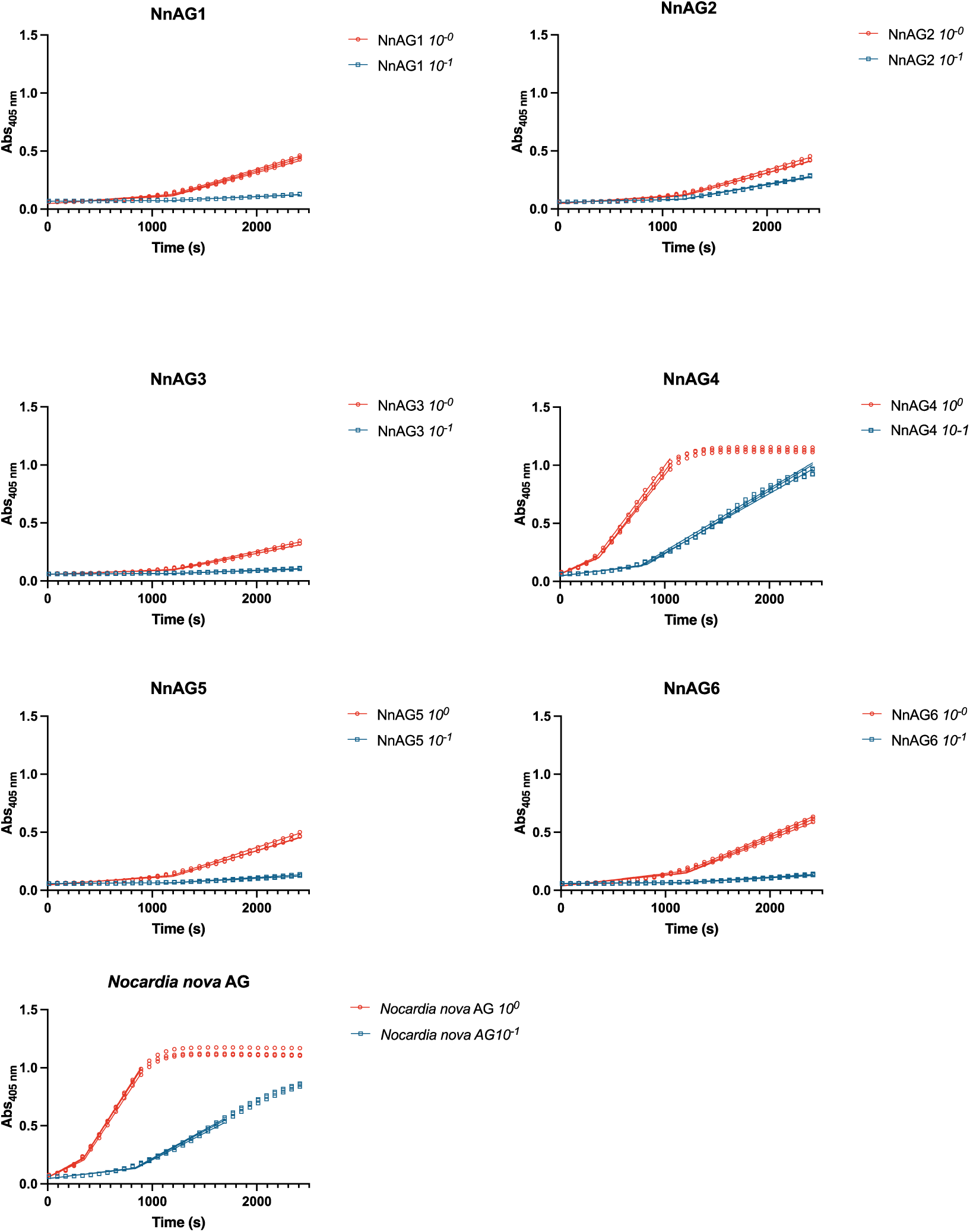
Fungitell assay for non-BDG samples (samples for Fig.6A). The slope of absorbance change at 405 nm over time is used with the calibration curve (SupFig. 7B) to obtain the reactivity of the sample in BDG equivalent.

**SupFig 9.**
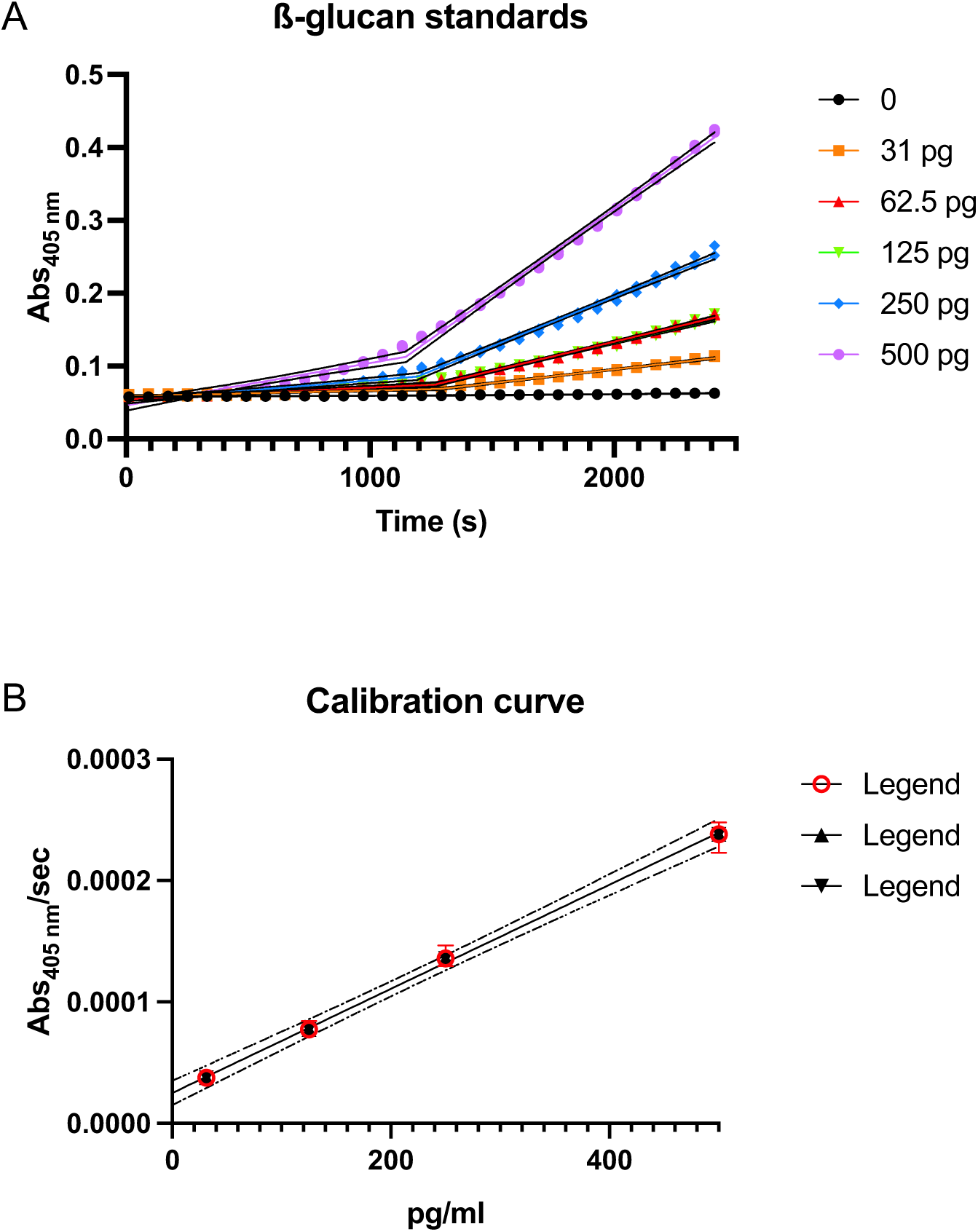
Development of standard curve for Fungitell assay for samples in Fig. 6B. (A) kinetic data obtained with BDG standards were extracted to determine the slope of absorbance at 405 nm over time, (B) calibration curve obtained used to quantify the reactivity of samples in terms of BDG equivalents (pg/ml).

**SupFig 10.**
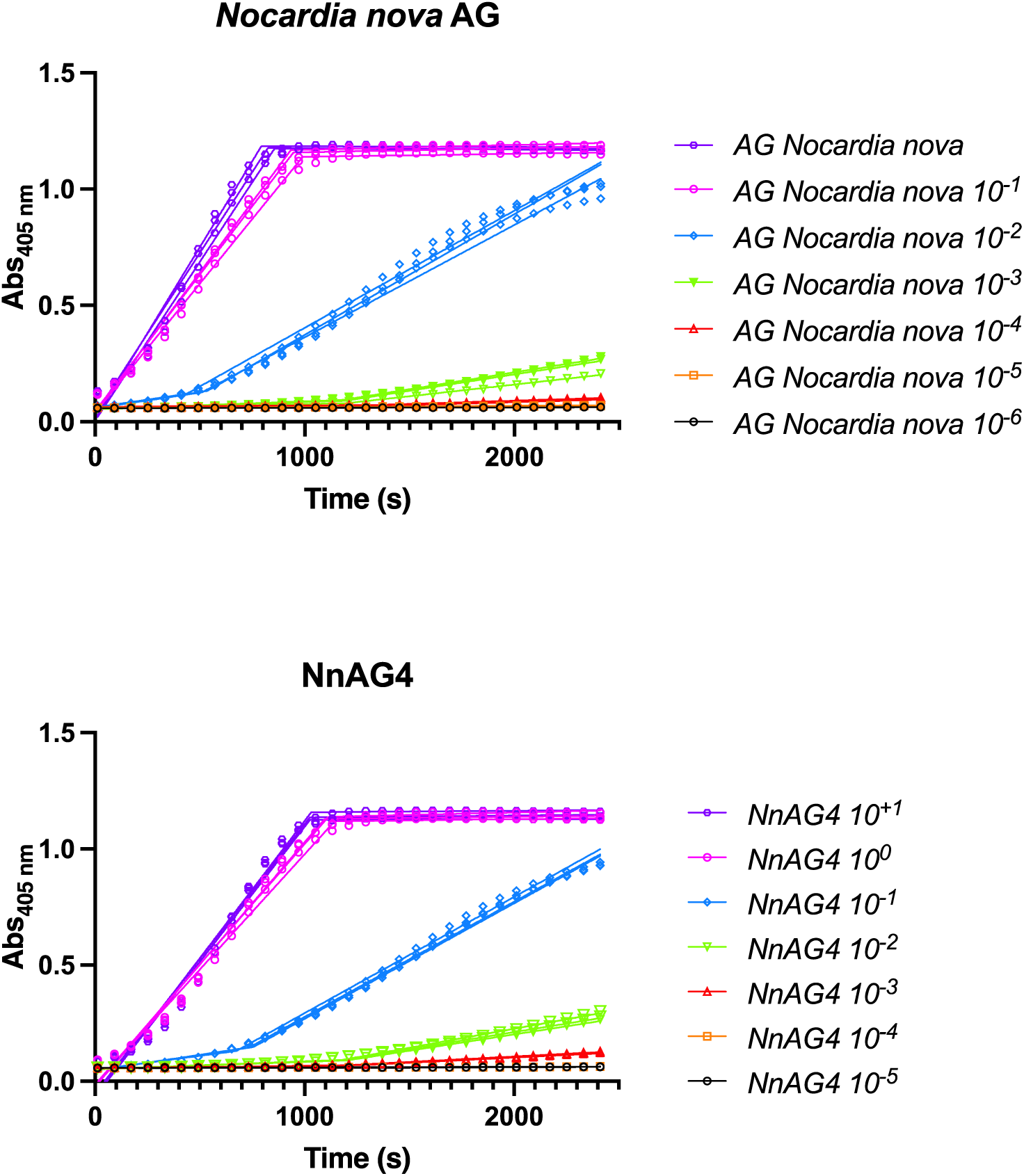
Fungitell assay for non-BDG samples (samples for. **Fig.6B**). The slope of absorbance change at 405 nm over time is used with the calibration curve (SupFig. 9B) to obtain the reactivity of the sample in BDG equivalent.

## Online methods

### Detailed procedures and data for the synthesis of NnAG1–NnAG6

#### General Methods

Reactions were carried out in oven-dried glassware under an atmosphere of dry argon. All reagents used were purchased from commercial sources and were used without further purification unless noted. Oven-dried (135 °C) 4 Å molecular sieves were further activated by flame-drying under vacuum (0.5 mmHg) just prior to use. Dowex® 50W ion-exchange resin were washed by methanol for preparation. Solvents used in reactions were purified by successive passage through columns of alumina and copper under argon. Unless stated otherwise, all reactions were carried out at room temperature (rt) under a positive pressure of argon and were monitored by TLC on silica gel 60 F_254_. Spots were detected under UV light or by charring with cerium molybdate stain or *p*-anisaldehyde stain. Unless otherwise indicated, normal-phase column chromatography was performed on silica gel 60 (40–60 µm). The ratio between silica gel and crude product ranged from 150 to 50:1 (w/w). Size exclusion column chromatography was performed on HW-20. Reversed-phase column chromatography was performed on octadecyl silica (40–63 μm). In the processing of reaction mixtures, solutions of organic solvents were washed with equal volumes of aqueous solutions. Organic solutions were concentrated under vacuum at < 40°C (bath). ^1^H NMR spectra were recorded at 500 or 600 MHz on a Bruker Avance 500 or 600 spectrometer, and chemical shifts are reported in ppm and referenced to either TMS (0.0 ppm, CDCl_3_) or HOD (4.78 ppm, D_2_O). ^13^C NMR spectra were recorded at 125 or 151 MHz, and ^13^C chemical shifts are reported in ppm referenced to internal CDCl_3_ (77.23 ppm, CDCl_3_) or external 2,2-dimethyl-2-silapentane-5-sulfonate sodium salt (DSS, –0.7 ppm, D_2_O). ^19^F NMR spectra were recorded at 470 or 564 MHz on a Bruker Avance 500 or 600 spectrometer. ^13^F chemical shifts are reported in ppm referenced to external CFCl_3_ (0 ppm, CDCl_3_). ^13^C NMR peak multiplicities, where reported, were inferred using either DEPT 135 or edited HSQC experiments. Where ^1^H and ^13^C NMR peak assignments are given, these were made unambiguously by a combination of ^1^H/^1^H COSY, ^1^H/^13^C HSQC, and ^1^H/^13^C HMBC experiments. Melting points were obtained using a Buchi melting point M-560 apparatus. Optical rotations were measured using Model 341 Polarimeter (PerkinElmer) at 22 ± 2 °C at the S4 sodium D line (589 nm) and are in units of deg·mL(dm·g)^ࢤ1^. High resolution mass spectra were measured by Mass Spectrometry facility of the Institute of Chemistry, Academia Sinica.

#### 8-Azidooctyl β-D-galactofuranosyl-(1→5)-[β-D-glucopyranosyl-(1→6)]-β-D-galactofuranoside (NnAG1)

To a solution of trisaccharide **13** (15 mg, 9.5 μmol) in 3:1 CH_3_OH–CH_2_Cl_2_ (2 mL) was added sodium methoxide in CH_3_OH (0.1 M) dropwise until the pH of the reaction mixture was 12. After the solution was stirred at room temperature overnight, acid washed Dowex® 50W ion-exchange resin was added to adjust the pH of the mixture from alkaline to ∼7. The resin was then removed by filtration and the filtrate was concentrated. The resulting residue was purified by reversed phase chromatography (18 mL gel, gradient from 10% CH_3_OH in H_2_O to 60% CH_3_OH in H_2_O). The product was evaporated to dryness, re-dissolved with H_2_O and lyophilized to afford **NnAG1 (**4 mg, 90%) as a white foam. *R*_f_ 0.55 (*n*-propanol–H_2_O–AcOH, 10:2:1); [α]_D_ –101.0 (*c* 0.5, H_2_O); ^1^H NMR (500 MHz, D_2_O, δ_H_) 5.30 (d, *J* = 1.9 Hz, 1H, H-1’), 4.97 (d, *J* = 2.4 Hz, 1H, H-1), 4.50 (d, *J* = 7.9 Hz, 1H, H-1’’), 4.15–4.10 (m, 4H, H-3, H-2’, H-3’’, H-4’’), 4.09–4.01 (m, 4H, H-2, H-4, H-4’, H-2’’), 3.95–3.82 (m, 3H, H-5’, H-5’’, H-6b’’), 3.75–3.70 (m, 3H, H-6b, H-6b’, H-6a’’), 3.66 (dd, *J* = 11.7, 7.3 Hz, 1H, H-6a’), 3.57 (dt, *J* = 9.9, 6.5 Hz, 1H, H-6a), 3.52–3.43 (m, 2H, OC<u>H_2_</u>CH_2_, H-5), 3.39 (t, *J* = 9.4 Hz, 1H, OC<u>H_2_</u>CH_2_), 3.34–3.28 (m, 3H, H-3’, C<u>H_2_</u>N_3_), 1.60 (q, *J* = 7.0 Hz, 4H, octyl CH_2_), 1.42–1.30 (m, 8H, octyl CH_2_); ^13^C NMR (126 MHz, D_2_O, δ_C_) 107.0 (C-1, C-1’), 102.7 (C-1’’), 82.6 (C-4), 81.4 (C-4’), 81.3 (C-2), 80.9 (C-2’), 76.5 (C-5), 76.3 (C-3’’), 75.9 (C-2’’), 75.6 (C-5’’), 74.2 (C-3), 73.1 (C-3’), 70.5 (C-5’), 69.8 (C-4’’), 69.6 (C-6), 68.7 (O<u>C</u>H_2_CH_2_), 62.8 (C-6’), 60.7 (C-6’’), 51.2 (<u>C</u>H_2_N_3_), 28.5 (octyl CH_2_), 28.2 (octyl CH_2_), 28.1 (octyl CH_2_), 27.9 (octyl CH_2_), 25.8 (octyl CH_2_), 25.0 (octyl CH_2_); HRMS (ESI) Calcd for (M + Na) C_26_H_47_N_3_O_16_Na: 680.2849. Found 680.2848.

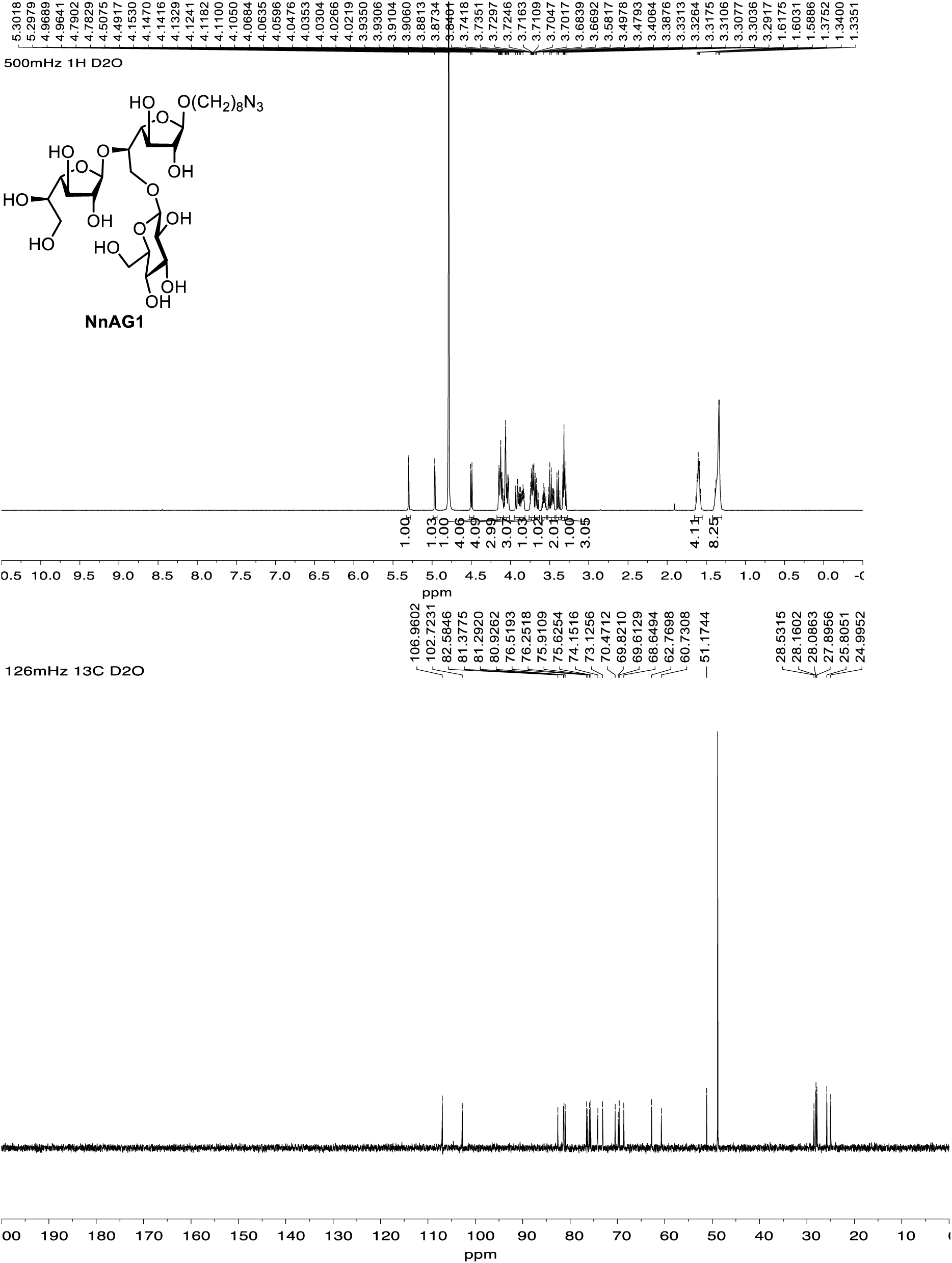

#### 8-Azidooctyl β-D-galactofuranosyl-(1→5)-[β-D-glucopyranosyl-(1→6)]-β-D-galactofuranosyl-(1→5)-β-D-galactofuranosyl-(1→5)-[β-D-glucopyranosyl-(1→6)]-β-D-galactofuranoside (NnAG2)

Using the procedure described for the synthesis of **NnAG1**, hexasaccharide **15** (35 mg, 11.2 μmol) in 3:1 CH_3_OH–CH_2_Cl_2_ (2 mL) was converted to **NnAG2** (7 mg, 90%), which was obtained as a white foam. *R*_f_ 0.43 (*n*-propanol–H_2_O–AcOH, 10:2:1); [α]_D_ –95.0 (*c* 1.4, H_2_O); ^1^H NMR (600 MHz, D_2_O, δ_H_) 5.29 (d, *J* = 2.0 Hz, 1H), 5.26 (d, *J* = 1.8 Hz, 1H), 5.19 (d, *J* = 1.7 Hz, 1H), 4.96 (d, *J* = 2.3 Hz, 1H), 4.52–4.48 (m, 2H), 4.18–4.00 (m, 16H), 3.96–3.81 (m, 6H), 3.79–3.78 (m, 2H), 3.75–3.69 (m, 4H), 3.66 (dd, *J* = 11.7, 7.3 Hz, 1H), 3.56 (dt, *J* = 10.0, 6.5 Hz, 1H), 3.52–3.43 (m, 4H), 3.40–3.36 (m, 2H), 3.35–3.28 (m, 4H), 1.63–1.55 (m, 4H), 1.40–1.30 (m, 8H); ^13^C NMR (151 MHz, D_2_O, δ_C_) 107.02 (x 2), 106.98, 106.93, 102.8, 102.7, 82.6, 81.54, 81.47, 81.4, 81.3, 80.9, 76.5, 76.30, 76.25, 75.9, 75.6, 75.44, 74.40, 73.9, 73.1, 73.0, 70.4, 69.9, 69.8, 69.58, 69.55, 68.6, 62.7, 61.1, 60.71, 60.67, 51.2, 28.5, 28.2, 28.1, 27.9, 25.8, 25.0; HRMS (ESI) Calcd for (M + Na) C_44_H_77_N_3_O_31_Na: 1166.4433. Found 1166.4423.

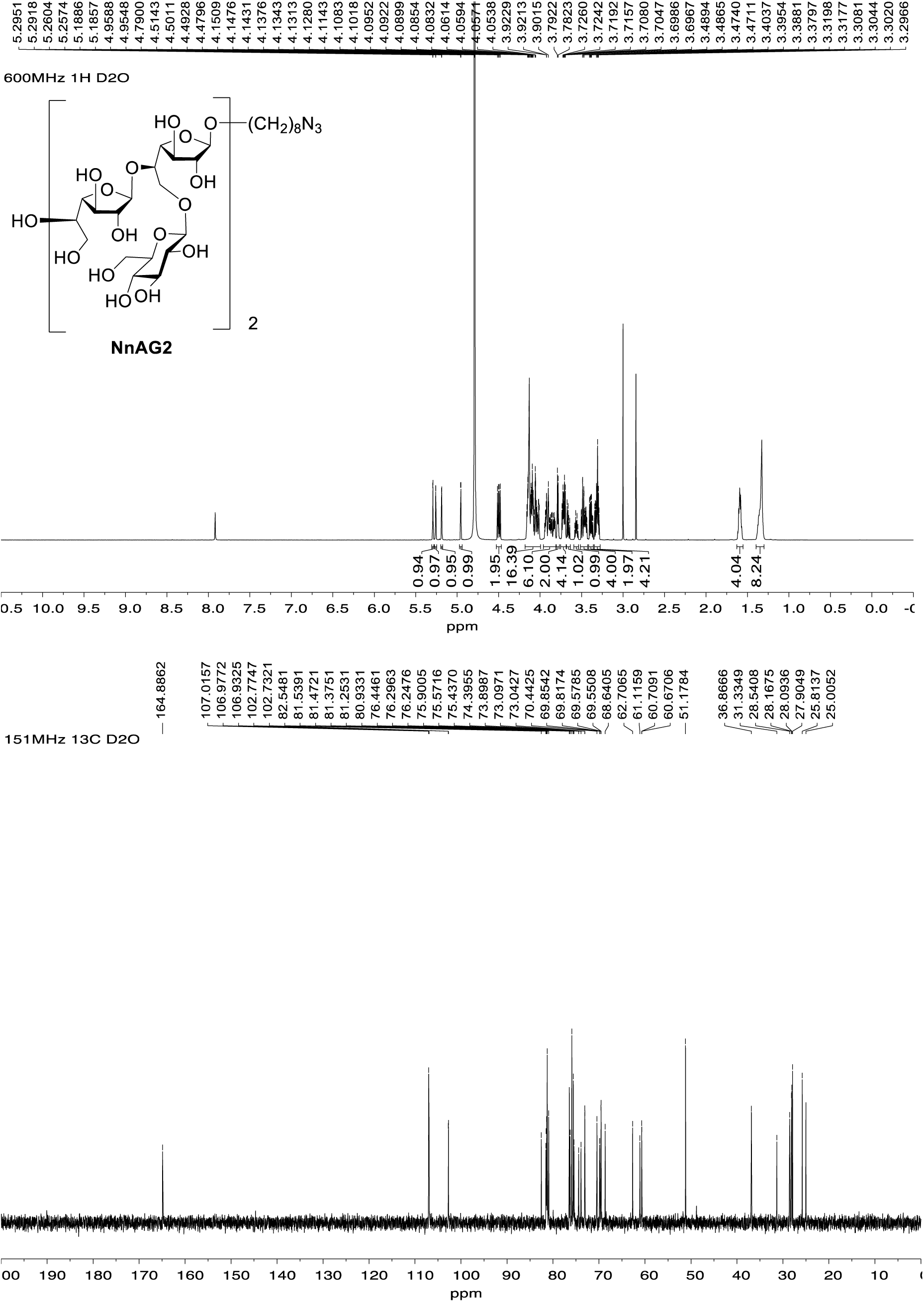

#### 8-Azidooctyl β-D-galactofuranosyl-(1→5)-[β-D-glucopyranosyl-(1→6)]-β-D-galactofuranosyl-(1→5)-β-D-galactofuranosyl-(1→5)-[β-D-glucopyranosyl-(1→6)]-β-D-galactofuranosyl-(1→5)-β-D-galactofuranosyl-(1→5)-[β-D-glucopyranosyl-(1→6)]-β-D-galactofuranoside (NaAG3)

Using the procedure described for the synthesis of **NnAG1**, nonasaccharide **17** (24 mg, 5.2 μmol) in 3:1 CH_3_OH–CH_2_Cl_2_ (0.6 mL) was converted to **NaAG3** (7 mg, 92%), which was obtained as a white foam. *R*_f_ 0.18 (*n*-propanol–H_2_O–AcOH, 10:2:1); [α]_D_ –71.4 (*c* 0.7, H_2_O); ^1^H NMR (500 MHz, D_2_O, δ_H_) 5.21 (d, *J* = 2.0 Hz, 1H), 5.18–5.17 (m, 2H), 5.10–5.09 (m, 2H), 4.87 (d, *J* = 2.4 Hz, 1H), 4.45–4.37 (m, 3H), 4.11–3.90 (m, 24H), 3.88–3.67 (m, 13H), 3.65–3.61 (m, 5H), 3.57 (dd, *J* = 11.7, 7.3 Hz, 1H), 3.48 (dt, *J* = 10.0, 6.5 Hz, 1H), 3.44–3.34 (m, 6H), 3.34–3.27 (m, 3H), 3.25–3.19 (m, 5H), 1.54–1.48 (m, 4H), 1.32–1.20 (m, 8H); ^13^C NMR (126 MHz, D_2_O, δ_C_) 107.02 (x 3), 106.97 (x 3), 102.8, 102.7 (x 2), 82.6, 81.53, 81.50, 81.47, 81.44, 81.39, 81.36, 81.34, 81.25, 80.9, 76.44, 76.38, 76.3, 76.2, 75.9, 75.6, 75.4, 75.3, 74.4, 74.0, 73.9, 73.1, 73.0, 70.4, 69.84, 69.81, 69.57, 69.55, 68.6, 62.7, 61.11, 61.06, 60.70, 60.67, 51.2, 28.5, 28.2, 28.1, 27.9, 25.8, 25.0; HRMS (ESI) Calcd for (M + Na) C_62_H_107_N_3_O_46_Na: 1652.6018. Found 1652.6015.

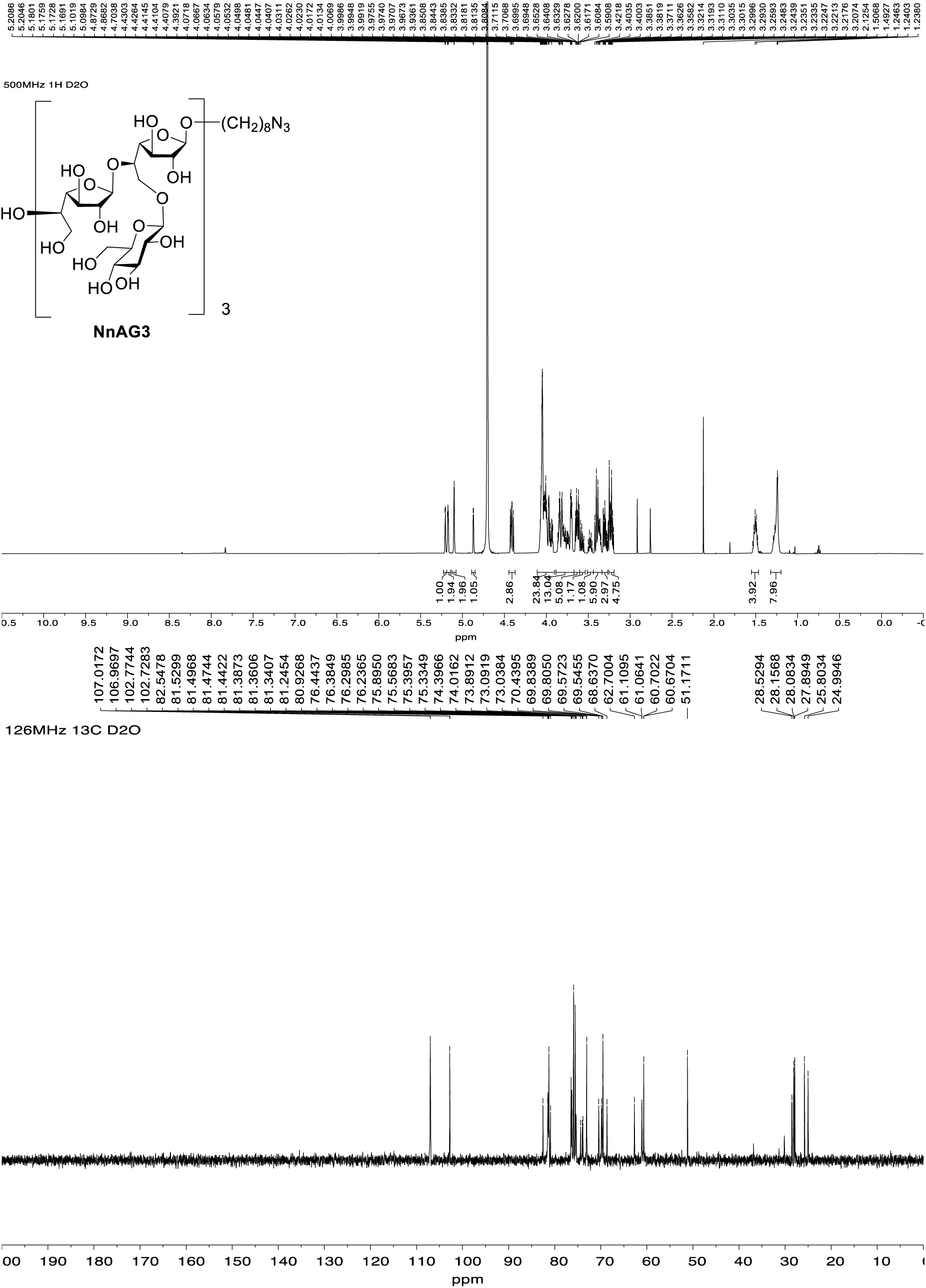

#### 8-azidooctyl β-D-galactofuranosyl-(1→5)-[β-D-glucopyranosyl-(1→6)]-β-D-galactofuranosyl-(1→5)-β-D-galactofuranosyl-(1→5)-[β-D-glucopyranosyl-(1→6)]-β-D-galactofuranosyl-(1→5)-β-D-galactofuranosyl-(1→5)-[β-D-glucopyranosyl-(1→6)]-β-D-galactofuranosyl-(1→5)-β-D-galactofuranosyl-(1→5)-[β-D-glucopyranosyl-(1→6)]-β-D-galactofuranoside (NnAG4)

Using the procedure described for the synthesis of **NnAG1**, dodecsaccharide **19** (21 mg, 3.5 μmol) in 3:1 CH_3_OH– CH_2_Cl_2_ (1 mL) was converted to **NnAG4** (5 mg, 72%), which was obtained as a white foam. *R*_f_ 0.25 (*n*-propanol–H_2_O–AcOH, 6:2:1); [α]_D_ –42.0 (*c* 0.5, H_2_O); ^1^H NMR (500 MHz, D_2_O, δ_H_) 5.21 (d, *J* = 2.0 Hz, 1H), 5.18–5.17 (m, 3H), 5.10–5.09 (m, 3H), 4.87 (d, *J* = 2.4 Hz, 1H), 4.46–4.37 (m, 4H), 4.14–3.90 (m, 32H), 3.89–3.68 (m, 18H), 3.68–3.53 (m, 7H), 3.53–3.15 (m, 19H), 1.53–1.48 (m, 4H), 1.34–1.20 (m, 8H); ^13^C NMR (126 MHz, D_2_O, δ_C_) 107.02 (x 4), 106.95 (x 3), 106.9, 102.8 (x 3), 102.7, 81.52, 81.47, 81.44, 81.37, 81.28, 81.25, 76.44, 76.38, 76.3, 76.2, 75.9, 75.6, 73.1, 73.0, 69.84, 69.81, 69.57, 69.55, 62.7, 61.12, 61.07, 60.7, 51.2, 28.5, 28.2, 28.1, 27.9, 25.8, 25.0; HRMS (ESI) Calcd for (M + Na) C_80_H_137_N_3_O_61_Na: 2138.76026. Found 2138.75970.

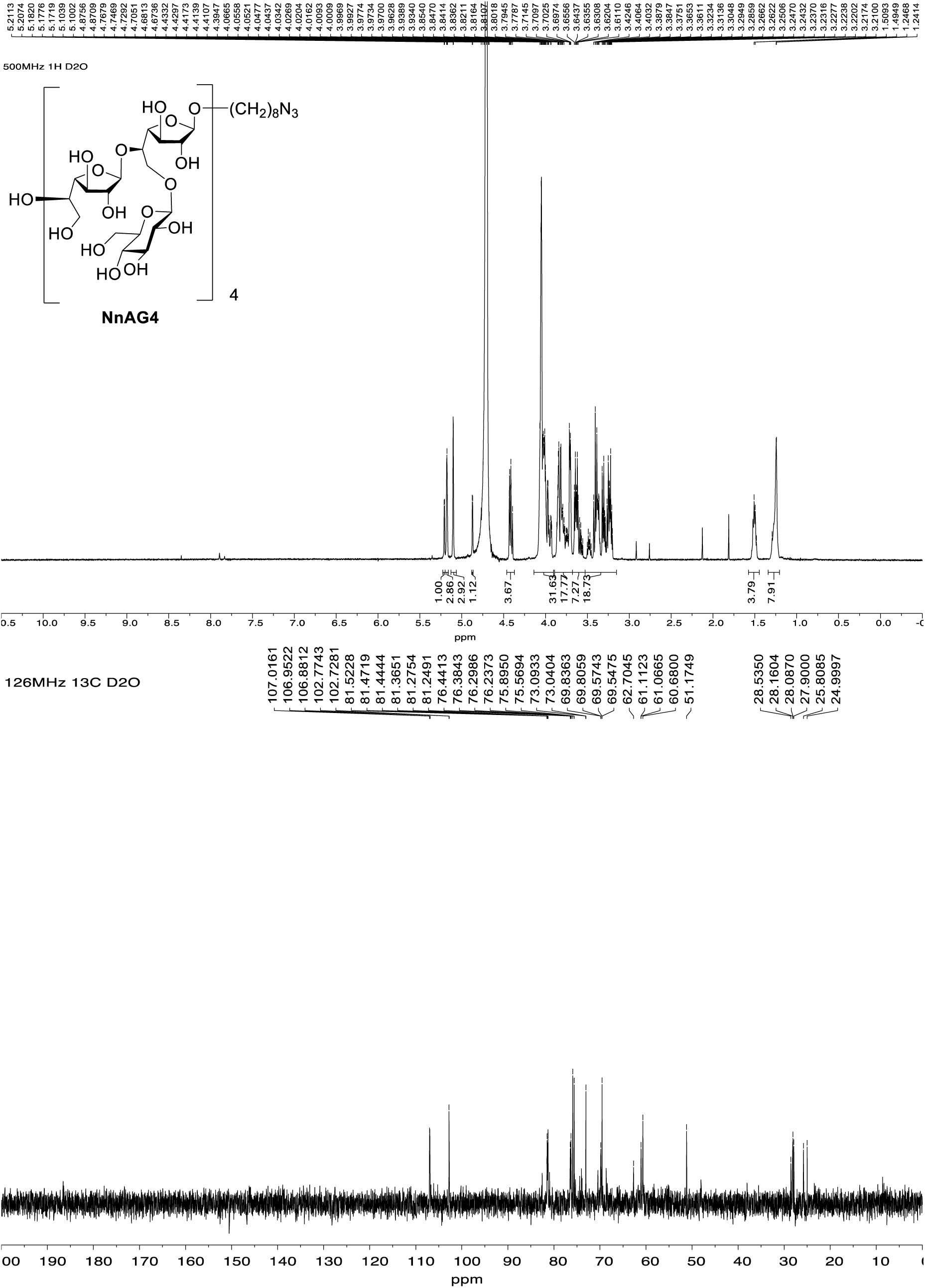

#### 8-Azidooctyl β-D-galactofuranosyl-(1→5)-[β-D-glucopyranosyl-(1→6)]-β-D-galactofuranosyl-(1→5)-β-D-galactofuranosyl-(1→5)-[β-D-glucopyranosyl-(1→6)]-β-D-galactofuranosyl-(1→5)-β-D-galactofuranosyl-(1→5)-[β-D-glucopyranosyl-(1→6)]-β-D-galactofuranosyl-(1→5)-β-D-galactofuranosyl-(1→5)-[β-D-glucopyranosyl-(1→6)]-β-D-galactofuranosyl-(1→5)-β-D-galactofuranosyl-(1→5)-[β-D-glucopyranosyl-(1→6)]-β-D-galactofuranoside (NaAG5)

Using the procedure described for the synthesis of **NaAG1**, pentadesaccharide **21** (25 mg, 3.4 μmol) in 3:1 CH_3_OH–CH_2_Cl_2_ (0.45 mL) was converted to **NaAG5** (5 mg, 75%), which was obtained as a white foam. *R*_f_ 0.23 (*n*-propanol–H_2_O–AcOH, 6:2:1); [α]_D_ –82.0 (*c* 0.5, H_2_O); ^1^H NMR (600 MHz, D_2_O, δ_H_) 5.30 (d, *J* = 2.0 Hz, 1H), 5.27–5.26 (m, 4H), 5.19–5.18 (m, 4H), 4.96 (d, *J* = 2.4 Hz, 1H), 4.53–4.47 (m, 5H), 4.17–4.00 (m, 42H), 3.95–3.92 (m, 6H), 3.92–3.85 (m, 8H), 3.80–3.79 (m, 8H), 3.75–3.69 (m, 8H), 3.51–3.44 (m, 10H), 3.41–3.36 (m, 5H), 3.36–3.29 (m, 7H), 1.64–1.54 (m, 4H), 1.38–1.29 (m, 8H); ^13^C NMR (151 MHz, D_2_O, δ_C_) 107.1, 107.03 (x 3), 107.02 (x 3), 107.01, 106.97, 106.9, 102.8 (x 4), 102.7, 82.6, 81.50, 81.45, 81.4, 81.3, 80.9, 76.5, 76.4, 76.3, 75.9, 75.6, 75.3, 74.4, 74.0, 73.9, 73.10, 73.05, 70.5, 69.9, 69.6, 68.7, 62.7, 61.7, 61.11, 61.07, 60.7, 51.2, 30.2, 28.5, 28.2, 28.1, 27.9, 25.8, 25.0; HRMS (ESI) Calcd for (M + Na) C_98_H_167_N_3_O_76_Na: 2624.9187. Found 2624.9204.

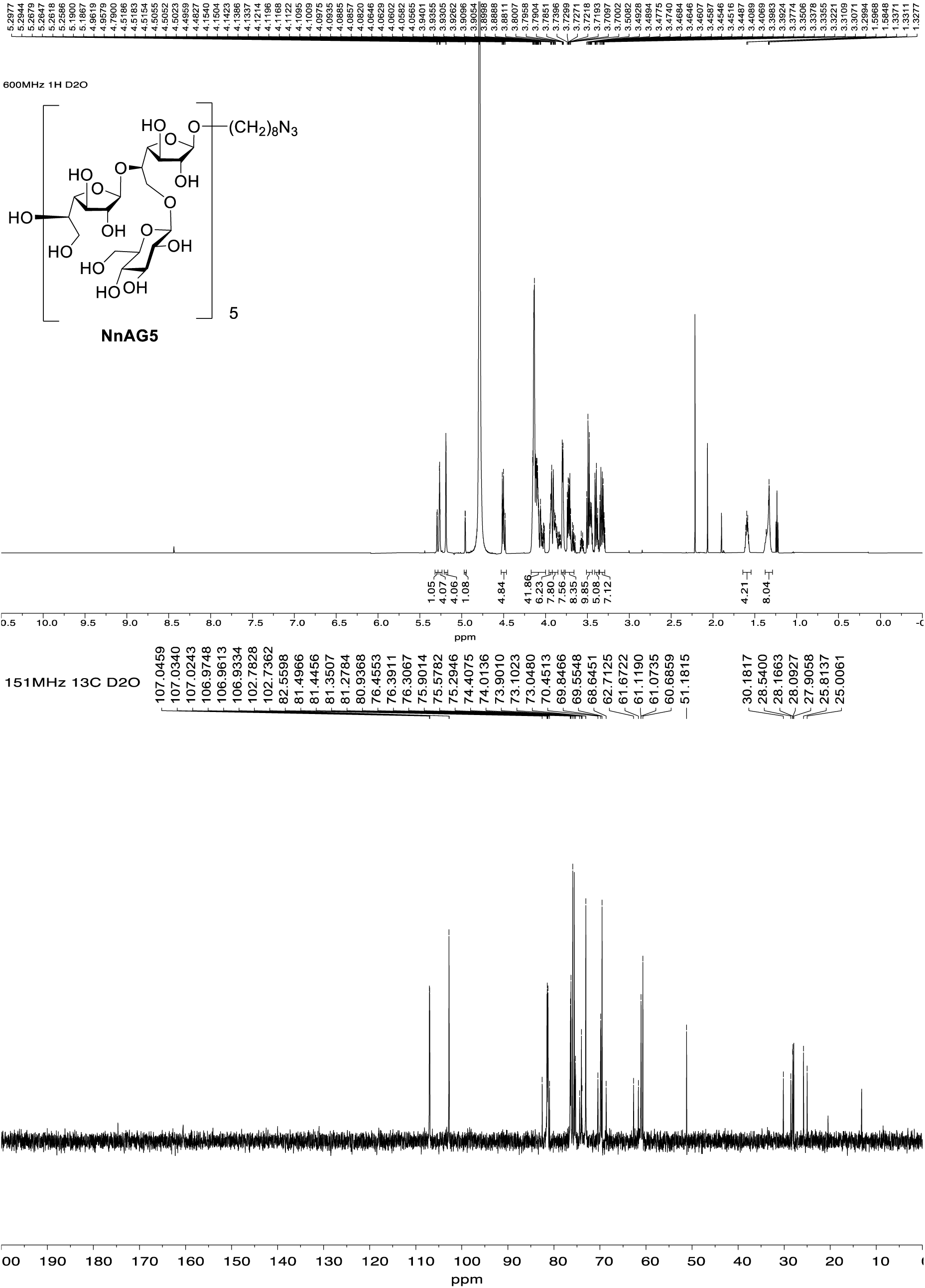

#### 8-Azidooctyl β-D-galactofuranosyl-(1→5)-[β-D-glucopyranosyl-(1→6)]-β-D-galactofuranosyl-(1→5)-β-D-galactofuranosyl-(1→5)-[β-D-glucopyranosyl-(1→6)]-β-D-galactofuranosyl-(1→5)-β-D-galactofuranosyl-(1→5)-[β-D-glucopyranosyl-(1→6)]-β-D-galactofuranosyl-(1→5)-β-D-galactofuranosyl-(1→5)-[β-D-glucopyranosyl-(1→6)]-β-D-galactofuranosyl-(1→5)-β-D-galactofuranosyl-(1→5)-[β-D-glucopyranosyl-(1→6)]-β-D-galactofuranosyl-(1→5)-β-D-galactofuranosyl-(1→5)-[β-D-glucopyranosyl-(1→6)]-β-D-galactofuranoside (NaAG6)

Using the procedure described for the synthesis of **NaAG1**, octadecasaccharide **23** (17 mg, 1.9 μmol) in 3:1 CH_3_OH–CH_2_Cl_2_ (0.2 mL) was converted to **NaAG6** (5 mg, 80%), which was obtained as a white foam. *R*_f_ 0.2 (*n*-propanol–H_2_O–AcOH, 6:2:1); [α]_D_ –50.0 (*c* 0.4, H_2_O); ^1^H NMR (500 MHz, D_2_O, δ_H_) 5.29 (d, *J* = 2.0 Hz, 1H), 5.27–5.26 (m, 5H), 5.19–5.18 (m, 5H), 4.96 (d, *J* = 2.4 Hz, 1H), 4.52–4.48 (m, 6H), 4.20–4.00 (m, 47H), 3.97–3.85 (m, 17H), 3.85–3.77 (m, 11H), 3.75–3.63 (m, 10H), 3.60–3.42 (m, 13H), 3.42–3.27 (m, 14H), 1.62–1.57 (m, 4H), 1.41–1.31 (m, 8H); ^13^C NMR (126 MHz, D_2_O, δ_C_) 107.029 (x 5), 107.026 (x 5), 106.95 (x 2), 102.8 (x 5), 102.7, 82.6, 81.47, 81.45, 81.34, 81.27, 81.2, 80.9, 76.44, 76.38, 76.3, 76.2, 75.9, 75.6, 75.4, 75.33, 75.28, 74.4, 74.0, 73.9, 73.1, 73.0, 70.4, 69.8, 69.5, 68.6, 62.7, 61.1, 60.7, 51.2, 28.5, 28.2, 28.1, 27.9, 25.8, 25.0; HRMS (ESI) Calcd for (M + Na) C_116_H_197_N_3_O_91_Na: 3111.0772 Found 3111.0783.

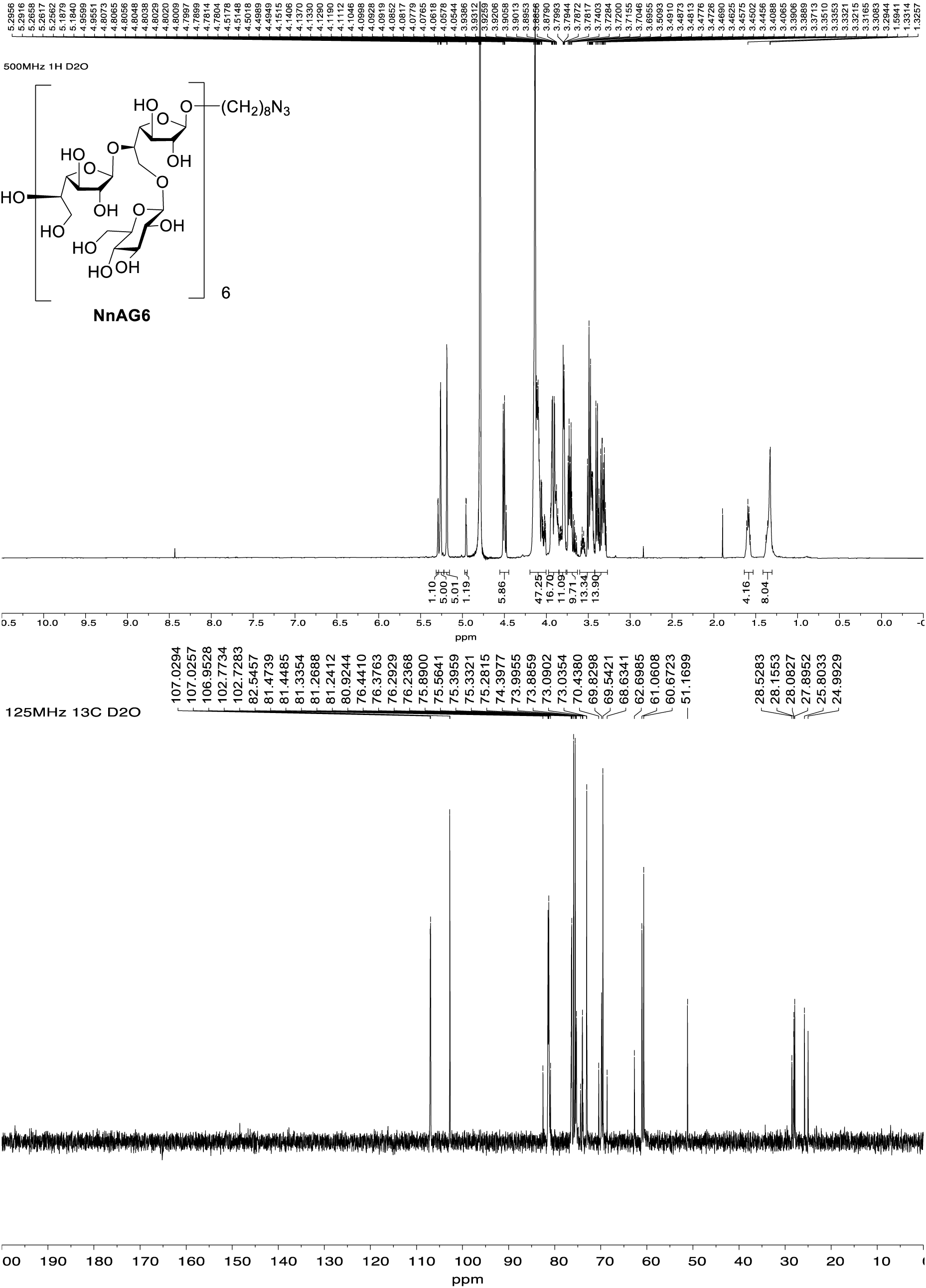

#### *p*-Tolyl 2,3-di-*O*-benzoyl-6-*O*-*t*-butyldiphenylsilyl-1-thio-β-D-galactofuranoside (1)

To a solution of **5** (0.35 g, 0.7 mmol) in pyridine (1.2 mL) and CH_2_Cl_2_ (5.8 mL) at 0 °C, *t*-butyldiphenylsilyl chloride (1.1 mL, 4.2 mmol) was added dropwise. The solution was then stirred overnight while warming to room temperature before CH_3_OH (5 mL) was added. After stirring for 30 min, the solution was poured into a satd aq NaHCO_3_ soln (50 mL) and then extracted with CH_2_Cl_2_ (50 mL). The organic layer was washed with brine (15 mL), dried over Na_2_SO_4_, filtered and the filtrate was concentrated. The syrupy residue was purified by chromatography (hexanes–EtOAc, 7:1) to afford **1** (0.29 g, 77%) as a thick syrup. *R*_f_ 0.1 (hexane–EtOAc, 10:1); [α]_D_ –27.9 (*c* 1.76, CHCl_3_); ^1^H NMR (600 MHz, CDCl_3_, δ_H_) δ 8.16–8.10 (m, 2H, ArH), 8.07–8.00 (m, 2H, ArH), 7.67–7.65 (m, 4H, ArH), 7.64–7.54 (m, 2H, ArH), 7.49 (app t, *J* = 7.8 Hz, 2H, ArH), 7.46–7.36 (m, 6H, ArH), 7.34 (app t, *J* = 7.4 Hz, 4H, ArH), 7.08 (d, *J* = 7.9 Hz, 2H, ArH), 5.73 (dd, *J* = 5.4, 1.8 Hz, 1H, H-3), 5.70 (d, *J* = 1.7 Hz, 1H, H-1), 5.68 (app t, *J* = 1.8 Hz, 1H, H-2), 4.76 (dd, *J* = 5.4, 2.4 Hz, 1H, H-4), 4.21–4.18 (m, 1H, H-5), 3.83 (dd, *J* = 10.2, 6.3 Hz, 1H, H-6a), 3.77 (dd, *J* = 10.2, 6.7 Hz, 1H, H-6b), 2.45 (d, *J* = 7.6 Hz, 1H, OH), 2.33 (s, 3H, ArC<u>H</u>_3_), 1.05 (s, 9H, (C<u>H</u>_3_)_3_C); ^13^C NMR (151 MHz, CDCl_3_, δ_C_) δ 165.7 (C=O), 165.3 (C=O), 138.1 (Ar), 135.6 (Ar), 133.6 (Ar), 133.2 (Ar), 133.10 (Ar), 133.05 (Ar), 130.0 (Ar), 129.9 (Ar), 129.80 (Ar), 129.75 (Ar), 129.2 (Ar), 129.1 (Ar), 128.6 (Ar), 128.5 (Ar), 127.8 (Ar), 91.7 (C-1), 82.0 (C-2, C-4), 78.1 (C-3), 70.8 (C-5), 64.7 (C-6), 26.8 (<u>C</u>H_3_)_3_C), 21.2 (Ar<u>C</u>H_3_), 19.2 ((CH_3_)_3_<u>C</u>); HRMS (ESI) Calcd for (M + Na) C_43_H_44_NaO_7_SSi: 755.2469. Found 755.2464.

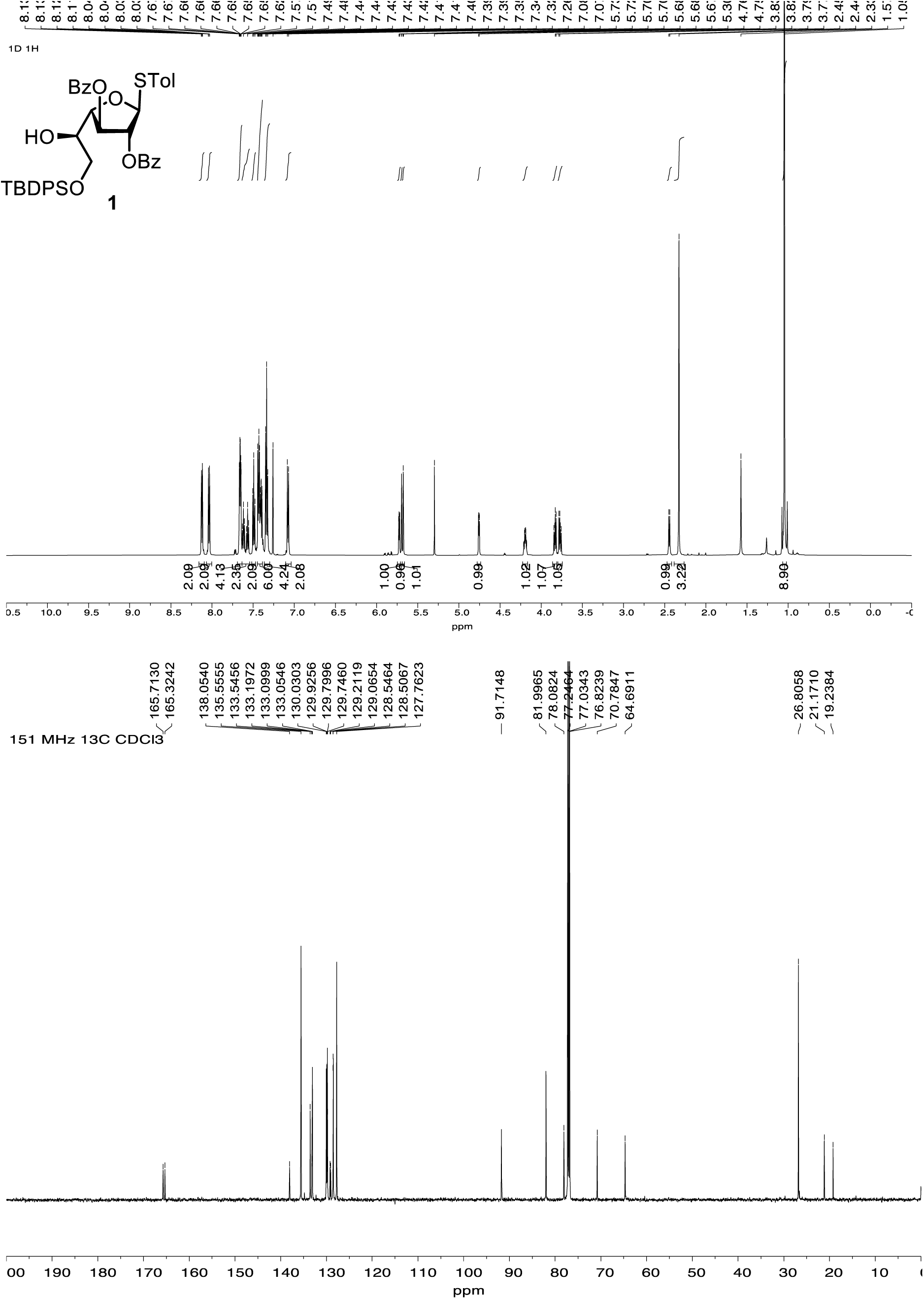

#### 2,3,6-Tri-*O*-benzoyl-5-*O*-levulinoyl-β-D-galactofuranosyl fluoride (2)

To a solution of **7** (6.3g, 9.0 mmol) in CH_2_Cl_2_ (213 mL) at 0 °C was added *N*,*N*-diethylamino sulfur trifluoride (4.3 mL, 32.5 mmol) followed by *N*-bromosuccinimide (5.68 g, 31.9 mmol). The reaction mixture was stirred overnight at room temperature and then CH_3_OH was added. The reaction mixture was diluted with CH_2_Cl_2_, and washed with satd aq NaHCO_3_ soln before the organic layer was separated. The aqueous layer was extracted again with CH_2_Cl_2_ and the combined organic layers were dried over Na_2_SO_4_. The solution was filtered and the filtrate was concentrated. The resulting residue was purified by chromatography (hexanes–EtOAc, 5:2) to afford **2** (5.3 g, 97%) as a thick syrup. *R*_f_ 0.2 (hexane–EtOAc, 2:1); [α]_D_ +28.2 (*c* 3.4, CHCl_3_); ^1^H NMR (600 MHz, CDCl_3_, δ_H_) δ 8.12–8.07 (m, 2H, ArH), 8.07–8.02 (m, 2H, ArH), 7.99–7.94 (m, 2H, ArH), 7.65–7.56 (m, 2H, ArH), 7.54–7.52 (m, 1H, ArH), 7.50–7.41 (m, 4H, ArH), 7.41–7.34 (m, 2H, ArH), 6.01 (d, *J* = 58.3 Hz, 1H, H-1), 5.79 (app dt, *J* = 7.0, 4.5 Hz, 1H, H-5), 5.68 (dd, *J* = 6.4, 1.1 Hz, 1H, H-2), 5.59 (dd, *J* = 4.1 Hz, 1.1 Hz, 1H, H-3), 4.82–4.76 (m, 1H, H-4), 4.71 (dd, *J* = 12.0, 4.5 Hz, 1H, H-6a), 4.57 (dd, *J* = 12.0, 7.0 Hz, 1H, H-6b), 2.86–2.38 (m, 4H, Lev CH_2_), 2.07 (s, 3H, Lev CH_3_); ^13^C NMR (151 MHz, CDCl_3_, δ_C_) 205.9 (Lev ketone C=O)), 171.9 (Lev ester C=O)), 166.0 (Ar), 133.9 (Ar), 133.8 (Ar), 133.2 (Ar), 130.0 (Ar), 129.8 (Ar), 129.5 (Ar), 128.7 (Ar), 128.5 (Ar), 128.4 (Ar), 112.3 (d, *J* = 226.5 Hz, C-1), 84.6 (C-4), 80.6 (d, *J* = 39.9 Hz, C-2), 76.1 (C-3), 70.0 (C-5), 63.0 (C-6), 37.9 (Lev CH_2_), 29.7 (Lev CH_3_), 28.0 (Lev CH_2_); ^19^F NMR (565 MHz, CDCl_3_, δ_F_) – 125.36; HRMS (ESI) Calcd for (M + Na) C_32_H_29_FNaO_10_: 615.1637. Found 615.1638.

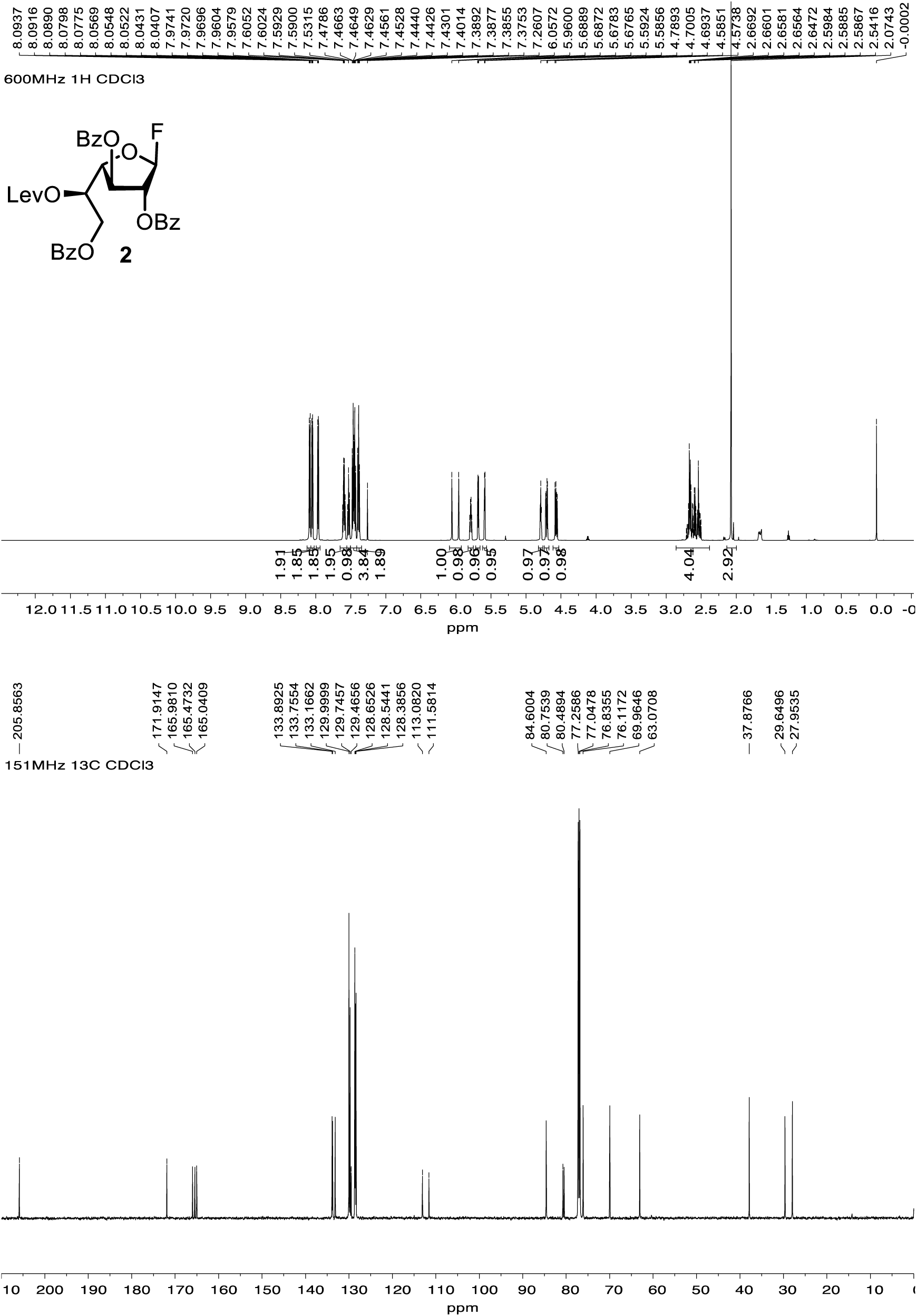

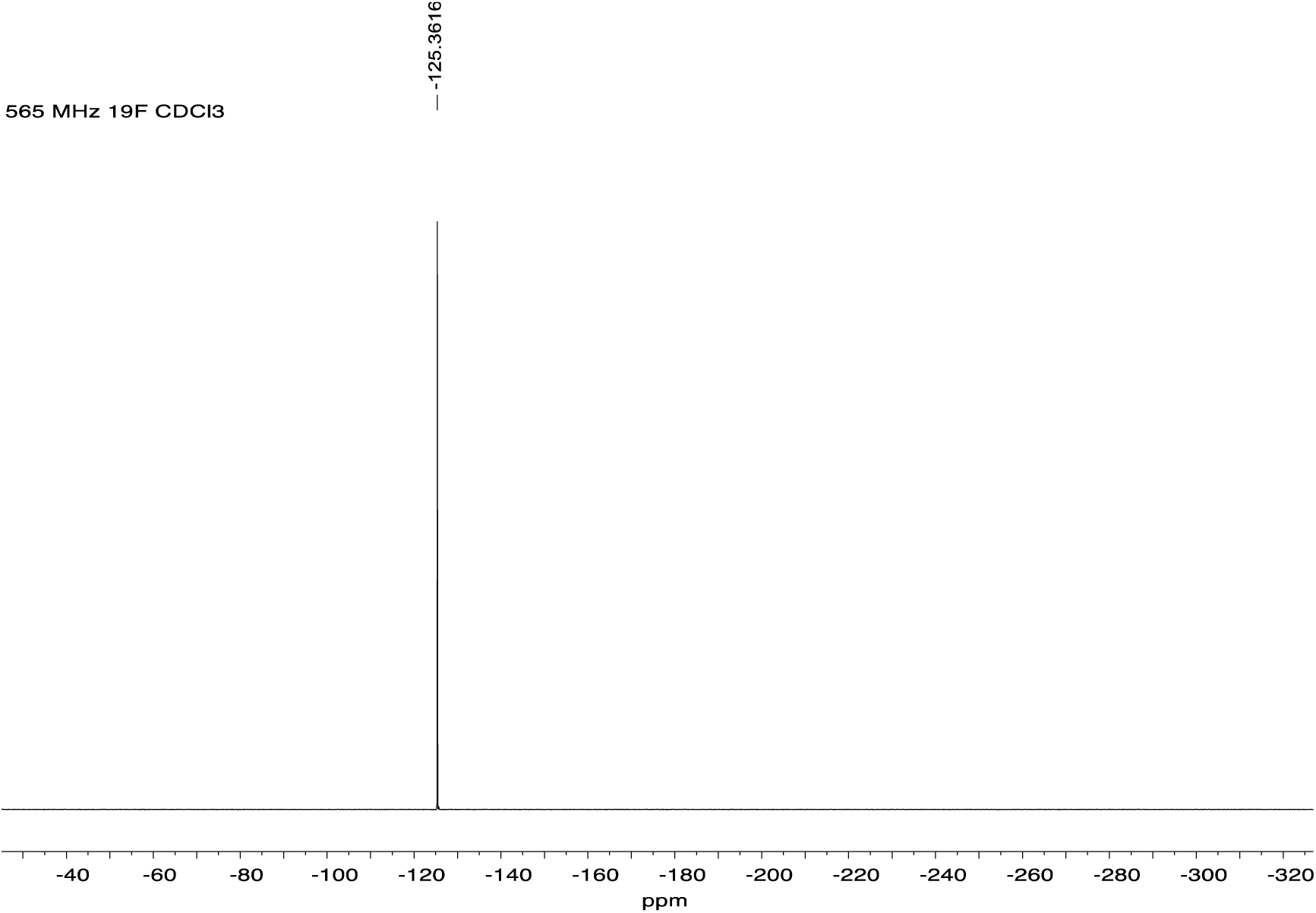

#### 2,3,4,6-Tetra-*O*-benzoyl-β-D-glucopyranosyl fluoride (3)

To a solution of thioglycoside **9** (4.71 g, 6.7 mmol) in CH_2_Cl_2_ (71 mL) at 0 °C was added *N*,*N*-diethylaminosulfur trifluoride (2.4 mL, 18.9 mmol) followed by *N*-bromosuccinimide (3.2 g, 18.0 mmol). The solution was stirred at room temperature overnight and then neutralized by the addition of satd aq NaHCO_3_ soln. The solution was extracted with CH_2_Cl_2_ (100 mL) and the CH_2_Cl_2_ layer was then washed with water (30 mL × 3). The separated CH_2_Cl_2_ layer was dried over Na_2_SO_4_, filtered and the filtrate was concentrated. The resulting residue was purified by chromatography (hexane–EtOAc, 5:1) to afford glycosyl fluoride **3** (2.78 g, 70%) as a colorless syrup. *R*_f_ 0.15 (hexane–EtOAc, 5:1); ^1^H NMR (600 MHz, CDCl_3_, δ_H_) 8.13–8.06 (m, 2H, ArH), 8.06–8.01 (m, 2H, ArH), 7.99–7.94 (m, 2H, ArH), 7.94–7.90 (m, 2H, ArH), 7.63–7.56 (m, 2H, ArH), 7.56–7.52 (m, 1H, ArH), 7.52–7.48 (m, 1H, ArH), 7.46–7.42 (m, 4H, ArH), 7.39–7.35 (m, 4H, ArH), 5.91–5.84 (m, 2H, H-3, H-4), 5.74 (dd, *J* = 51.5, 5.4 Hz, 1H, H-1), 5.63 (ddd, *J* = 9.6, 7.3, 5.4 Hz, 1H, H-2), 4.74 (dd, *J* = 12.2, 3.7 Hz, 1H, H-6a), 4.59 (dd, *J* = 12.2, 5.2 Hz, 1H, H-6b), 4.44 (ddd, *J* = 8.8, 5.1, 3.8 Hz, 1H, H-5); ^13^C NMR (151 MHz, CDCl_3_, δ_C_) 166.1 (C=O), 165.5 (C=O), 165.1 (C=O), 164.9 (C=O), 133.6 (Ar), 133.6 (Ar), 133.5 (Ar), 133.2 (Ar), 130.0 (Ar), 129.9 (Ar), 129.9 (Ar), 129.8 (Ar), 128.7 (Ar), 128.6 (Ar), 128.6 (Ar), 128.5 (Ar), 128.5 (Ar), 128.4 (Ar), 106.5 (d, *J* = 221.1 Hz, C-1), 72.1 (d, *J* = 3.0 Hz, C-5), 71.4 (d, *J* = 7.2 Hz, C-3), 71.3 (d, *J* = 16.4 Hz, C-2), 68.2 (C-4), 63.0 (C-6); ^19^F NMR (565 MHz, CDCl_3_, δ_F_) –135.19; HRMS (ESI) Calcd for (M + Na) C_34_H_27_FNaO_9_: 621.1531. Found 621.1527.

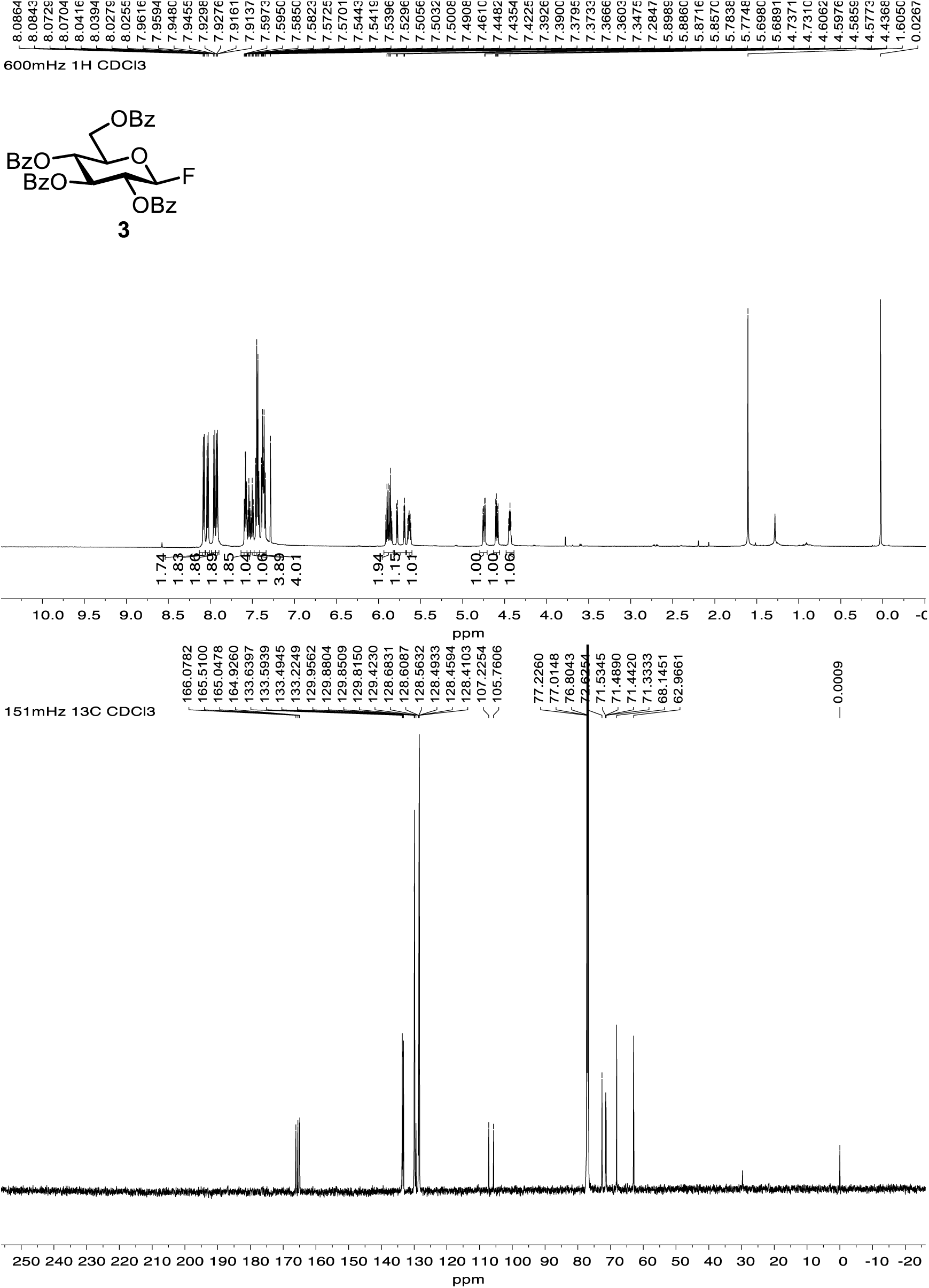

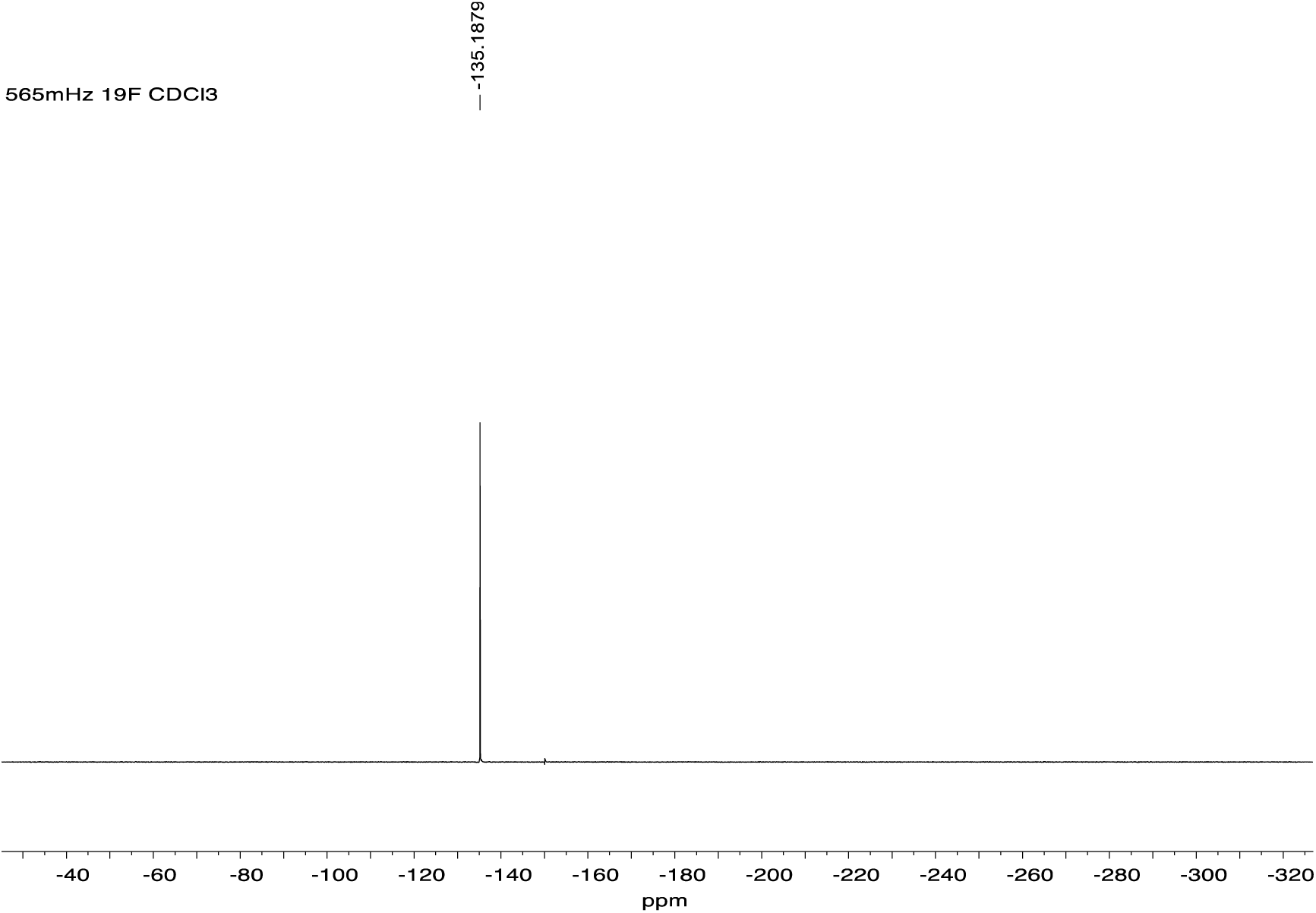

#### *p*-Tolyl 2,3-di-*O*-benzoyl-1-thio-β-D-galactofuranoside (5)

Compound **4** ^37^ (0.235 g, 0.72 mmol) was dissolved in pyridine (1.2 mL) and the solution cooled to 0 °C before benzoyl chloride (0.8 mL, mmol) was added dropwise. The reaction mixture was warmed to room temperature and stirred for 2 h. Excess benzoyl chloride was quenched by the addition of chilled water (50 mL) and the mixture was extracted with CH_2_Cl_2_ (20 mL). The CH_2_Cl_2_ layer was washed with satd aq NaHCO_3_ soln (10 mL) and water (10 mL × 3). The separated CH_2_Cl_2_ layer was dried over Na_2_SO_4_, filtered and the filtrate was concentrated. The syrupy residue obtained was dissolved in 4:1 acetic acid–H_2_O (3.6 mL) and the solution was heated at 65–70 °C overnight. After cooling to rt, the reaction mixture was concentrated and the residue was purified by column chromatography (hexanes–EtOAc, 3:2) to afford **5** (0.29 g, 83% over two steps) as a thick syrup. *R*_f_ 0.1 (hexane–EtOAc, 2:1); [α]_D_ –92.9 (*c* 1.13, CHCl_3_); ^1^H NMR (600 MHz, CDCl_3_, δ_H_) 8.16–8.09 (m, 2H, ArH), 8.08–8.03 (m, 2H, ArH), 7.65–7.55 (m, 2H, ArH), 7.53–7.41 (m, 6H, ArH), 7.14 (d, *J* = 7.9 Hz, 2H, ArH), 5.69 (s, 1H, H-1), 5.69 (app t, *J* = 1.5 Hz, 1H, H-3), 5.69–5.65 (m, 1H, H-2), 4.57 (dd, *J* = 4.8, 3.4 Hz, 1H, H-4), 4.20–4.13 (m, 1H, H-5), 3.89–3.75 (m, 2H, H-6 x 2), 2.78 (d, *J* = 7.7 Hz, 1H, O<u>H</u>), 2.34 (s, 3H, ArC<u>H</u>_3_); ^13^C NMR (151 MHz, CDCl_3_, δ_C_) δ 166.0 (<u>C</u>=O), 165.3 (<u>C</u>=O), 138.4 (Ar), 133.8 (Ar), 133.7 (Ar), 133.2 (Ar), 130.1 (Ar), 129.94 (Ar), 129.90 (Ar), 129.4 (Ar), 128.9 (Ar), 128.6 (Ar), 91.8 (C-1), 84.1 (C-4), 81.8 (C-2), 78.1 (C-3), 70.5 (C-5), 64.3 (C-6), 21.2 (Ar<u>C</u>H_3_); HRMS (ESI) Calcd for (M + Na) C_27_H_26_NaO_7_S: 517.1291. Found 517.1282.

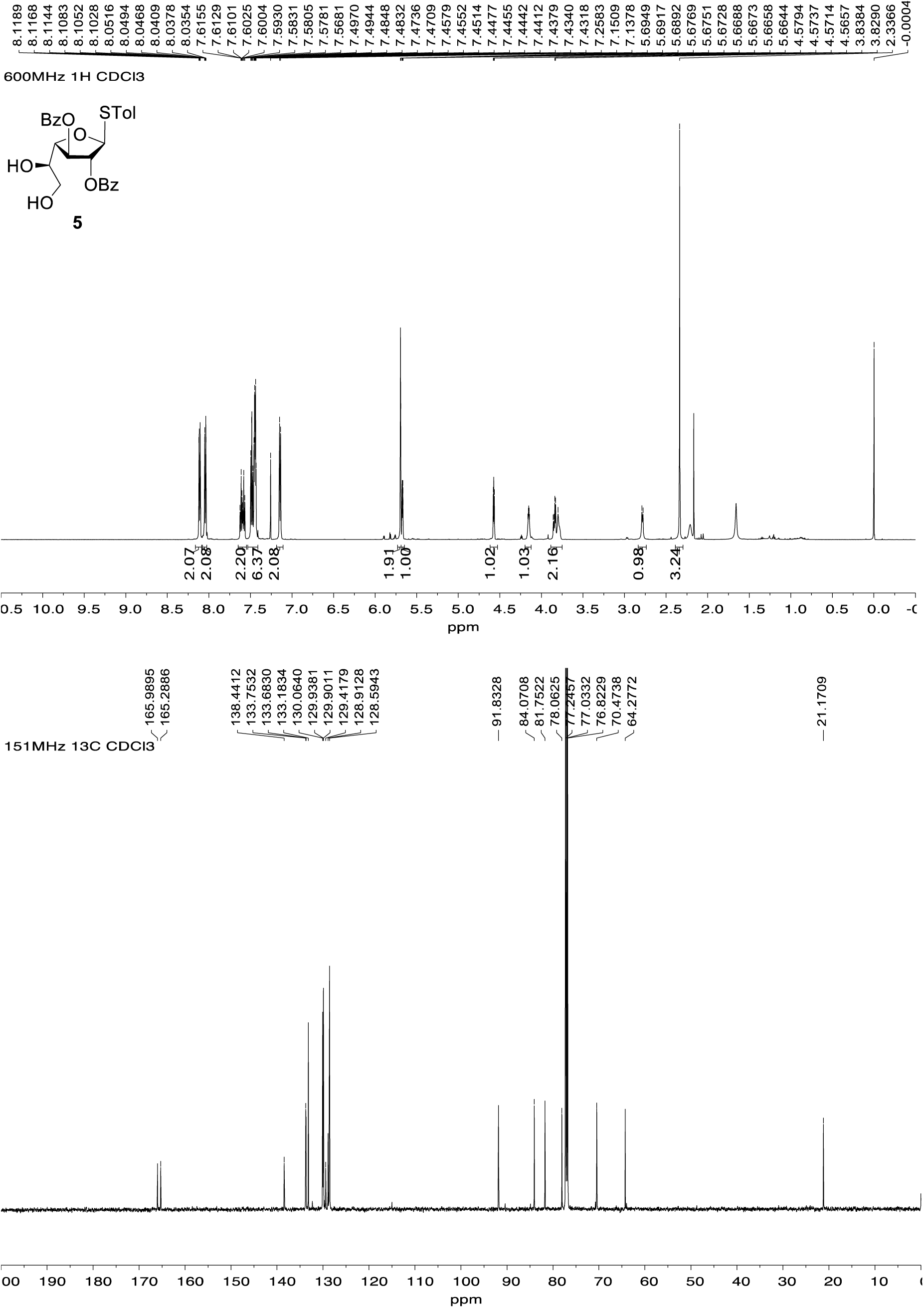

#### *p*-Tolyl 2,3,6-tri-*O*-benzoyl-5-*O*-levulinoyl-1-thio-β-D-galactofuranoside (7)

Compound **6** ^24^ (0.5 g, 0.6 mmol) was dissolved in a solution of CH_2_Cl_2_–CH_3_OH (50:1, 12 mL) at 0 °C and *p*-toluenesulfonic acid (0.11 g, 0.6 mmol) was added. The reaction mixture was stirred overnight at rt. The mixture was diluted with EtOAc, poured into a satd aq NaHCO_3_ soln (20 mL) and extracted with EtOAc (40 mL). The organic layer was washed with water, dried over Na_2_SO_4_, filtered and the filtrate was concentrated to give the corresponding alcohol as a yellow syrup. This material was dissolved in pyridine (10 mL) and the solution was cooled to 0 °C before the addition of benzoyl chloride (1.1 mL, 10.1 mmol) dropwise. The reaction mixture was warmed to room temperature and stirred overnight. Excess benzoyl chloride was quenched by the addition of chilled water (30 mL) and the solution was extracted with CH_2_Cl_2_ (20 mL). The CH_2_Cl_2_ layer was washed with 10% aq copper sulfate soln (20 mL × 3) and then water (20 mL). The separated CH_2_Cl_2_ layer was dried over Na_2_SO_4_, filtered and the filtrate was concentrated. This material was then dissolved in CH_2_Cl_2_ (5 mL) and then levulinic acid (0.12 mL, 1.2 mmol), *N*-(3-dimethylaminopropyl)-*N’*-ethylcarbodiimide hydrochloride (0.28 g, 1.8 mmol), and 4-(dimethylamino) pyridine (0.36 g, 2.9 mmol) were added. The mixture was stirred for 4 h at rt. The reaction mixture washed with a satd aq NaHCO_3_ soln (5 mL) and brine (10 mL). The organic layer was then dried over Na_2_SO_4_, filtered and the filtrate was concentrated. The syrup residue was purified by column chromatography (hexanes–EtOAc, 2:1) to afford **7** (0.4 g, 83% over two steps) as a white foam. *R*_f_ 0.2 (hexane–EtOAc, 2:1); [α]_D_ –28.3 (*c* 2.3, CHCl_3_); ^1^H NMR (500 MHz, CDCl_3_, δ_H_) 8.11–8.04 (m, 4H, ArH), 8.00–7.96 (m, 2H, ArH), 7.62–7.38 (m, 11H, ArH), 7.12–7.08 (m, 2H, ArH), 5.78 (app dt, *J* = 7.0, 4.3 Hz, 1H, H-5), 5.72 (d, *J* = 1.9 Hz, 1H, H-1), 5.66 (app t, *J* = 1.9 Hz, 1H, H-2), 5.60 (dd, *J* = 4.6, 1.9 Hz, 1H, H-3), 4.79 (app t, *J* = 4.6 Hz, 1H, H-4), 4.66 (dd, *J* = 11.9, 4.3 Hz, 1H, H-6a), 4.54 (dd, *J* = 11.9, 7.0 Hz, 1H, H-6b), 2.71–2.52 (m, 4H, Lev CH_2_), 2.32 (s, 3H, ArC<u>H</u>_3_), 2.07 (s, 3H, Lev CH_3_); ^13^C NMR (151 MHz, CDCl_3_, δ_C_) 205.9 (Lev ketone C=O), 172.0 (Lev ester C=O), 166.0 (C=O), 165.4 (C=O), 165.3 (C=O), 138.3 (Ar), 133.6 (Ar), 133.1 (Ar), 133.1 (Ar), 130.1 (Ar), 129.93 (Ar), 129.87 (Ar), 129.8 (Ar), 129.6 (Ar), 128.9 (Ar), 128.6 (Ar), 128.4 (Ar), 91.4 (C-1), 82.0 (C-2), 81.1 (C-4), 77.4 (C-3), 69.9 (C-5), 63.2 (C-6), 37.9 (Lev CH_2_), 29.7 (Lev CH_3_), 28.0 (Lev CH_2_), 21.2 (Ar<u>C</u>H_3_); HRMS (ESI) Calcd for (M + Na) C_39_H_36_NaO_10_S: 719.1921. Found 719.1929.

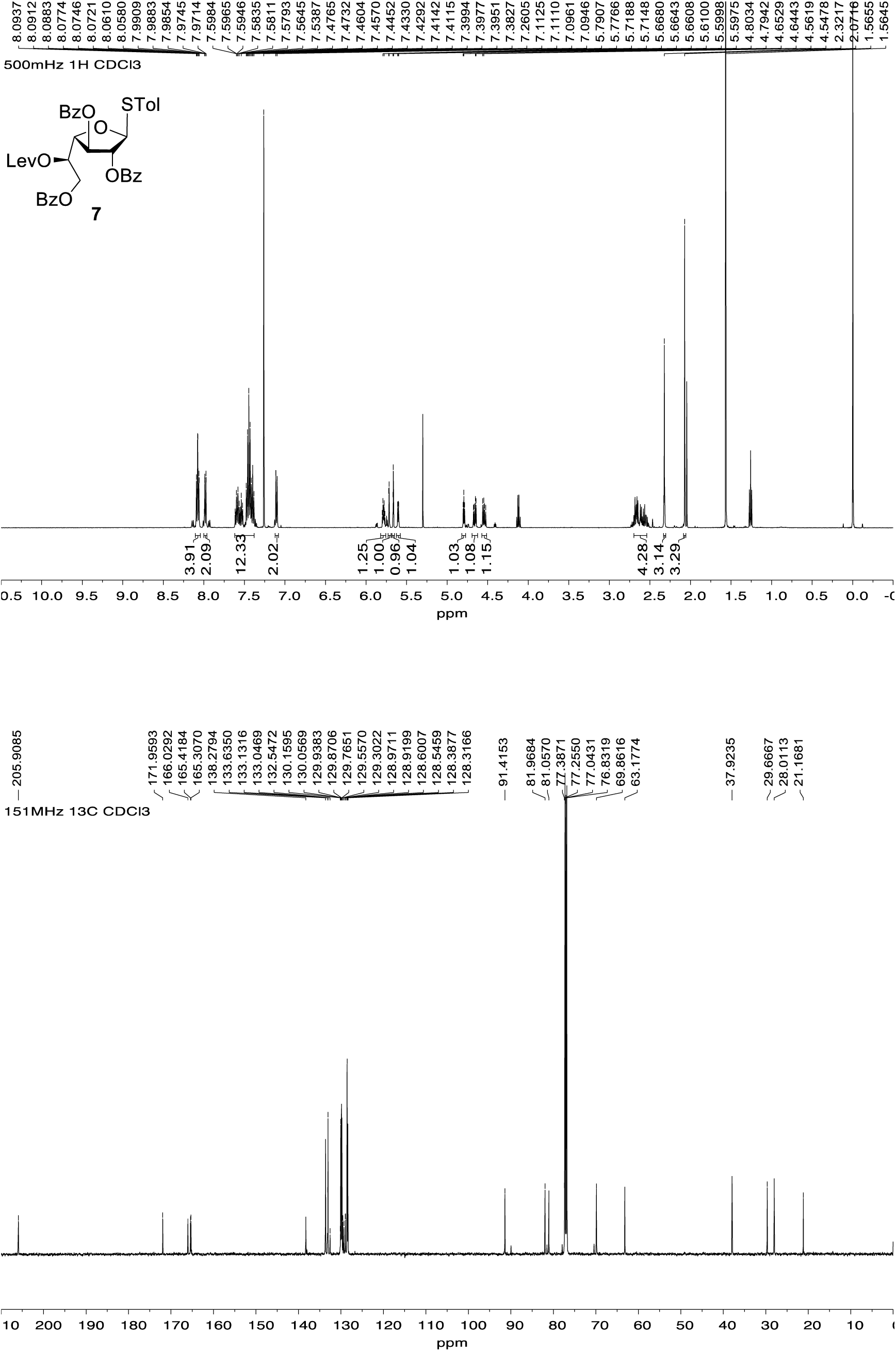

#### *p*-Tolyl 2,3,4,6-tetra-*O*-benzoyl-1-thio-β-D-glucopyranoside (9)

To a solution of **8** ^25^(1.07 g, 1.5 mmol) in CH_2_Cl_2_ (2.8 mL) was added *p*-toluenethiol (0.21 g, 1.8 mmol). The mixture was cooled to 0 °C and then boron trifluoride diethyl etherate (2.1 mL, 1.8 mmol) was added dropwise. The solution was stirred at room temperature overnight and then neutralized by the addition of satd aq NaHCO_3_ soln dropwise. The solution was extracted with CH_2_Cl_2_ (100 mL) and the CH_2_Cl_2_ layer was washed with water (20 mL × 3). The separated CH_2_Cl_2_ layer was dried over Na_2_SO_4_, filtered and the filtrate was concentrated. The residue was recrystallized from EtOAc to give **9** as a white foam (460 mg). The mother liquor was concentrated and then purified by chromatography (hexane–EtOAc, 6:1) to give additional **9** (total yield = 0.97 g, 90%). *R*_f_ 0.15 (hexane–EtOAc, 7:1); mp 115 °C; ^1^H NMR (500 MHz, CDCl_3_, δ_H_) 8.04 (dd, *J* = 8.4, 1.4 Hz, 2H, ArH), 8.00–7.95 (m, 2H, ArH), 7.92–7.87 (m, 2H, ArH), 7.82–7.76 (m, 2H, ArH), 7.59 (ddt, *J* = 8.7, 7.3, 1.3 Hz, 1H, ArH), 7.55–7.31 (m, 11H, ArH), 7.29–7.23 (m, 2H, ArH), 6.98–6.89 (m, 2H, ArH), 5.90 (app t, *J* = 9.8, 9.7 Hz, 1H, H-3), 5.59 (app t, *J* = 9.8 Hz, 1H, H-4), 5.45 (app t, *J* = 9.7 Hz, 1H, H-2), 4.98 (d, *J* = 10.0 Hz, 1H, H-1), 4.68 (dd, *J* = 12.2, 2.8 Hz, 1H, H-6b), 4.48 (dd, *J* = 12.2, 5.7 Hz, 1H, H-6a), 4.17 (ddd, *J* = 10.0, 5.7, 2.8 Hz, 1H, H-5), 2.34 (s, 3H, ArC<u>H</u>_3_); ^13^C NMR (126 MHz, CDCl_3_, δ_C_) 166.1 (C=O), 165.8 (C=O), 165.2 (C=O), 165.0 (C=O), 138.6 (Ar), 133.9 (Ar), 133.5 (Ar), 133.3 (Ar), 133.2 (Ar), 133.1 (Ar), 129.88 (Ar), 129.86 (Ar), 129.8 (Ar), 129.6 (Ar), 129.3 (Ar), 128.8 (Ar), 128.7 (Ar), 128.4 (Ar), 128.3 (Ar), 127.6 (Ar), 86.2 (C-1), 76.3 (C-3), 74.2 (C-2), 70.5 (C-5), 69.4 (C-4), 63.1 (C-6), 21.1 (Ar<u>C</u>H_3_); HRMS (ESI) Calcd for (M + Na) C_41_H_34_O_9_NaS: 725.1816. Found 725.1812.

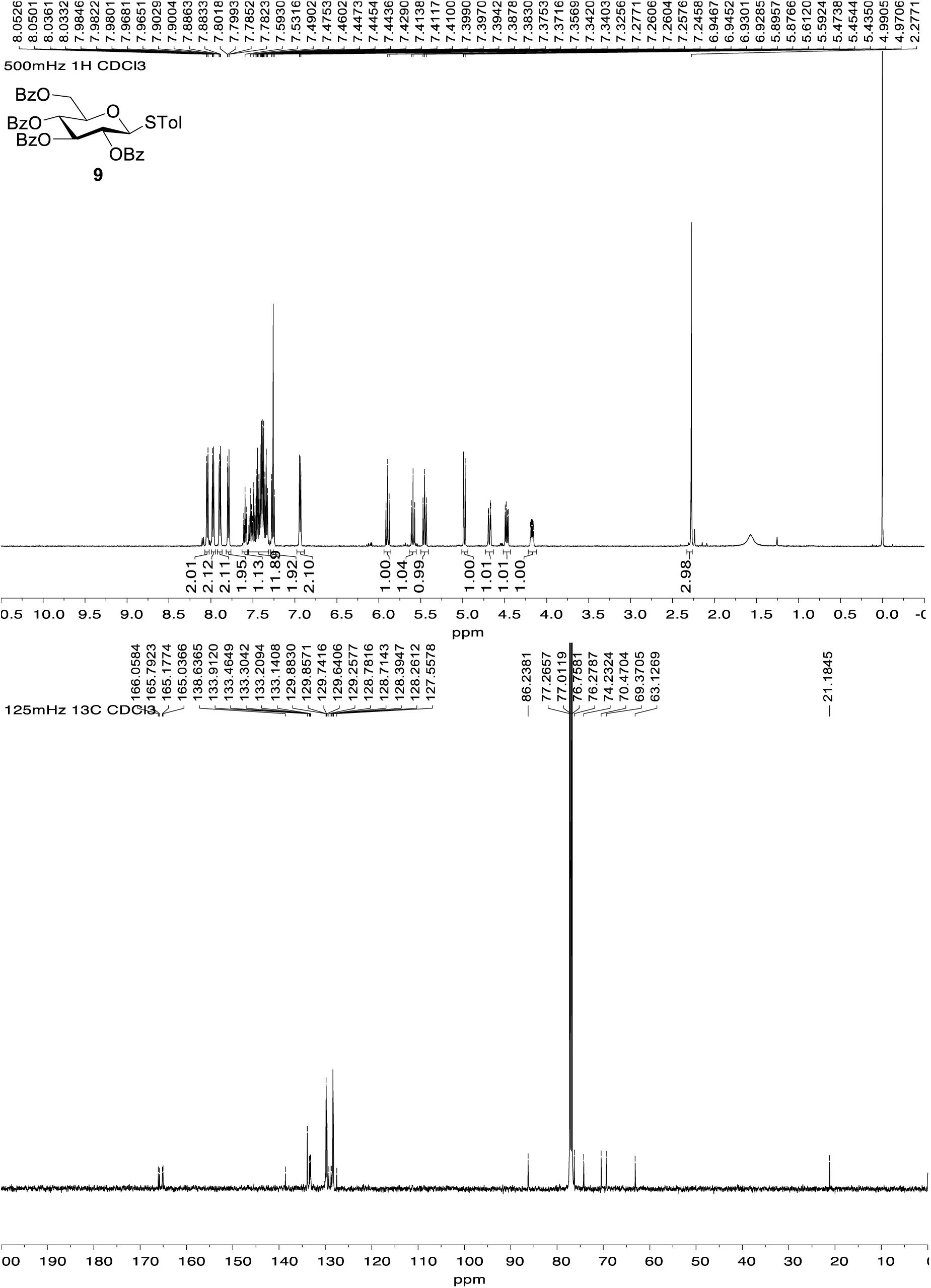

#### *p*-Tolyl 2,3,6-tri-*O*-benzoyl-5-*O*-levulinoyl-β-D-galactofuranosyl-(1→5)-2,3-di-*O*-benzoyl-6-*O*-*t*-butyldiphenylsilyl-1-thio-β-D-galactofuranoside (10)

A mixture of **1** (0.87 g, 1.2 mmol) and 4 Å molecular sieves (powder, 5.3 g) in CH_2_Cl_2_ (17 mL) was stirred at room temperature for 30 min under an Ar atmosphere, and then cooled to –10 °C before bis(cyclopentadienyl)zirconium (IV) dichloride (87 mg, 0.3 mmol) and silver trifluoromethanesulfonate (152 mg, 0.6 mmol) were added successively. After stirring for 10 min at –10 °C, glycosyl fluoride **2** (0.77 g, 1.3 mmol) in CH_2_Cl_2_ (2 mL) was added dropwise over 30 min. After stirring at –10 °C for 4 h, triethylamine was added and the resulting mixture was filtered through Celite. The filtrate was diluted with CH_2_Cl_2_ (30 mL) and washed with satd aq NaHCO_3_ soln. The organic layer was separated and the aqueous layer was extracted with CH_2_Cl_2_ (15 mL × 3). The combined organic layers were dried over Na_2_SO_4_, filtered, the filtrate was concentrated and the residue was purified by chromatography (hexanes–EtOAc, 5:2) to afford **10** (1.43 g, 92%) as a white foam. *R*_f_ 0.1 (hexane–EtOAc, 3:1); [α]_D_ –30.77 (*c* 0.78, CHCl_3_); ^1^H NMR (600 MHz, CDCl_3_, δ_H_) 8.04–7.98 (m, 4H, ArH), 7.94–7.90 (m, 2H, ArH), 7.90–7.85 (m, 4H, ArH), 7.63–7.62 (m, 4H, ArH), 7.60–7.53 (m, 2H, ArH), 7.49–7.36 (m, 9H, ArH), 7.36–7.26 (m, 10H, ArH), 7.21–7.18 (m, 2H, ArH), 7.00–6.98 (m, 2H, ArH), 5.76 (dd, *J* = 6.0, 2.8, 1H, H-3), 5.68 (dt, *J* = 7.8, 3.9 Hz, 1H, H-5’), 5.62 (d, *J* = 3.0 Hz, 1H, H-1), 5.61–5.58 (m, 2H, H-2, H-2’), 5.54 (app s, 1H, H-1’), 5.46 (dd, *J* = 5.2, 1.7 Hz, 1H, H-3’), 4.78 (dd, *J* = 5.2, 3.9 Hz, 1H, H-4’), 4.73–4.68 (m, 1H, H-4), 4.51 (dd, *J* = 12.0, 3.9 Hz, 1H, H-6a’), 4.46 (dd, *J* = 12.0, 7.8 Hz, 1H, H-6b’), 4.35 (dt, *J* = 6.5, 3.6 Hz, 1H, H-5), 3.97 (dd, *J* = 10.8, 6.5 Hz, 1H, H-6a), 3.93 (dd, *J* = 10.8, 3.6 Hz, 1H, H-6b), 2.65–2.34 (m, 4H, Lev CH_2_), 2.26 (s, 3H, ArC<u>H</u>_3_), 2.00 (s, 3H, Lev CH_3_), 0.98 (s, 9H, (C<u>H</u>_3_)_3_C); ^13^C NMR (151 MHz, CDCl_3_, δ_C_) δ 205.9 (Lev ketone C=O), 171.9 (Lev ester C=O), 165.9 (Ar), 165.5 (Ar), 165.40 (Ar), 165.36 (Ar), 165.0 (Ar), 137.9 (Ar), 135.5 (Ar), 133.45 (Ar), 133.43 (Ar), 133.36 (Ar), 133.3 (Ar), 132.90 (Ar), 132.87 (Ar), 132.82 (Ar), 132.79 (Ar), 130.0 (Ar), 129.9 (Ar), 129.8 (Ar), 129.73 (Ar), 129.69 (Ar), 129.60 (Ar), 129.55 (Ar), 129.1 (Ar), 129.0 (Ar), 128.86 (Ar), 128.85 (Ar), 128.5 (Ar), 128.4 (Ar), 128.2 (Ar), 127.8 (Ar), 127.7 (Ar), 105.3 (C-1’), 90.6 (C-1), 81.7 (C-2’), 81.6 (C-2), 81.5 (C-4’), 81.0 (C-4), 77.1 (C-3, C-3’), 75.9 (C-5), 70.0 (C-5’), 63.6 (C-6’), 63.4 (C-6), 37.9 (Lev CH_2_), 29.6 (Lev CH_3_), 27.9 (Lev CH_2_), 26.7 ((<u>C</u>H_3_)_3_C), 21.1 (Ar<u>C</u>H_3_), 19.1 (CH_3_)_3_<u>C</u>); HRMS (ESI) Calcd for (M + Na) C_75_H_72_NaO_17_SSi: 1327.4151. Found 1327.4148.

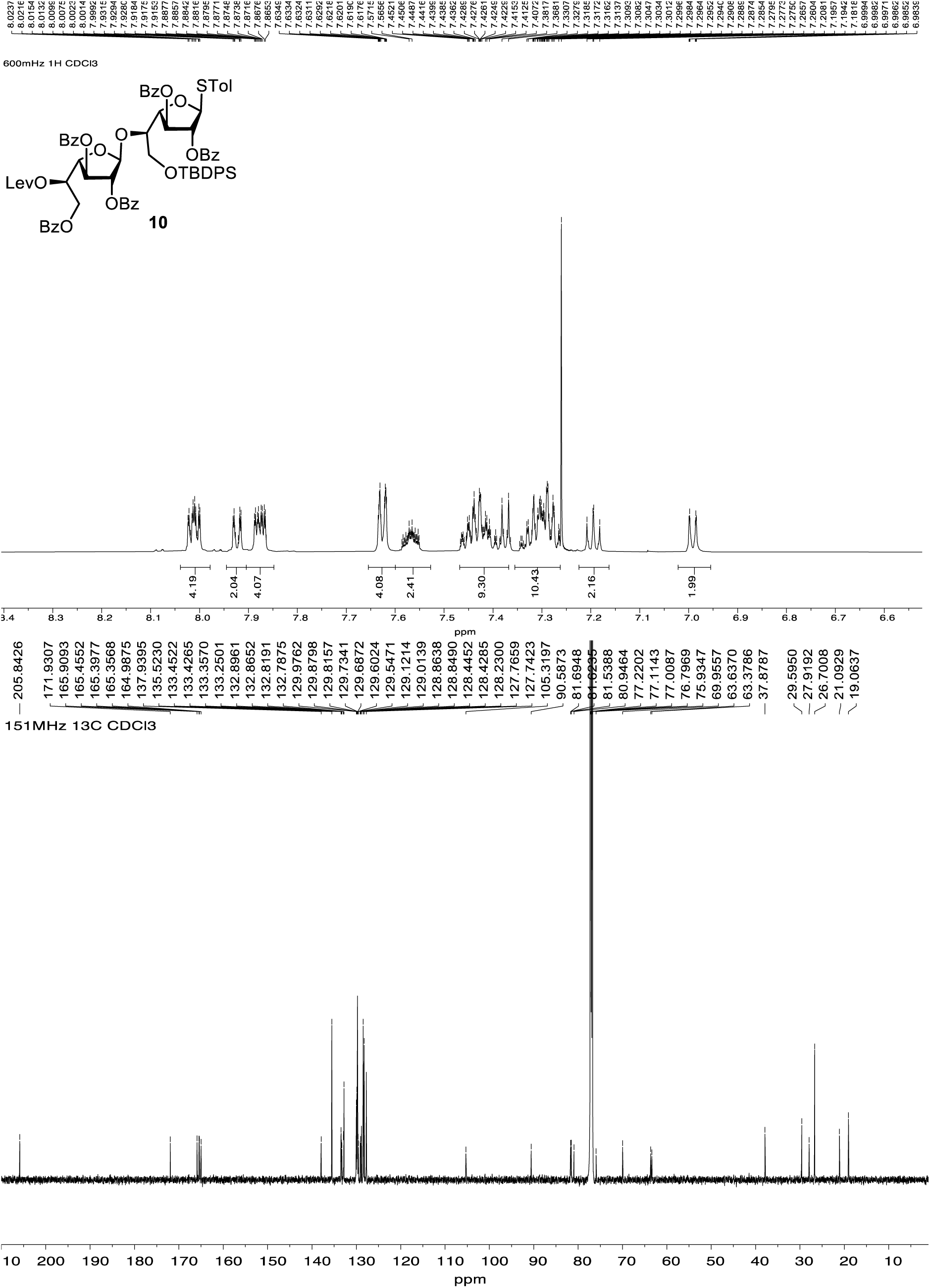

#### *p*-Tolyl 2,3,6-tri-*O*-benzoyl-5-*O*-levulinoyl-β-D-galactofuranosyl-(1→5)-2,3-di-*O*-benzoyl-1-thio-β-D-galactofuranoside (11)

Compound **10** (97 mg, 74 μmol) was dissolved in a solution of pyridine–THF (1:1, 0.8 mL) at 0 °C and 70% HF·pyridine (0.1 mL) was added dropwise. After the reaction was stirred for 15 h at rt, the mixture was diluted with EtOAc, poured into a satd aq NaHCO_3_ soln (10 mL) and extracted with EtOAc (30 mL). The organic layer was washed with water, dried over Na_2_SO_4_, filtered and the filtrate was concentrated. The resulting crude residue was purified by chromatography (hexane– EtOAc, 5:2) to afford alcohol **11** (65 mg, 82%) as a white foam. *R*_f_ 0.18 (hexane–EtOAc, 5:2); [α]_D_ – 38.5 (*c* 0.52, CHCl_3_); ^1^H NMR (600 MHz, CDCl_3_, δ_H_) 8.12–8.07 (m, 2H, ArH), 8.04–8.02 (m, 2H, ArH), 7.99–7.98 (m, 2H, ArH), 7.95–7.93 (m, 2H, ArH), 7.89–7.87 (m, 2H, ArH), 7.62–7.54 (m, 2H, ArH), 7.52–7.46 (m, 3H, ArH), 7.46–7.41 (m, 6H, ArH), 7.40–7.35 (m, 2H, ArH), 7.35–7.31 (m, 2H, ArH), 7.25–7.21 (m, 2H, ArH), 7.10–7.05 (m, 2H, ArH), 5.80 (dd, *J* = 5.5, 2.2 Hz, 1H, H-3), 5.69 (d, *J* = 2.2 Hz, 1H, H-1), 5.65 (app dt, *J* = 7.7, 3.9 Hz, 1H, H-5’), 5.63–5.60 (m, 2H, H-1’, H-2), 5.56–5.55 (m, 2H, H-2’, H-4’), 4.77–4.72 (m, 1H, H-3’), 4.70 (dd, *J* = 5.5, 4.2 Hz, 1H, H-4), 4.48 (dd, *J* = 12.1, 3.9 Hz, 1H, H-6a’), 4.42 (dd, *J* = 12.1, 7.7 Hz, 1H, H-6b’), 4.33 (app dt, *J* = 6.0, 4.2 Hz, 1H, H-5), 4.00–3.93 (m, 2H, H-6a, H-6b), 2.69–2.48 (m, 4H, Lev CH_2_), 2.28 (s, 3H, ArCH_3_), 2.05 (s, 3H, Lev CH_3_); ^13^C NMR (151 MHz, CDCl_3_) δ 206.0 (Lev ketone C=O), 172.0 (Lev ester C=O), 166.00 (Ar), 165.95 (Ar), 165.50 (Ar), 165.47 (Ar), 165.4 (Ar), 138.4 (Ar), 133.7 (Ar), 133.60 (Ar), 133.56 (Ar), 133.4 (Ar), 133.2 (Ar), 133.01 (Ar), 130.03 (Ar), 130.0 (Ar), 129.9 (Ar), 129.82 (Ar), 129.78 (Ar), 129.7 (Ar), 129.6 (Ar), 129.3 (Ar), 128.92 (Ar), 128.90 (Ar), 128.87 (Ar), 128.8 (Ar), 128.6 (Ar), 128.5 (Ar), 128.3 (Ar), 105.3 (C-1’), 91.3 (C-1), 83.0 (C-4), 82.7 (C-4’), 81.7 (C-2, C-2’), 80.9 (C-3’), 77.4 (C-3), 75.9 (C-5), 69.9 (C-5’), 63.6 (C-6’), 62.2 (C-6), 37.9 (Lev CH_2_), 29.7 (Lev CH_3_), 28.0 (Lev CH_2_), 21.1 (Ar<u>C</u>H_3_); HRMS (ESI) Calcd for (M + Na) C_59_H_54_NaO_17_S: 1089.2974. Found 1089.2979.

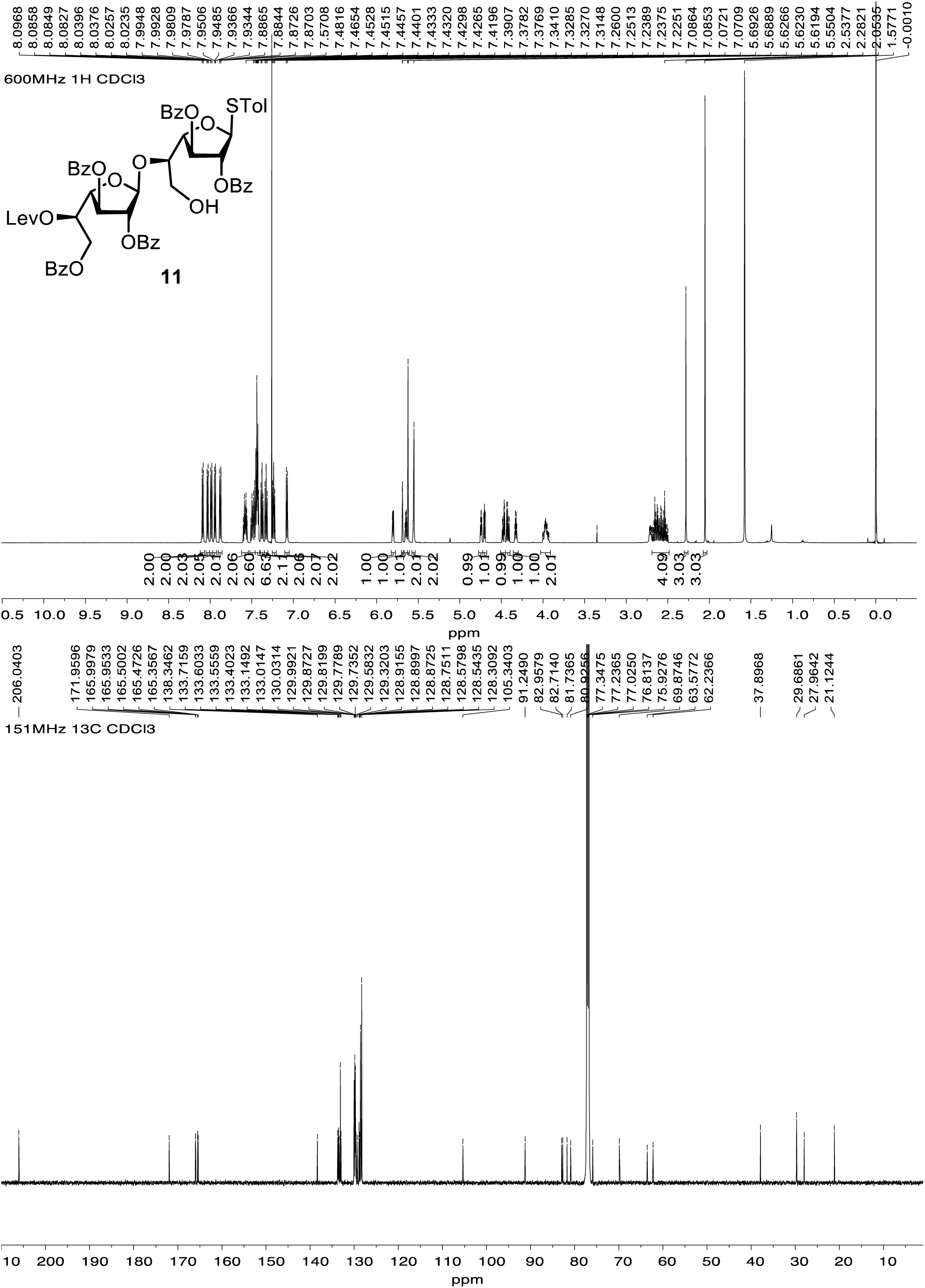

#### 2,3-di-*O*-benzoyl-1-thio-β-D-galactofuranoside (12)

A mixture of 11 (21 mg, 19.7 μmol) and 4 Å molecular sieves (powder, 0. g) in CH_2_Cl_2_ (0.6 mL) was stirred at room temperature for 30 min under an Ar atmosphere and then cooled to –10 °C before bis(cyclopentadienyl)zirconium (IV) dichloride (11 mg, 1.81 mmol) and silver trifluoromethanesulfonate (19 mg, 34.0 μmol) were added successively. After stirring for 10 min at –10 °C, glycosyl fluoride **3** (30 mg, 37.6 μmol) in CH_2_Cl_2_ (0.2 mL) was added dropwise over 30 min. After stirring at –10 °C for 4 h, triethylamine was added and the resulting mixture was filtered through Celite. The filtrate was diluted with CH_2_Cl_2_ (2 mL), washed with satd aq NaHCO_3_ soln and the organic layer was separated and the aqueous layer was extracted with CH_2_Cl_2_ (2mL × 3). The combined organic layers were dried over Na_2_SO_4_, filtered, and the filtrate was concentrated. The residue was purified by chromatography (hexanes–acetone, 5:2) to afford **12 (**27 mg, 87%) as a white foam. *R*_f_ 0.2 (hexanes– acetone, 5:2); [α]_D_ –24.0 (*c* 2.3, CHCl_3_); ^1^H NMR (500 MHz, CDCl_3_, δ_H_) δ 8.16–8.12 (m, 2H, ArH), 8.02–7.94 (m, 4H, ArH), 7.93–7.89 (m, 4H, ArH), 7.88–7.84 (m, 2H, ArH), 7.83–7.78 (m, 2H, ArH), 7.77–7.73 (m, 2H, ArH), 7.64–7.55 (m, 2H, ArH), 7.54–7.40 (m, 15H, ArH), 7.37–7.32 (m, 10H, ArH), 7.27–7.26 (m, 2H, ArH), 7.22–7.19 (m, 2H, ArH), 7.13–7.06 (m, 2H, ArH), 5.95 (app t, *J* = 9.6 Hz, 1H, H-3’’), 5.79–5.75 (m, 1H, H-3), 5.70–5.63 (m, 2H, H-4’’, H-5’), 5.60–5.59 (m, 2H, H-1, H-1’), 5.58–5.56 (m, 2H, H-2’’, H-2), 5.53 (app d, *J* = 1.5 Hz, 1H, H-2’), 5.46 (app d, *J* = 1.5, 4.7 Hz, 1H, H-3’), 5.07 (d, *J* = 7.8 Hz, 1H, H-1’’), 4.76 (app t, *J* = 4.7 Hz, 1H, H-4’), 4.70–4.64 (m, 1H, H-4), 4.59 (dd, *J* = 12.1, 3.2 Hz, 1H, H-6b’’), 4.54 (dd, *J* = 12.1, 3.7 Hz, 1H, H-6b’), 4.51–4.48 (m, 1H, H-5), 4.43 (dd, *J* = 12.0, 5.2 Hz, 1H, H-6a’’), 4.37 (dd, *J* = 12.0, 7.5 Hz, 1H, H-6a’), 4.26–4.20 (m, 2H, H-5’’, H-6b), 4.12 (dd, *J* = 10.9, 6.9 Hz, 1H, H-6a), 2.68–2.35 (m, 4H, Lev CH_2_), 2.30 (s, 3H, ArCH_3_), 2.03 (s, 3H, Lev CH_3_); ^13^C NMR (126 MHz, CDCl_3_, δ_C_) δ 205.9 (Lev ketone C=O), 171.2 (Lev ester C=O), 166.0 (Ar), 165.9 (Ar), 165.7 (Ar), 165.42 (Ar), 165.38 (Ar), 165.3 (Ar), 165.1 (Ar), 164.9 (Ar), 138.1 (Ar), 133.52 (Ar), 133.47 (Ar), 133.4 (Ar), 133.3 (Ar), 133.1 (Ar), 133.0 (Ar), 132.9 (Ar), 130.1 (Ar), 130.0 (Ar), 129.83 (Ar), 129.77 (Ar), 129.73 (Ar), 129.70 (Ar), 129.67 (Ar), 129.6 (Ar), 129.1 (Ar), 129.0 (Ar), 128.9 (Ar), 128.83 (Ar), 128.77 (Ar), 128.53 (Ar), 128.47 (Ar), 128.4 (Ar), 128.33 (Ar), 128.27 (Ar), 128.23 (Ar), 128.18 (Ar), 105.3 (C-1’), 100.6 (C-1’’), 90.9 (C-1), 81.9 (C-4), 81.7 (C-4’), 81.6 (C-2), 81.4 (C-2’), 77.2 (C-3), 77.3 (C-3’), 74.0 (C-5), 72.8 (C-3’’), 72.2 (C-5’’), 71.8 (C-2’’), 70.1 (C-5’), 69.8 (C-4’’), 67.9 (C-6), 63.4 (C-6’), 63.2 (C-6’’), 37.9 (Lev CH_2_), 29.6 (Lev CH_3_), 27.9 (Lev CH_2_), 21.1 (ArCH_3_); HRMS (ESI) Calcd for (M + Na) C_93_H_80_NaO_26_S: 1667.4550. Found 1667.4554.

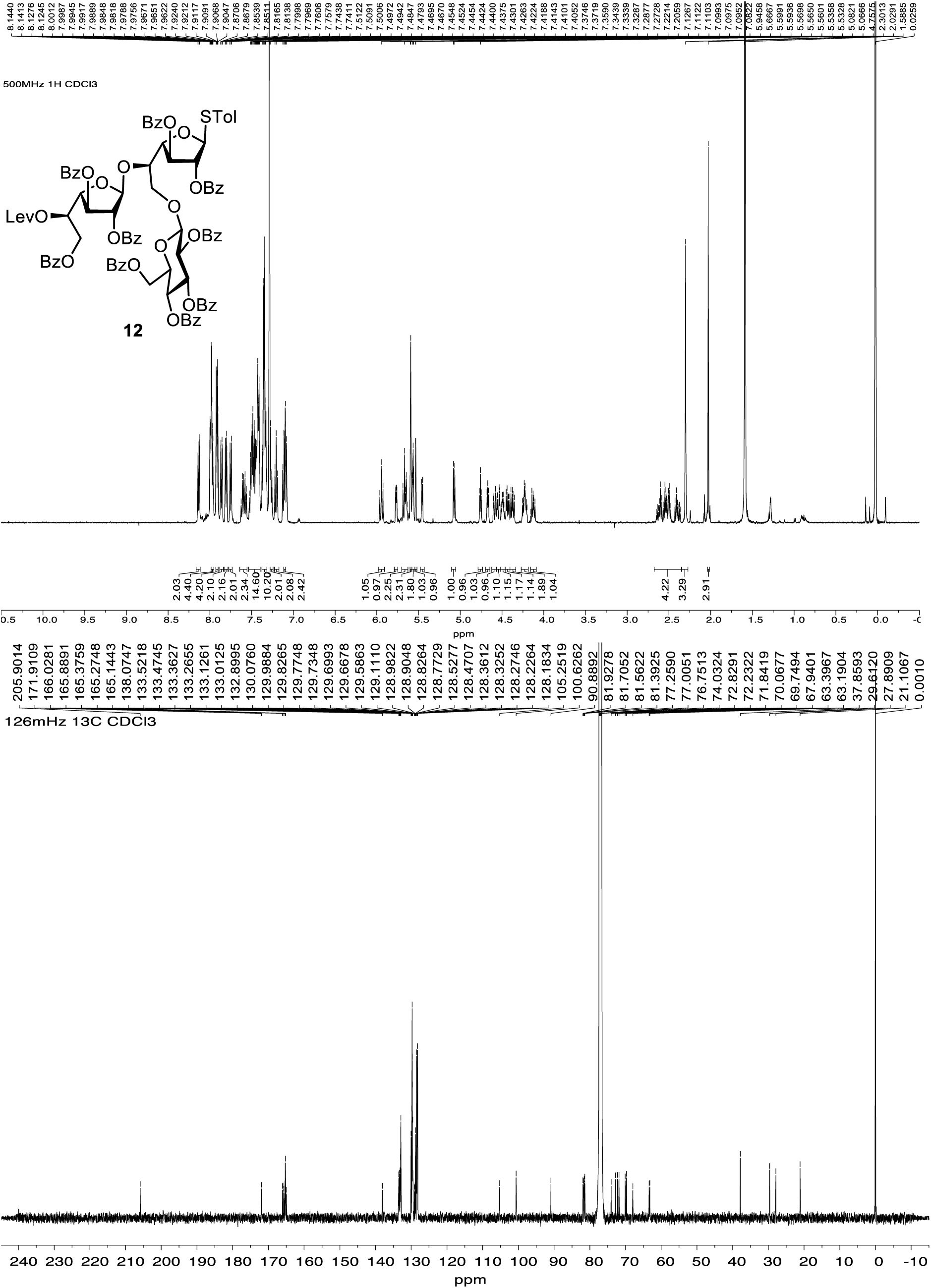

#### 8-Azidooctyl 2,3,6-tri-*O*-benzoyl-5-*O*-levulinoyl-β-D-galactofuranosyl-(1→5)-[2,3,4,6-tetra-*O*-benzoyl-β-D-glucopyranosyl-(1→6)]-2,3-di-*O*-benzoyl-β-D-galactofuranoside (13)

A mixture of 12 (0.15 g, 94 μmol), 8-azido-1-octanol (32 mg, 0.18 mmol) and 4 Å molecular sieves (powder, 1 g) in CH_2_Cl_2_ (3 mL) was stirred at room temperature for 30 min under an Ar atmosphere and then cooled to –10 °C before silver trifluoromethanesulfonate (24 mg, 93 μmol) and *N*-iodosuccinimide (27 mg, 93 μmol) were added successively. After stirring at –10 °C, for 4 h, triethylamine was added and the resulting mixture was filtered through Celite. The filtrate was diluted with CH_2_Cl_2_ (10 mL) and washed with satd aq NaHCO_3_ soln. The organic layer was separated and the aqueous layer was extracted with CH_2_Cl_2_ (10 mL × 3). The combined organic layers were dried over Na_2_SO_4_, filtered, the filtrate was concentrated and the residue was purified by chromatography (hexanes–acetone, 5:2) to afford **13** (0.16 g, 99%) as a white foam. *R*_f_ 0.15 (hexanes–acetone, 5:2); [α]_D_ –20.0 (*c* 0.15, CHCl_3_); ^1^H NMR (500 MHz, CDCl_3_, δ_H_) 8.15–8.09 (m, 2H, ArH), 8.00–7.91 (m, 8H, ArH), 7.91–7.86 (m, 2H, ArH), 7.85–7.80 (m, 2H, ArH), 7.80–7.76 (m, 2H, ArH), 7.76–7.72 (m, 2H, ArH), 7.62–7.57 (m, 1H, ArH), 7.57–7.51 (m, 1H, ArH), 7.51–7.29 (m, 21H, ArH), 7.27–7.18 (m, 2H, ArH), 7.14–7.07 (m, 2H, ArH), 5.93 (app t, *J* = 9.6 Hz, 1H, H-3”), 5.71–5.61 (m, 3H, H-3, H-5’, H-4”), 5.59 (app s, 1H, H-1’), 5.57–5.51 (m, 2H, H-2’, H-2”), 5.42 (dd, *J* = 5.1, 1.5 Hz, 1H, H-3’), 5.37 (app d, *J* = 1.6 Hz, 1H, H-2), 5.09 (app s, 1H, H-1), 5.06 (d, *J* = 7.8 Hz, 1H, H-1”), 4.79 (dd, *J* = 5.1, 4.3 Hz, 1H, H-4’), 4.63 (dd, *J* = 12.1, 3.5 Hz, 1H, H-6b”), 4.56 (dd, *J* = 12.1, 3.3 Hz, 1H, H-6b’), 4.50–4.35 (m, 4H, H-4, H-5, H-6a’, H-6a”), 4.29–4.16 (m, 2H, H-6b, H-5”), 4.10 (dd, *J* = 10.7, 7.0 Hz, 1H, H-6a), 3.63 (dt, *J* = 9.6, 6.6 Hz, 1H, OC<u>H_2_</u>CH_2_), 3.37 (dt, *J* = 9.6, 6.3 Hz, 1H, OC<u>H_2_</u>CH_2_), 3.21 (t, *J* = 7.0 Hz, 2H, C<u>H_2_</u>N_3_), 2.66–2.33 (m, 4H, Lev CH_2_), 2.00 (s, 3H, Lev CH_3_), 1.58–1.48 (m, 4H, octyl CH_2_), 1.34–1.26 (m, 8H, octyl CH_2_); ^13^C NMR (126 MHz, CDCl_3_, δ_C_) 205.9 (Lev ketone C=O), 171.9 (Lev ester C=O), 166.0 (Ar), 165.9 (Ar), 165.7 (Ar), 165.5 (Ar), 165.4 (Ar), 165.3 (Ar), 165.2 (Ar), 164.9 (Ar), 133.5 (Ar), 133.38 (Ar), 133.35 (Ar), 133.2 (Ar), 133.1 (Ar), 133.03 (Ar), 132.97 (Ar), 130.1 (Ar), 129.9 (Ar), 129.82 (Ar), 129.79 (Ar), 129.75 (Ar), 129.71 (Ar), 129.68 (Ar), 129.65 (Ar), 129.6 (Ar), 129.5 (Ar), 129.13 (Ar), 129.10 (Ar), 129.0 (Ar), 128.82 (Ar), 128.80 (Ar), 128.54 (Ar), 128.46 (Ar), 128.39 (Ar), 128.37 (Ar), 128.32 (Ar), 128.29 (Ar), 128.24 (Ar), 128.20 (Ar), 128.16 (Ar), 105.2 (C-1), 105.1 (C-1’), 100.6 (C-1”), 81.9 (C-4), 81.8 (C-4’), 81.6 (C-2), 81.5 (C-2’), 76.9 (C-3, C-3’), 73.7 (C-5), 72.9 (C-3”), 72.2 (C-2”), 71.8 (C-5”), 70.1 (C-5’), 69.8 (C-4”), 68.0 (C-6), 67.3 (O<u>C</u>H_2_CH_2_), 63.4 (C-6’), 63.2 (C-6”), 51.4 (CH_2_N_3_), 37.9 (Lev CH_2_), 29.6 (octyl CH_2_), 29.4 (octyl CH_2_), 29.3 (octyl CH_2_), 29.1 (octyl CH_2_), 28.8 (Lev CH_2_), 27.9 (Lev CH_3_), 26.7 (octyl CH_2_), 26.0 (octyl CH_2_); HRMS (ESI) Calcd for (M + Na) C_94_H_89_N_3_NaO_27_: 1714.5576. Found 1714.5575.

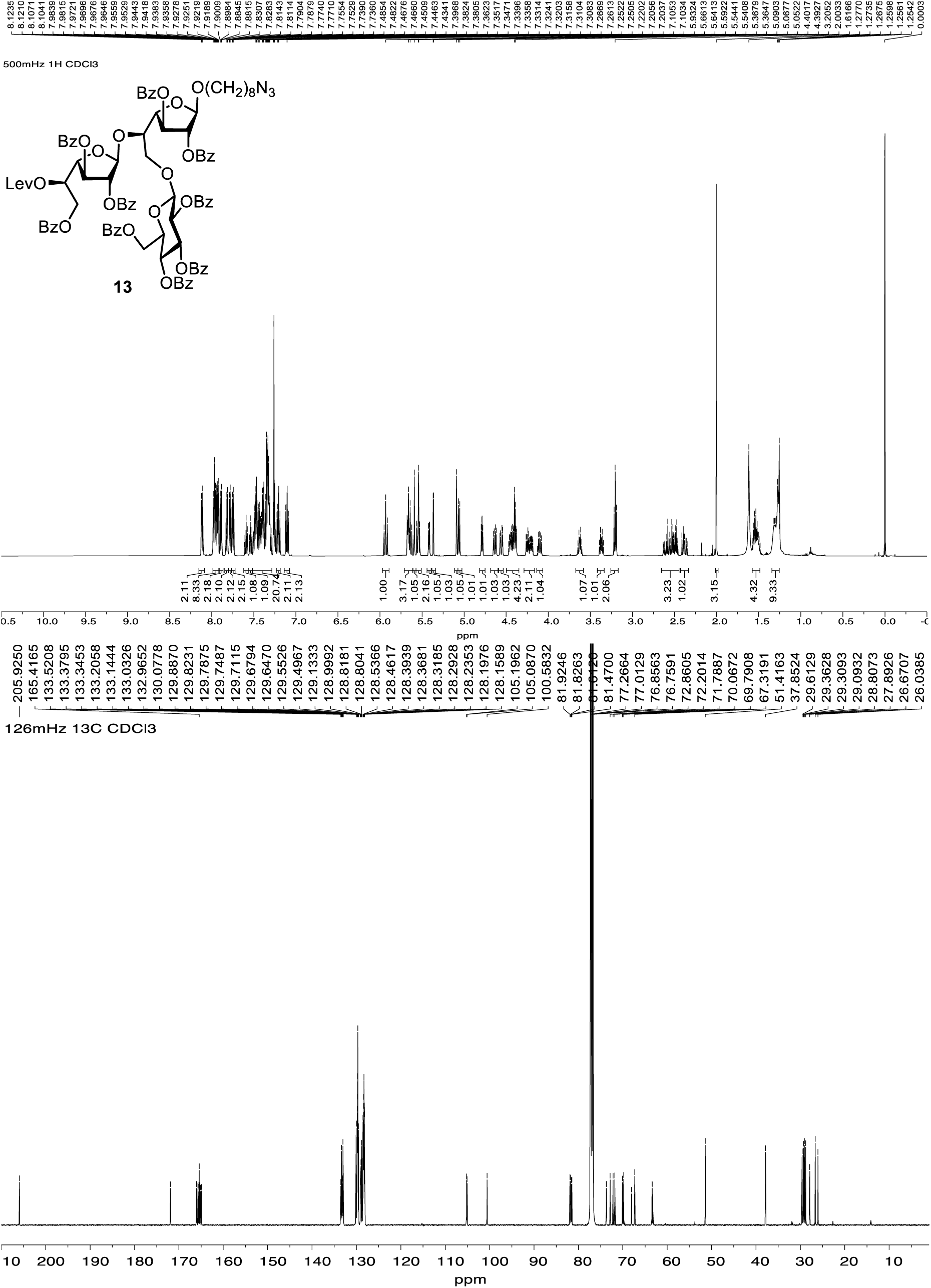

#### 8-Azidooctyl 2,3,6-tri-*O*-benzoyl-β-D-galactofuranosyl-(1→5)-[2,3,4,6-tetra-*O*-benzoyl-β-D-glucopyranosyl-(1→6)]-2,3-di-*O*-benzoyl-β-D-galactofuranoside (14)

To a solution of 13 (150 mg, 88.6 μmol) and hydrazine acetate (13 mg, 14.1 μmol) were dissolved in CH_2_Cl_2_–CH_3_OH (1:2 mL). After stirring at room temperature for 4 h, the mixture was diluted with CH_2_Cl_2_ (10 mL) and washed with satd NaCl soln. The organic layer was separated and the aqueous layer was extracted with CH_2_Cl_2_ (10 mL × 3). The combined organic layers were dried over Na_2_SO_4_, filtered and the filtrate was concentrated. The residue was purified by chromatography (hexanes–acetone, 5:2) to afford **14** (0.12 g, 82%) as a white foam. *R*_f_ 0.23 (hexanes–acetone, 5:2); [α]_D_ –42.9 (*c* 0.07, CHCl_3_); ^1^H NMR (500 MHz, CDCl_3_, δ_H_) 8.12–8.03 (m, 2H, ArH), 8.01–7.97 (m, 4H, ArH), 7.94–7.89 (m, 6H, ArH), 7.82–7.71 (m, 6H, ArH), 7.62–7.57 (m, 1H, ArH), 7.56–7.28 (m, 20H, ArH), 7.25–7.21 (m, 2H, ArH), 7.19–7.16 (m, 2H, ArH), 7.12–7.05 (m, 2H, ArH), 5.94 (app t, *J* = 9.6 Hz, 1H, H-3”), 5.72–5.62 (m, 2H, H-3, H-4”), 5.62–5.56 (m, 2H, H-1’, H-2”), 5.55 (dd, *J* = 1.8, 0.7 Hz, 1H, H-2’), 5.52 (dd, *J* = 5.1, 1.8 Hz, 1H, H-3’), 5.36 (dd, *J* = 1.7, 0.6 Hz, 1H, H-2), 5.11–5.02 (m, 2H, H-1”, H-1), 4.68 (dd, *J* = 5.1, 3.0 Hz, 1H, H-4’), 4.59 (dd, *J* = 12.1, 3.3 Hz, 1H, H-6b), 4.53–4.42 (m, 3H, H-6a”, H-6b”, H-6b’), 4.40–4.33 (m, 2H, H-4, H-5), 4.27–4.24 (m, 2H, H-6a, H-5’), 4.21 (ddd, *J* = 9.1, 5.1, 3.3 Hz, 1H, H-5”), 4.10 (dd, *J* = 10.9, 7.3 Hz, 1H, H-6a’), 3.60 (dt, *J* = 9.5, 6.6 Hz, 1H, OC<u>H_2_</u>CH_2_), 3.36 (dt, *J* = 9.5, 6.3 Hz, 1H, OC<u>H_2_</u>CH_2_), 3.20 (t, *J* = 7.0 Hz, 2H, C<u>H_2_</u>N_3_), 2.63 (d, *J* = 9.0 Hz, 1H, O<u>H</u>), 1.52 (dd, *J* = 14.7, 7.6 Hz, 4H, octyl CH_2_), 1.35–1.22 (m, 8H, octyl CH_2_); ^13^C NMR (126 MHz, CDCl_3_, δ_C_) δ 166.1 (Ar), 165.8 (Ar), 165.6 (Ar), 133.5 (Ar), 133.40 (Ar), 133.35 (Ar), 133.2 (Ar), 133.14 (Ar), 133.07 (Ar), 132.9 (Ar), 130.1 (Ar), 129.9 (Ar), 129.8 (Ar), 129.70 (Ar), 129.68 (Ar), 129.6 (Ar), 129.5 (Ar), 129.1 (Ar), 129.0 (Ar), 128.9 (Ar), 128.8 (Ar), 128.6 (Ar), 128.5 (Ar), 128.38 (Ar), 128.35 (Ar), 128.32 (Ar), 128.26 (Ar), 128.2 (Ar), 128.1 (Ar), 105.3 (C-1), 105.0 (C-1’), 100.6 (C-1”), 83.2 (C-4), 82.1 (C-4’), 82.0 (C-2), 81.8 (C-2’), 78.2 (C-3’), 76.9 (C-3), 73.4 (C-5), 72.8 (C-3”), 72.2 (C-2”), 71.9 (C-5”), 69.8 (C-4”), 69.4 (C-5’), 69.1 (C-6), 67.3 (OC<u>H_2_</u>CH_2_), 66.2 (C-6’), 63.2 (C-6”), 51.4 (<u>C</u>H_2_N_3_), 29.4 (octyl CH_2_), 29.3 (octyl CH_2_), 29.1 (octyl CH_2_), 28.81 (octyl CH_2_), 26.7 (octyl CH_2_), 26.1 (octyl CH_2_); HRMS (ESI) Calcd for (M + Na) C_89_H_83_N_3_NaO_25_:1616.2508 Found 1616.2506.

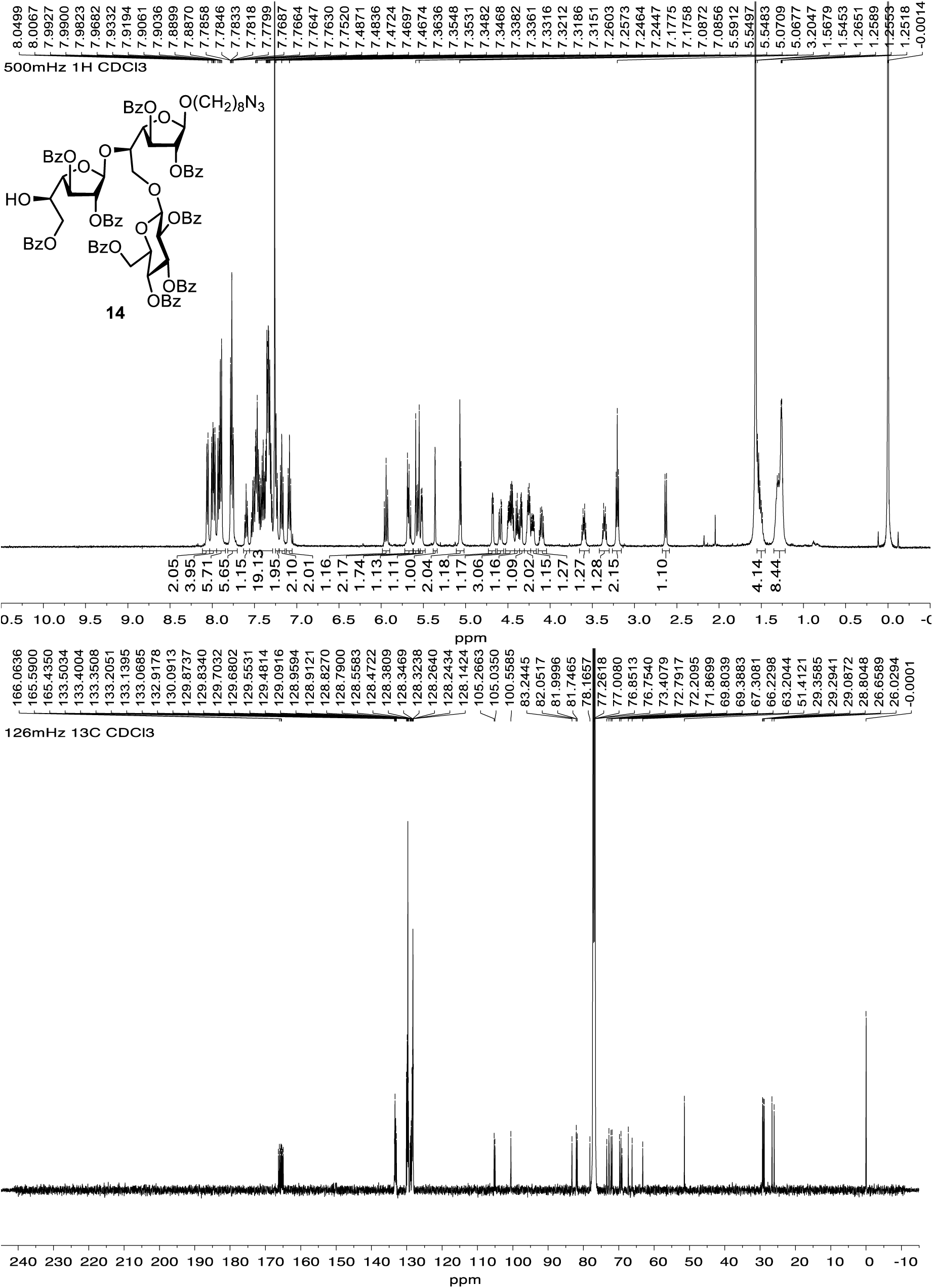

#### 8-Azidooctyl 2,3,6-tri-*O*-benzoyl-5-*O*-levulinoyl-β-D-galactofuranosyl-(1→5)-[2,3,4,6-tetra-*O*-benzoyl-β-D-glucopyranosyl-(1→6)]-2,3-di-*O*-benzoyl-β-D-galactofuranosyl-(1→5)-2,3,6-tri-*O*-benzoyl-β-D-galactofuranosyl-(1→5)-[2,3,4,6-tetra-*O*-benzoyl-β-D-glucopyranosyl-(1→6)]-2,3-di-*O*-benzoyl-β-D-galactofuranoside (15)

A mixture of alcohol **14** (50 mg, 31 μmol), thioglycoside **12** (62 mg, 38 μmol) and 4 Å molecular sieves (powder, 0.33 g) in CH_2_Cl_2_ (1.2 mL) was stirred at room temperature for 30 min and then the mixture as cooled to –5 °C before silver trifluoromethanesulfonate (17 mg, 65 μmol) and *N*-iodosuccinimide (16 mg, 63 μmol) were added successively. After stirring at – 5 °C for 4 h, triethylamine was added and the resulting mixture was filtered through Celite. The filtrate was diluted with CH_2_Cl_2_ (10 mL), washed with satd aq NaHCO_3_ soln and the organic layer was separated. The aqueous layer was extracted with CH_2_Cl_2_ (10 mL × 3) and the combined organic layer was dried over Na_2_SO_4_, filtered and the filtrate was concentrated. The residue was purified by chromatography (hexanes–acetone, 2:1) to afford **15** (65 mg, 73%) as a white foam. *R*_f_ 0.1 (hexanes–acetone, 2:1); [α]_D_ –7.3 (*c* 3.2, CHCl_3_); ^1^H NMR (600 MHz, CDCl_3_, δ_H_) 8.12–8.03 (m, 4H), 7.98–7.91 (m, 6H), 7.91–7.83 (m, 11H), 7.82–7.62 (m, 16H), 7.56 (ddt, *J* = 7.9, 7.0, 1.3 Hz, 1H), 7.50–7.26 (m, 33H), 7.26–7.12 (m, 13H), 7.11–6.99 (m, 6H), 5.91 (app t, *J* = 9.6 Hz, 1H), 5.79 (app t, *J* = 9.6 Hz, 1H), 5.70–5.68 (m, *J* = 5.5 Hz, 2H), 5.67–5.65 (m, 2H), 5.63–5.61 (m, 2H), 5.59–5.58 (m, 3H), 5.55 (s, 1H), 5.54–5.49 (m, 2H), 5.49–5.45 (m, 1H), 5.43 (dd, *J* = 1.7, 0.6 Hz, 1H), 5.37–5.32 (m, 2H), 5.07 (s, 1H), 5.03 (d, *J* = 7.7 Hz, 1H), 4.83–4.81 (m, 2H), 4.67 (dd, *J* = 11.8, 3.8 Hz, 1H), 4.62 (app t, *J* = 4.7 Hz, 1H), 4.59 (dt, *J* = 7.0, 3.4 Hz, 1H), 4.54 (dd, *J* = 6.2, 3.1 Hz, 1H), 4.53–4.47 (m, 3H), 4.46–4.43 (m, 2H), 4.40–4.35 (m, 3H), 4.34–4.27 (m, 2H), 4.24 (dd, *J* = 10.6, 5.6 Hz, 1H), 4.19–4.16 (m, 2H), 4.10–4.03 (m, 2H), 3.95–3.92 (m, 1H), 3.60 (dt, *J* = 9.6, 6.6 Hz, 1H, OC<u>H_2_</u>CH_2_), 3.34 (dt, *J* = 9.6, 6.3 Hz, 1H, OC<u>H_2_</u>CH_2_), 3.19 (t, *J* = 7.0 Hz, 2H, C<u>H_2_</u>N_3_), 2.53–2.11 (m, 4H, Lev CH_2_), 1.93 (s, 3H, Lev CH_3_), 1.56–1.43 (m, 4H, octyl CH_2_), 1.34–1.21 (m, 8H, octyl CH_2_); ^13^C NMR (151 MHz, CDCl_3_, δ_C_) 205.9, 166.0, 165.9, 165.8, 165.7, 165.5, 165.4, 165.3, 165.1, 164.9, 164.8, 133.3, 133.2, 132.9, 132.8, 130.1, 130.0, 129.82, 129.76, 129.71, 129.69, 129.66, 129.60, 129.55, 129.5, 129.1, 128.98, 128.95, 128.92, 128.86, 128.85, 128.8, 128.7, 128.51, 128.46, 128.4, 128.30, 128.28, 128.21, 128.17, 128.11, 128.07, 105.8, 105.1, 105.0, 101.1, 100.5, 83.2, 82.1, 82.0, 81.8, 81.5, 77.8, 73.5, 73.0, 72.9, 72.2, 72.0, 71.8, 71.3, 70.1, 69.9, 69.5, 67.2, 65.0, 63.6, 63.3, 62.7, 51.4, 37.8, 29.6, 29.3, 29.1, 28.8, 27.8, 26.7, 26.0; HRMS (ESI) Calcd for (M + Na) C_175_H_155_N_3_NaO_51_: 3139.1428. Found 3139.1471.

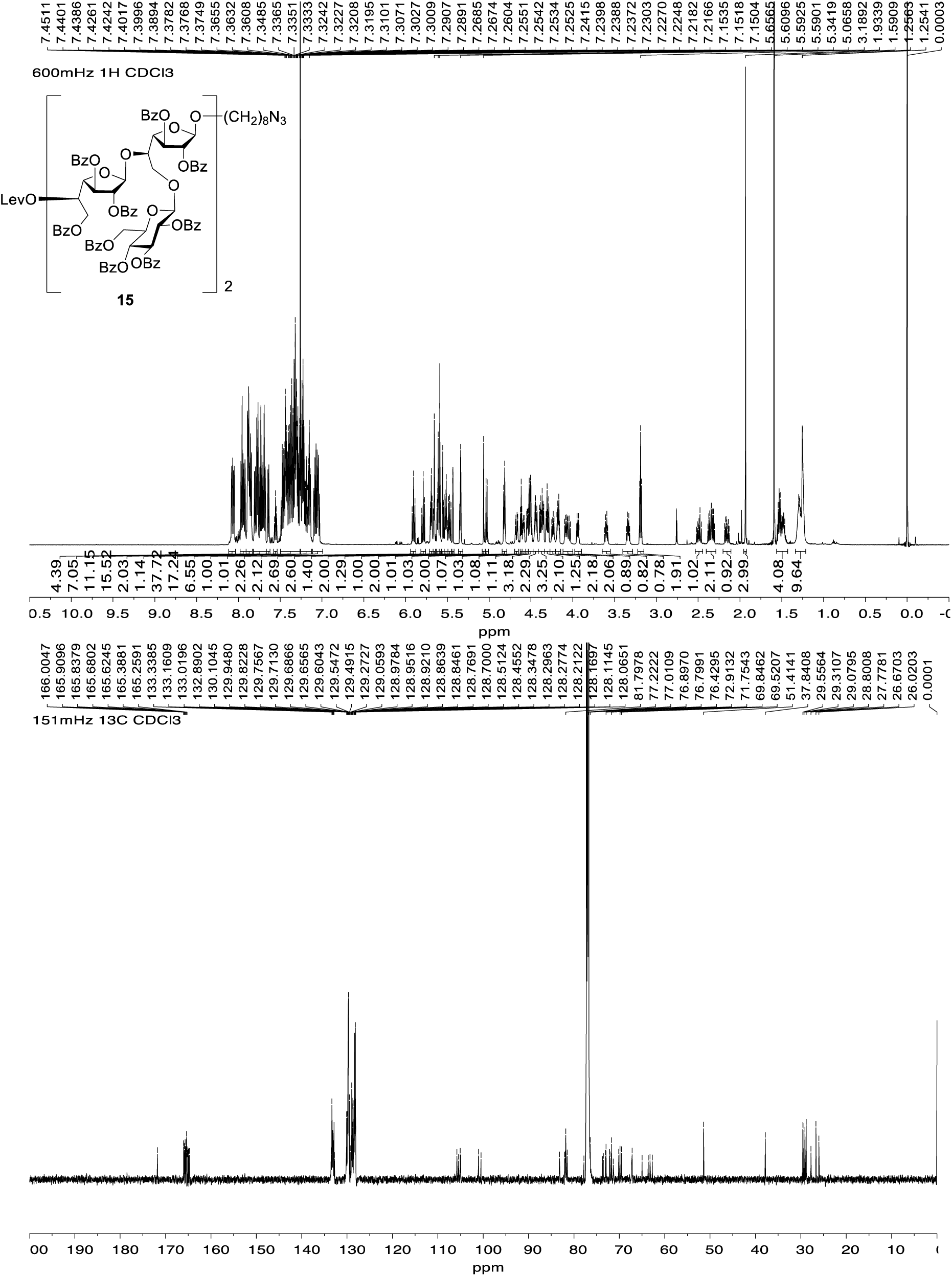

#### 8-Azidooctyl 2,3,6-tri-*O*-benzoyl-β-D-galactofuranosyl-(1→5)-[2,3,4,6-tetra-*O*-benzoyl-β-D-glucopyranosyl-(1→6)]-2,3-di-*O*-benzoyl-β-D-galactofuranosyl-(1→5)-2,3,6-tri-*O*-benzoyl-β-D-galactofuranosyl-(1→5)-[2,3,4,6-tetra-*O*-benzoyl-β-D-glucopyranosyl-(1→6)]-2,3-di-*O*-benzoyl-β-D-galactofuranoside (16)

Hexasaccharide **15** (200 mg, 64 μmol) and hydrazine acetate (12 mg, 0.12 mmol) were dissolved in CH_2_Cl_2_–CH_3_OH (0.7:1.4 mL). After stirring at room temperature for 4 h, the mixture was diluted with CH_2_Cl_2_ (10 mL) and washed with a satd NaCl aq soln. The organic layer was separated and the aqueous layer was extracted with CH_2_Cl_2_ (10 mL × 3). The combined organic layers were dried over Na_2_SO_4_, filtered, and the filtrate was concentrated. The residue was purified by chromatography (hexanes–EtOAc, 1:2) to afford **16** (0.16 g, 83%) as a white foam. *R*_f_ 0.49 (hexanes–EtOAc, 1:1); [α]_D_ –12.7 (*c* 0.71, CHCl_3_); ^1^H NMR (500 MHz, CDCl_3_, δ_H_) δ 8.07–8.02 (m, 4H), 7.99–7.95 (m, 8H), 7.92–7.87 (m, 8H), 7.84–7.78 (m, 6H), 7.77–7.70 (m, 8H), 7.68–7.65 (m, 2H), 7.62–7.57 (m, 1H), 7.53–7.48 (m, 4H), 7.48–7.40 (m, 10H), 7.40–7.31 (m, 17H), 7.28–7.24 (m, 9H), 7.24–7.04 (m, 13H), 5.93 (app t, *J* = 9.6 Hz, 1H), 5.86 (app t, *J* = 9.6 Hz, 1H), 5.73–5.71 (m, 3H), 5.68 (d, *J* = 9.7 Hz, 1H), 5.65–5.63 (m, 3H), 5.62–5.59 (m, 2H), 5.58–5.51 (m, 3H), 5.48–5.47 (m, 2H), 5.36 (d, *J* = 1.4 Hz, 1H), 5.09 (s, 1H), 5.05 (d, *J* = 7.8 Hz, 1H), 4.88 (d, *J* = 7.8 Hz, 1H), 4.83 (dd, *J* = 4.6, 3.1 Hz, 1H), 4.74–4.67 (m, 1H), 4.62–4.52 (m, 5H), 4.51–4.44 (m, 2H), 4.44–4.16 (m, 9H), 4.16–4.03 (m, 3H), 4.00 (dt, *J* = 9.9, 3.6 Hz, 1H), 3.67–3.58 (m, 1H, OC<u>H_2_</u>CH_2_), 3.39–3.34 (m, 1H, OC<u>H_2_</u>CH_2_), 3.22 (t, *J* = 7.0 Hz, 2H, <u>CH_2_</u>N_3_), 2.68 (d, *J* = 8.9 Hz, 1H, OH), 1.58–1.45 (m, 4H, octyl CH_2_), 1.33–1.26 (m, 8H, octyl CH_2_); ^13^C NMR (126 MHz, CDCl_3_, δ_C_) 166.0, 165.9, 165.8, 165.7, 165.2, 164.9, 133.3, 133.2, 133.1, 132.9, 130.1, 130.0, 129.84, 129.77, 129.7, 129.6, 129.5, 129.4, 129.2, 129.1, 129.0, 128.9, 128.7, 128.5, 128.4, 128.3, 128.21, 128.17, 128.1, 105.5, 105.0, 101.0, 100.5, 83.4, 83.1, 82.6, 82.0, 81.9, 81.81, 81.77, 81.7, 77.6, 77.5, 73.0, 72.9, 72.2, 72.0, 71.80, 71.76, 69.9, 69.6, 69.5, 67.4, 67.2, 66.3, 65.1, 63.3, 62.8, 62.7, 51.4, 29.3, 29.1, 28.8, 26.7, 26.0; HRMS (ESI) Calcd for (M + Na) C_170_H_149_O_49_N_3_Na: 3038.9157. Found 3038.9152.

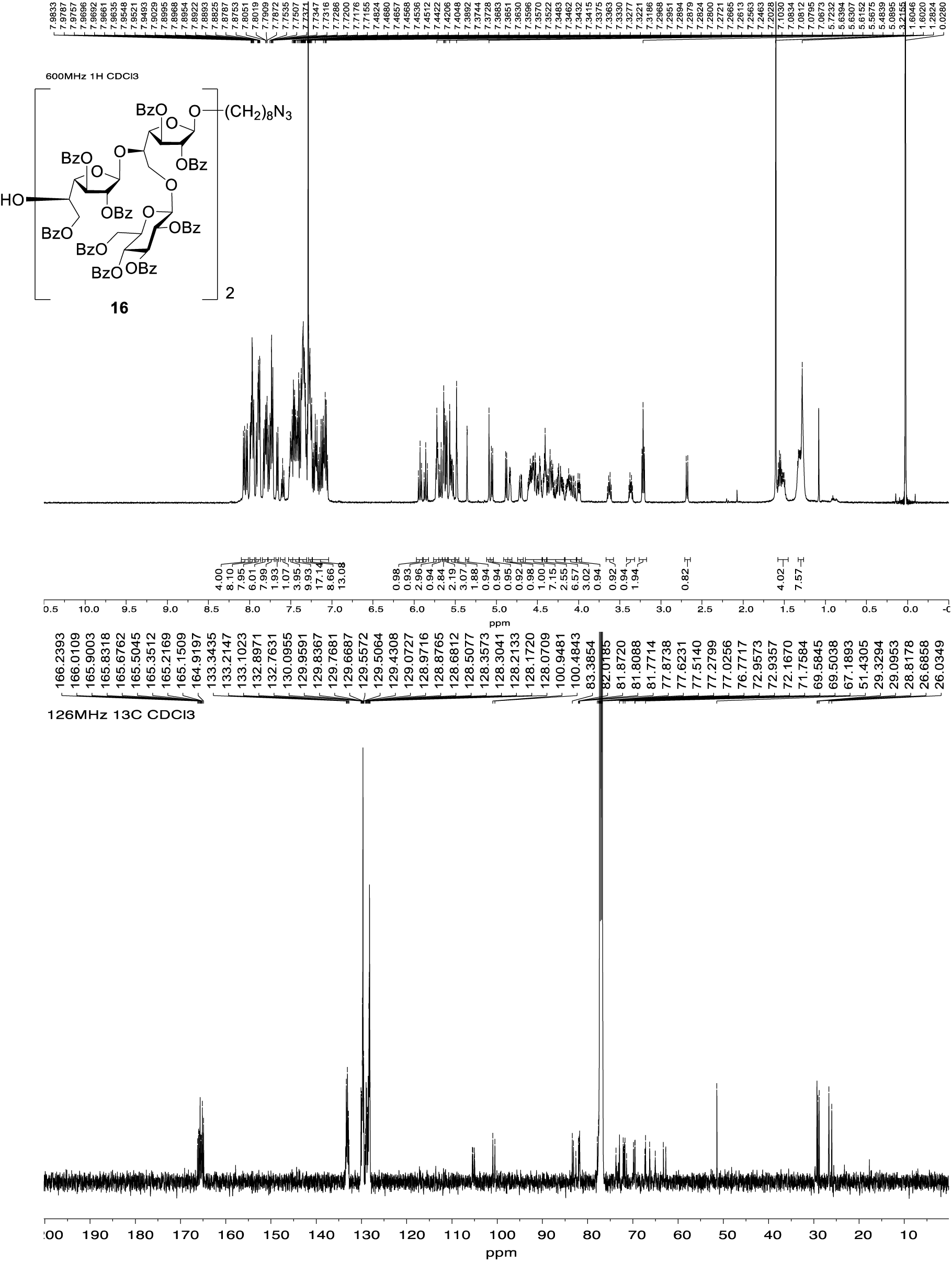

#### 8-Azidooctyl 2,3,6-tri-*O*-benzoyl-5-*O*-levulinoyl-β-D-galactofuranosyl-(1→5)-[2,3,4,6-tetra-*O*-benzoyl-β-D-glucopyranosyl-(1→6)]-2,3-di-*O*-benzoyl-β-D-galactofuranosyl-(1→5)-2,3,6-tri-*O*-benzoyl-β-D-galactofuranosyl-(1→5)-[2,3,4,6-tetra-*O*-benzoyl-β-D-glucopyranosyl-(1→6)]-2,3-di-*O*-benzoyl-β-D-galactofuranosyl-(1→5)-2,3,6-tri-*O*-benzoyl-β-D-galactofuranosyl-(1→5)-[2,3,4,6-tetra-*O*-benzoyl-β-D-glucopyranosyl-(1→6)]-2,3-di-*O*-benzoyl-β-D-galactofuranoside (17)

A mixture of alcohol **16** (69 mg, 23 μmol), thioglycoside **12** (57 mg, 35 μmol) and 4 Å molecular sieves (powder, 0.25 g) in CH_2_Cl_2_ (0.8 mL) was stirred at room temperature for 30 min and then cooled to –5 °C before silver trifluoromethanesulfonate (10 mg, 32 μmol) and *N*-iodosuccinimide (8 mg, 32 μmol) were added successively. After stirring at –5 °C for 4 h, triethylamine was added and the resulting mixture was filtered through Celite. The filtrate was diluted with CH_2_Cl_2_ (10 mL), washed with satd NaHCO_3_ aq soln and the organic layer was separated. The aqueous layer was extracted with CH_2_Cl_2_ (10 mL × 3) and the combined organic layers were dried over Na_2_SO_4_, filtered and concentrated. The residue was purified by chromatography (hexanes–acetone, 2:3) to afford **17** (88 mg, 85%) as a white foam. *R*_f_ 0.15 (hexanes–acetone, 2:3); [α]_D_ –11.3 (*c* 1.6, CHCl_3_); ^1^H NMR (500 MHz, CDCl_3_, δ_H_) δ 8.12–8.04 (m, 6H), 8.00–7.97 (m, 4H), 7.96–7.89 (m, 10H), 7.89–7.85 (m, 8H), 7.84–7.77 (m, 12H), 7.76–7.71 (m, 11H), 7.69–7.64 (m, 3H), 7.63–7.60 (m, 2H), 7.51–7.41 (m, 15H), 7.41–7.38 (m, 9H), 7.37–7.30 (m, 24H), 7.26–7.17 (m, 18H), 7.12–7.05 (m, 13H), 5.93 (app t, *J* = 9.5 Hz, 1H), 5.87–5.79 (m, 2H), 5.79–5.74 (m, 3H), 5.71 (s, 2H), 5.69–5.45 (m, 14H), 5.43 (d, *J* = 1.8 Hz, 1H), 5.39–5.33 (m, 2H), 5.09 (s, 1H), 5.05 (d, *J* = 7.8 Hz, 1H), 4.89 (d, *J* = 7.8 Hz, 1H), 4.82–4.81 (m, 2H), 4.72 (d, *J* = 4.9 Hz, 1H), 4.70–4.57 (m, 6H), 4.57–4.38 (m, 11H), 4.38–4.16 (m, 9H), 4.13–4.05 (m, 3H), 4.03–3.99 (m, 2H), 3.97–3.91 (m, 1H), 3.65–3.58 (m, 1H, OC<u>H_2_</u>CH_2_), 3.36–3.31 (m, 1H, OC<u>H_2_</u>CH_2_), 3.21 (t, *J* = 7.0 Hz, 2H, <u>CH_2_</u>N_3_), 2.57–2.46 (m, 2H, Lev CH_2_), 2.41–2.31 (m, 2H, Lev CH_2_), 1.95 (s, 3H, Lev CH_3_), 1.54–1.50 (m, 4H, octyl CH_2_), 1.28–1.27 (m, 8H, octyl CH_2_); ^13^C NMR (126 MHz, CDCl_3_, δ_C_) 206.0, 172.0, 171.9, 171.84, 171.78, 166.1, 166.0, 165.92, 165.86, 165.82, 165.79, 165.68, 165.67, 165.6, 165.54, 165.47, 165.4, 165.43, 165.42, 165.3, 165.2, 165.14, 165.09, 165.02, 164.98, 164.9, 164.8, 164.7, 133.3, 133.0, 132.9, 130.1, 130.03, 129.95, 129.8, 129.74, 129.66, 129.6, 129.5, 129.3, 129.0, 128.9, 128.7, 128.6, 128.5, 128.4, 128.3, 128.22, 128.17, 128.1, 105.8, 105.7, 105.5, 105.4, 105.0, 104.9, 101.13, 101.06, 100.5, 92.3, 83.2, 81.9, 81.8, 81.72, 81.69, 81.66, 81.6, 73.7, 73.5, 73.4, 73.1, 73.0, 72.9, 72.2, 72.14, 72.05, 71.9, 71.8, 71.7, 70.20, 70.18, 70.11, 70.07, 69.9, 69.7, 69.52, 69.46, 67.2, 63.6, 63.5, 63.3, 63.17, 63.15, 63.0, 62.92, 62.88, 62.8, 62.67, 62.66, 36.6, 29.7, 29.4, 29.3113, 29.3106, 29.3, 28.4, 26.7, 26.0, 24.7, 23.3; HRMS (ESI) Calcd for (M + Na) C_256_H_221_O_75_N_3_Na: 4559.3469. Found 4559.5341.

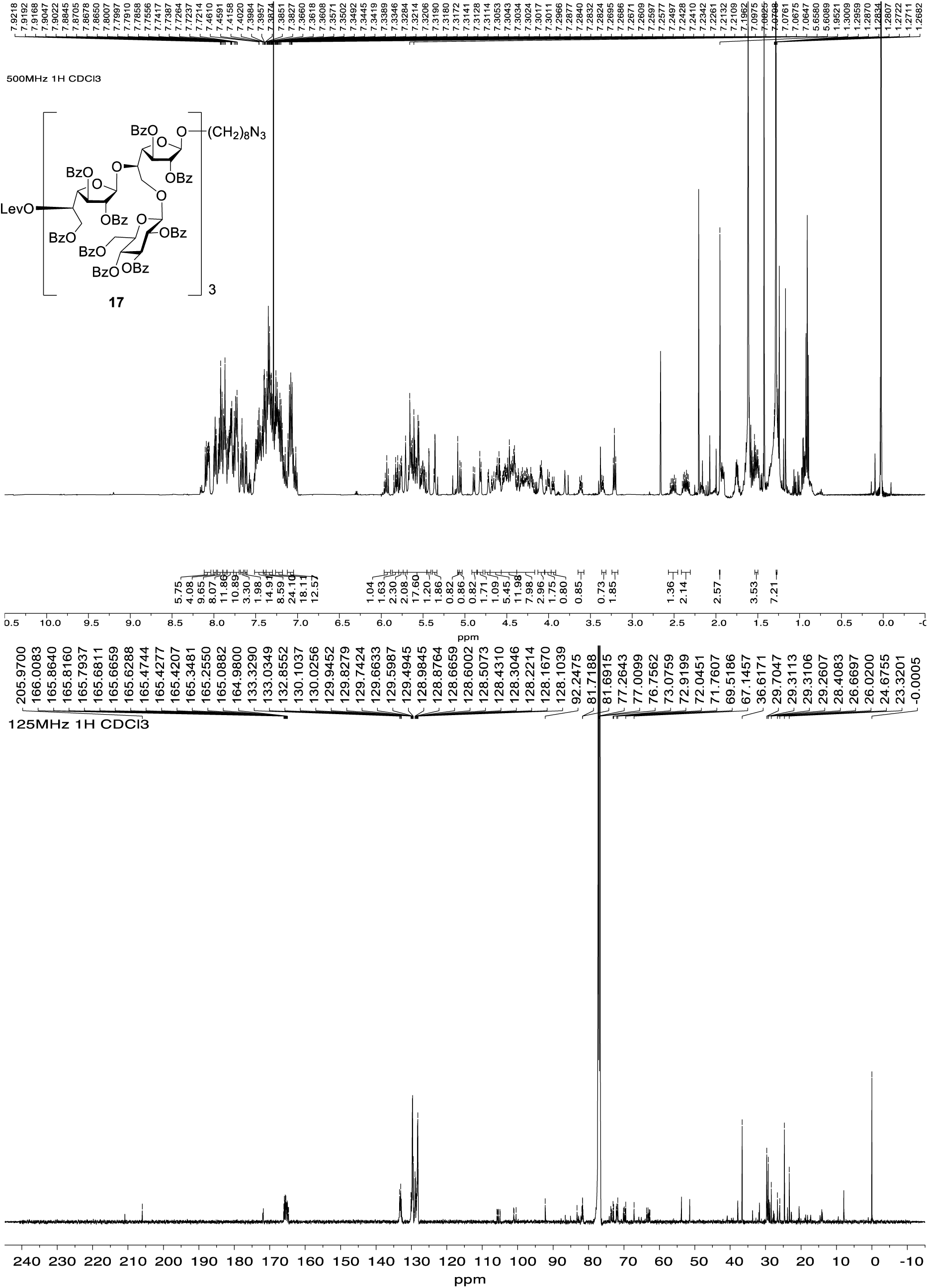

#### 8-Azidooctyl 2,3,6-tri-*O*-benzoyl-β-D-galactofuranosyl-(1→5)-[2,3,4,6-tetra-*O*-benzoyl-β-D-glucopyranosyl-(1→6)]-2,3-di-*O*-benzoyl-β-D-galactofuranosyl-(1→5)-2,3,6-tri-*O*-benzoyl-β-D-galactofuranosyl-(1→5)-[2,3,4,6-tetra-*O*-benzoyl-β-D-glucopyranosyl-(1→6)]-2,3-di-*O*-benzoyl-β-D-galactofuranosyl-(1→5)-2,3,6-tri-*O*-benzoyl-β-D-galactofuranosyl-(1→5)-[2,3,4,6-tetra-*O*-benzoyl-β-D-glucopyranosyl-(1→6)]-2,3-di-*O*-benzoyl-β-D-galactofuranoside (18)

Nonasaccharide **17** (88 mg, 19 μmol) and hydrazine acetate (4 mg, 77 μmol) were dissolved in CH_2_Cl_2_–CH_3_OH (0.7:1.4 mL). After stirring at room temperature for 4 h, the mixture was diluted with CH_2_Cl_2_ (10 mL), and washed with a satd aq NaCl soln. The organic layer was separated and the aqueous layer was extracted with CH_2_Cl_2_ (10 mL × 3). The combined organic layers were dried over Na_2_SO_4_, filtered and concentrated. The residue was purified by chromatography (hexanes–EtOAc, 2:3) to afford **18** (80 mg, 78%) as a white foam. *R*_f_ 0.24 (hexanes–EtOAc, 1:1); [α]_D_ –14.3 (*c* 0.21, CHCl_3_); ^1^H NMR (600 MHz, CDCl_3_, δ_H_) 8.08–8.01 (m, 6H), 7.99–7.96 (m, 4H), 7.96–7.92 (m, 4H), 7.91–7.89 (m, 4H), 7.89–7.84 (m, 9H), 7.84–7.82 (m, 2H), 7.82–7.77 (m, 6H), 7.76–7.75 (m, 2H), 7.75–7.70 (m, 10H), 7.70–7.65 (m, 4H), 7.63–7.60 (m, 2H), 7.60–7.55 (m, 1H), 7.51–7.40 (m, 12H), 7.40–7.36 (m, 7H), 7.36–7.26 (m, 26H), 7.26–7.18 (m, 16H), 7.18–7.11 (m, 7H), 7.11–7.03 (m, 11H), 7.03–6.98 (m, 2H), 5.93 (app t, *J* = 9.5 Hz, 1H), 5.86–5.81 (m, 2H), 5.71–5.62 (m, 2H), 5.72–5.62 (m, 7H), 5.62–5.58 (m, 4H), 5.57–5.53 (m, 5H), 5.53–5.43 (m, 4H), 5.35 (d, *J* = 1.5 Hz, 1H), 5.08 (s, 1H), 5.05 (d, *J* = 7.8 Hz, 1H), 4.88 (d, *J* = 7.9 Hz, 1H), 4.85 (d, *J* = 7.8 Hz, 1H), 4.81 (dd, *J* = 4.9, 2.7 Hz, 1H), 4.70 (dd, *J* = 4.8, 2.4 Hz, 1H), 4.67 (dd, *J* = 11.8, 3.6 Hz, 1H), 4.62–4.59 (m, 3H), 4.56–4.38 (m, 13H), 4.34–4.23 (m, 5H), 4.22–4.18 (m, 2H), 4.10–4.05 (m, 4H), 4.01–3.95 (m, 3H), 3.61 (dt, *J* = 9.6, 6.6 Hz, 1H, OC<u>H_2_</u>CH_2_), 3.35 (dt, *J* = 9.6, 5.9 Hz, 1H, OC<u>H_2_</u>CH_2_), 3.20 (t, *J* = 7.0 Hz, 2H, C<u>H_2_</u>N_3_), 1.58–1.46 (m, 5H, OH, octyl CH_2_), 1.31–1.27 (m, 8H, octyl CH_2_); ^13^C NMR (151 MHz, CDCl_3_, δ_C_) 166.0, 165.9, 165.69, 165.66, 165.51, 165.45, 165.23, 165.18, 165.1, 165.0, 164.7, 133.4, 133.32, 133.27, 133.1, 132.9, 132.8, 130.11, 130.05, 130.0, 129.88, 129.85, 129.8, 129.74, 129.71, 129.69, 129.64, 129.58, 129.55, 129.53, 129.45, 129.3, 129.14, 129.10, 129.03, 129.00, 128.95, 128.91, 128.88, 128.73, 128.71, 128.69, 128.59, 128.55, 128.5, 128.44, 128.38, 128.36, 128.33, 128.29, 128.25, 128.23, 128.19, 128.15, 128.1, 105.7, 105.6, 105.5, 105.1, 105.0, 101.1, 101.0, 100.5, 83.3, 83.2, 82.3, 82.2, 81.9, 81.7, 81.6, 77.7, 73.9, 73.8, 73.6, 73.5, 73.1, 73.00, 72.96, 72.2, 72.1, 72.0, 71.8, 70.8, 69.9, 69.6, 67.2, 66.3, 65.8, 65.1, 63.3, 62.9, 62.7, 51.4, 29.7, 29.4, 29.1, 28.8, 26.7, 26.1; HRMS (ESI) Calcd for (M + Na) C_251_H_215_O_73_N_3_Na: 4464.4453. Found: 4463.4397.

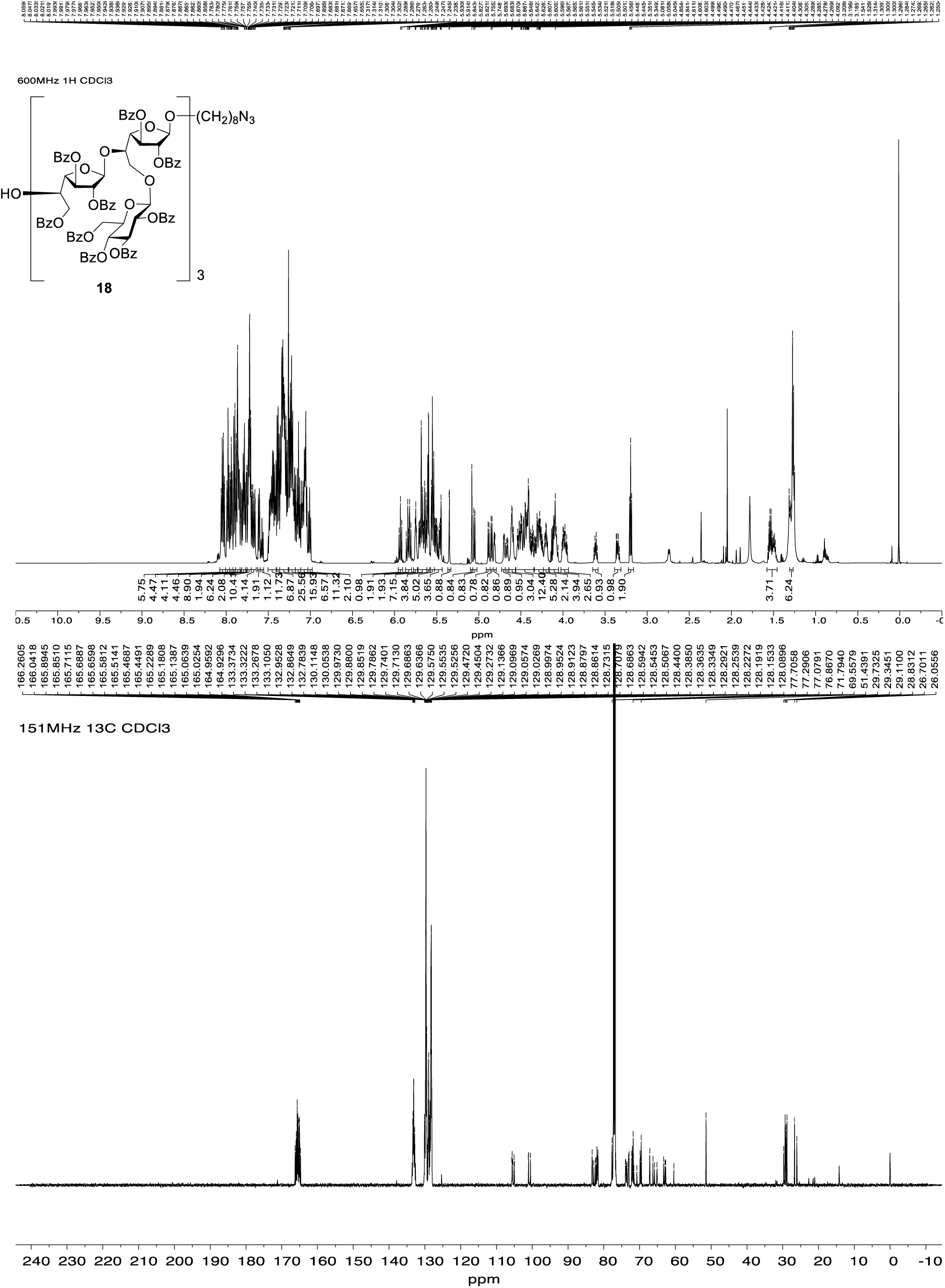

#### 8-Azidooctyl 2,3,6-tri-*O*-benzoyl-5-*O*-levulinoyl-β-D-galactofuranosyl-(1→5)-[2,3,4,6-tetra-*O*-benzoyl-β-D-glucopyranosyl-(1→6)]-2,3-di-*O*-benzoyl-β-D-galactofuranosyl-(1→5)-2,3,6-tri-*O*-benzoyl-β-D-galactofuranosyl-(1→5)-[2,3,4,6-tetra-*O*-benzoyl-β-D-glucopyranosyl-(1→6)]-2,3-di-*O*-benzoyl-β-D-galactofuranosyl-(1→5)-2,3,6-tri-*O*-benzoyl-β-D-galactofuranosyl-(1→5)-[2,3,4,6-tetra-*O*-benzoyl-β-D-glucopyranosyl-(1→6)]-2,3-di-*O*-benzoyl-β-D-galactofuranosyl-(1→5)-2,3,6-tri-*O*-benzoyl-β-D-galactofuranosyl-(1→5)-[2,3,4,6-tetra-*O*-benzoyl-β-D-glucopyranosyl-(1→6)]-2,3-di-*O*-benzoyl-β-D-galactofuranoside (19)

A mixture of alcohol **18** (71 mg, 16 μmol), glycosyl fluoride **24** (86 mg, 56 μmol) and 4 Å molecular sieves (powder, 0.25 g) in CH_2_Cl_2_ (0.8 mL) was stirred at room temperature for 30 min and then cooled to 0 °C before bis(cyclopentadienyl)zirconium (IV) dichloride (16 mg, 55 μmol) and silver trifluoromethanesulfonate (28 mg, 11 μmol) were added successively. After stirring for 4 h at rt, triethylamine was added and the resulting mixture was filtered through Celite. The filtrate was diluted with CH_2_Cl_2_ (10 mL) and washed with satd NaHCO_3_ soln. The organic layer was separated and the aqueous layer was extracted with CH_2_Cl_2_ (10 mL × 3). The combined organic layer was dried over Na_2_SO_4_, filtered and the filtrate was concentrated. The residue was purified by chromatography (hexanes–acetone, 2:3) to afford **19** (81 mg, 85%) as a white foam. *R*_f_ 0.1 (hexanes– acetone, 2:3); [α]_D_ –18.8 (*c* 0.48, CHCl_3_); ^1^H NMR (600 MHz, CDCl_3_, δ_H_) 8.11–8.02 (m, 8H), 8.02–7.68 (m, 56H), 7.68–7.63 (m, 4H), 7.63–7.58 (m, 4H), 7.58–7.53 (m, 1H), 7.50–7.27 (m, 52H), 7.25–7.15 (m, 31H), 7.15–6.96 (m, 24H), 5.96–5.92 (m, 1H), 5.86–5.78 (m, 3H), 5.78–5.57 (m, 18H), 5.57–5.45 (m, 11H), 5.45–5.41 (m, 1H), 5.39–5.34 (m, 2H), 5.09 (s, 1H), 5.06 (d, *J* = 8.6 Hz, 1H), 4.91 (d, *J* = 7.8 Hz, 1H), 4.85 (d, *J* = 7.7 Hz, 1H), 4.83–4.81 (m, 2H), 4.71–4.66 (m, 3H), 4.65–4.17 (m, 29H), 4.12–4.07 (m, 5H), 4.06–3.98 (m, 3H), 4.04–3.99 (m, 2H), 3.64–3.58 (m, 1H, OC<u>H_2_</u>CH_2_), 3.35 (d, *J* = 8.5 Hz, 1H, OC<u>H_2_</u>CH_2_), 3.20 (t, *J* = 7.0 Hz, 2H, C<u>H_2_</u>N_3_), 2.58–2.22 (m, 4H, Lev CH_2_), 1.94 (s, 3H, Lev CH_3_), 1.55–1.48 (m, 4H, octyl CH_2_), 1.33–1.28 (m, 8H, octyl CH_2_); ^13^C NMR (151 MHz, CDCl_3_, δ_C_) δ 206.0, 171.2, 165.91, 165.87, 165.68, 165.66, 165.50, 165.45, 165.3, 165.23, 165.16, 165.1, 165.02, 164.97, 164.8, 164.7, 133.4, 133.3, 133.1, 132.9, 132.7, 130.14, 130.07, 130.0, 129.9, 129.8, 129.7, 129.64, 129.58, 129.53, 129.46, 129.2, 129.14, 129.10, 129.02, 128.99, 128.93, 128.91, 128.88, 128.8, 128.7, 128.63, 128.55, 128.5, 128.4, 128.34, 128.30, 128.26, 128.2, 128.14, 128.09, 105.9, 105.8, 105.7, 105.5, 105.1, 105.0, 101.2, 101.1, 100.5, 83.3, 82.8, 82.6, 82.2, 82.0, 81.9, 81.7, 81.5, 77.9, 77.7, 76.5, 74.3, 73.7, 73.6, 73.1, 73.0, 72.2, 72.1, 71.8, 70.2, 69.9, 69.8, 69.7, 69.5, 67.2, 65.9, 65.1, 63.7, 63.3, 62.8, 62.7, 51.4, 29.7, 29.6, 29.34, 29.29, 29.1, 28.8, 27.8, 26.1; HRMS (ESI) Calcd for (M + Na) C_337_H_287_O_99_N_3_Na:5985.9507. Found 5985.9546.

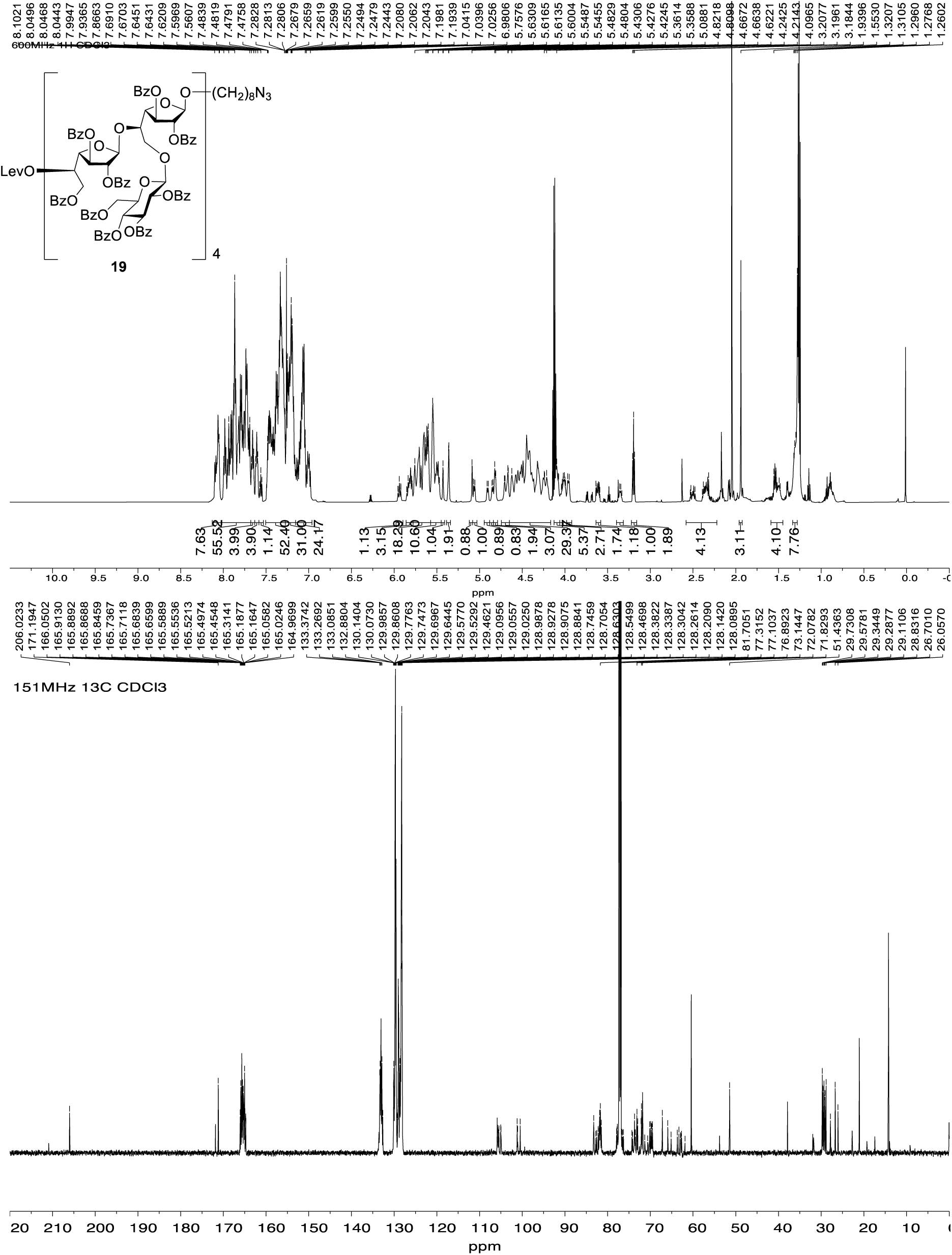

#### 8-Azidooctyl 2,3,6-tri-*O*-benzoyl-β-D-galactofuranosyl-(1→5)-[2,3,4,6-tetra-*O*-benzoyl-β-D-glucopyranosyl-(1→6)]-2,3-di-*O*-benzoyl-β-D-galactofuranosyl-(1→5)-2,3,6-tri-*O*-benzoyl-β-D-galactofuranosyl-(1→5)-[2,3,4,6-tetra-*O*-benzoyl-β-D-glucopyranosyl-(1→6)]-2,3-di-*O*-benzoyl-β-D-galactofuranosyl-(1→5)-2,3,6-tri-*O*-benzoyl-β-D-galactofuranosyl-(1→5)-[2,3,4,6-tetra-*O*-benzoyl-β-D-glucopyranosyl-(1→6)]-2,3-di-*O*-benzoyl-β-D-galactofuranosyl-(1→5)-2,3,6-tri-*O*-benzoyl-β-D-galactofuranosyl-(1→5)-[2,3,4,6-tetra-*O*-benzoyl-β-D-glucopyranosyl-(1→6)]-2,3-di-*O*-benzoyl-β-D-galactofuranoside (20)

Dodecasaccharide **19** (60 mg, 10 μmol) and hydrazine acetate (1 mg, 11 μmol) were dissolved in CH_2_Cl_2_–CH_3_OH (0.9:0.3 mL). After stirring at room temperature for 4 h, the resulting mixture was diluted with CH_2_Cl_2_ (10 mL) and washed with satd NaCl solution. The organic layer was separated and the aqueous layer was extracted with CH_2_Cl_2_ (10 mL × 3). The combined organic layer was dried over Na_2_SO_4_, filtered, and the filtrate was concentrated. The residue was purified by chromatography (hexanes–EtOAc, 2:3) to afford **20** (44 mg, 70%) as a white foam. *R*_f_ 0.15 (hexanes– EtOAc, 1:1); [α]_D_ –18.6 (*c* 0.43, CHCl_3_); ^1^H NMR (600 MHz, CDCl_3_, δ_H_) 8.10–8.04 (m, 7H), 8.03–7.95 (m, 9H), 7.94–7.80 (m, 26H), 7.78–7.68 (m, 26H), 7.66–7.58 (m, 5H), 7.52–7.29 (m, 55H), 7.28–7.17 (m, 29H), 7.15–7.00 (m, 23H), 5.98–5.94 (m, 1H), 5.89–5.81 (m, 4H), 5.80–5.66 (m, 12H), 5.66–5.61 (m, 7H), 5.59–5.45 (m, 12H), 5.38 (d, *J* = 4.2 Hz, 1H), 5.11 (d, *J* = 4.4 Hz, 1H), 5.08 (app t, *J* = 6.4 Hz, 1H), 4.93 (app t, *J* = 6.4 Hz, 1H), 4.89–4.83 (m, 3H), 4.75–4.62 (m, 5H), 4.61–4.38 (m, 19H), 4.36–4.20 (m, 9H), 4.15–4.11 (m, 5H), 4.05–3.96 (m, 4H), 3.65–3.63 (m, 1H, OC<u>H_2_</u>CH_2_), 3.41–3.34 (m, 1H, OC<u>H_2_</u>CH_2_), 3.22 (t, *J* = 7.0 Hz, 2H, C<u>H_2_</u>N_3_), 1.59–1.50 (m, 4H, octyl CH_2_), 1.36–1.30 (m, 8H, octyl CH_2_); ^13^C NMR (151 MHz, CDCl_3_, δ_C_) 166.2, 166.0, 165.84, 165.82, 165.63, 165.61, 165.49, 165.47, 165.5, 165.44, 165.41, 165.3, 165.2, 165.14, 165.11, 165.0, 164.92, 164.88, 164.69, 164.66, 133.5, 133.32, 133.28, 133.2, 133.1, 133.0, 132.9, 132.8, 132.7, 132.6, 130.1, 130.0, 129.9, 129.81, 129.77, 129.7, 129.64, 129.59, 129.53, 129.48, 129.2, 129.1, 129.04, 129.01, 128.97, 128.93, 128.89, 128.88, 128.81, 128.77, 128.7, 128.6, 128.52, 128.50, 128.45, 128.4, 128.32, 128.29, 128.25, 128.21, 128.15, 128.0, 105.7, 105.60, 105.55, 105.5, 105.4, 105.0, 104.9, 101.1, 101.01, 101.00, 100.4, 83.2, 83.1, 82.7, 82.5, 82.2, 82.1, 81.94, 81.86, 81.8, 81.7, 81.6, 81.5, 81.4, 77.8, 77.72, 77.65, 76.7, 76.63, 76.61, 76.5, 76.4, 74.1, 73.92, 73.86, 73.7, 73.5, 73.4, 73.13, 73.07, 73.0, 72.9, 72.1, 72.0, 71.9, 71.80, 71.75, 71.6, 71.23, 71.22, 71.19, 70.6, 69.9, 69.7, 69.6, 69.5, 67.1, 66.2, 65.9, 65.1, 63.2, 62.9, 62.8, 62.6, 51.4, 29.7, 29.3, 29.1, 28.8, 26.7, 26.0; HRMS (ESI) Calcd for (M + Na) C_332_H_281_O_97_N_3_Na: 5887.8491. Found 5886.8447.

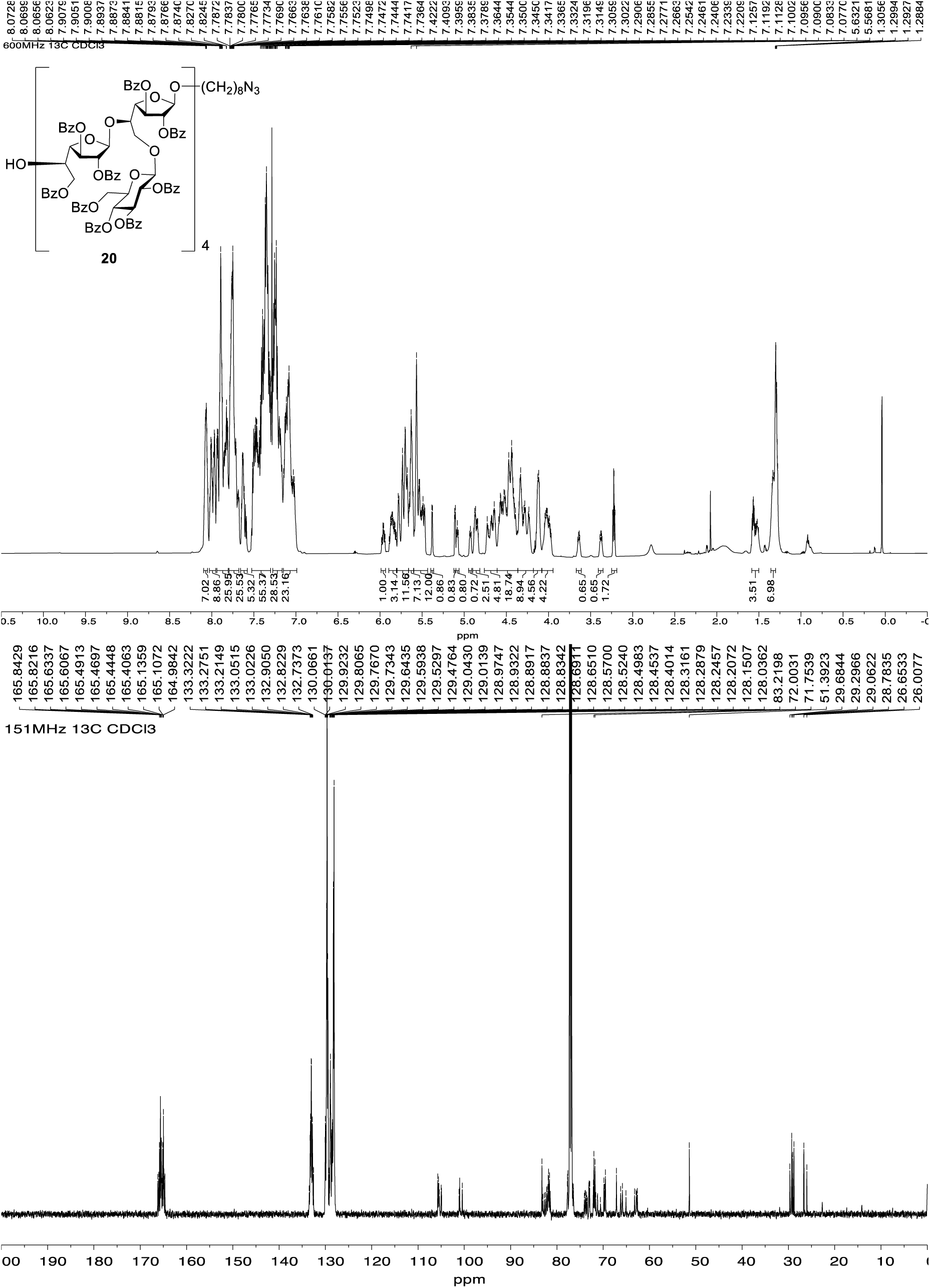

#### 8-Azidooctyl 2,3,6-tri-*O*-benzoyl-5-*O*-levulinoyl-β-D-galactofuranosyl-(1→5)-[2,3,4,6-tetra-*O*-benzoyl-β-D-glucopyranosyl-(1→6)]-2,3-di-*O*-benzoyl-β-D-galactofuranosyl-(1→5)-2,3,6-tri-*O*-benzoyl-β-D-galactofuranosyl-(1→5)-[2,3,4,6-tetra-*O*-benzoyl-β-D-glucopyranosyl-(1→6)]-2,3-di-*O*-benzoyl-β-D-galactofuranosyl-(1→5)-2,3,6-tri-*O*-benzoyl-β-D-galactofuranosyl-(1→5)-[2,3,4,6-tetra-*O*-benzoyl-β-D-glucopyranosyl-(1→6)]-2,3-di-*O*-benzoyl-β-D-galactofuranosyl-(1→5)-2,3,6-tri-*O*-benzoyl-β-D-galactofuranosyl-(1→5)-[2,3,4,6-tetra-*O*-benzoyl-β-D-glucopyranosyl-(1→6)]-2,3-di-*O*-benzoyl-β-D-galactofuranosyl-(1→5)-2,3,6-tri-*O*-benzoyl-β-D-galactofuranosyl-(1→5)-[2,3,4,6-tetra-*O*-benzoyl-β-D-glucopyranosyl-(1→6)]-2,3-di-*O*-benzoyl-β-D-galactofuranoside (21)

A mixture of alcohol **20** (40 mg, 6.8 μmol), glycosyl fluoride **24** (53 mg, 34 μmol) and 4 Å molecular sieves (powder, 0.40 g) in CH_2_Cl_2_ (1.2 mL) was stirred at room temperature for 30 min and then cooled to 0 °C before bis(cyclopentadienyl)zirconium (IV) dichloride (10 mg, 34 μmol) and silver trifluoromethanesulfonate (18 mg, 68 μmol) were added successively. After stirring for 4h at rt, triethylamine was added and the resulting mixture was filtered through Celite. The filtrate was diluted with CH_2_Cl_2_ (10 mL) and washed with satd NaHCO_3_ soln. The organic layer was separated and the aqueous layer was extracted with CH_2_Cl_2_ (10 mL × 3). The combined organic layers were dried over Na_2_SO_4_, filtered, the filtrate was concentrated and the residue was purified by chromatography (hexanes– acetone, 1:1.17). The resulting compounds was purified again by LH-20 size exclusion chromatography (180 mL, CH_2_Cl_2_–CH_3_OH, 1:3) to afford **21** (43 mg, 87%) as a white foam. *R*_f_ 0.3 (hexanes–acetone, 1:1.17); [α]_D_ –12.9 (*c* 1.63, CHCl_3_); ^1^H NMR (600 MHz, CDCl_3_, δ_H_) δ 8.15–7.99 (m, 10H), 7.97–7.95 (m, 4H), 7.94–7.52 (m, 78H), 7.51–7.27 (m, 61H), 7.25–6.92 (m, 72H), 5.91 (app t, *J* = 9.5 Hz, 1H), 5.84–5.76 (m, 4H), 5.76–5.71 (m, 2H), 5.71–5.65 (m, 8H), 5.64–5.56 (m, 13H), 5.55–5.43 (m, 15H), 5.40 (d, *J* = 1.9 Hz, 1H), 5.34–5.33 (m, 2H), 5.06 (s, 1H), 5.03 (d, *J* = 7.8 Hz, 1H), 4.88 (d, *J* = 7.8 Hz, 1H), 4.86–4.76 (m, 4H), 4.73–4.57 (m, 6H), 4.57–4.46 (m, 10H), 4.44–4.34 (m, 15H), 4.33–4.26 (m, 6H), 4.25–4.16 (m, 4H), 4.11–4.04 (m, 4H), 4.04–3.96 (m, 4H), 3.95–3.92 (m, 3H), 3.59 (dt, *J* = 9.6, 6.5 Hz, 1H, OC<u>H_2_</u>CH_2_), 3.33 (dt, *J* = 9.5, 6.3 Hz, 1H, OC<u>H_2_</u>CH_2_), 3.19 (t, *J* = 7.0 Hz, 2H, <u>CH_2_</u>N_3_), 2.53–2.11 (m, 1H, Lev CH_2_), 2.39 – 2.24 (m, 2H, Lev CH_2_), 2.20 – 2.03 (m, 1H, Lev CH_2_), 1.93 (s, 3H, Lev CH_3_), 1.56–1.37 (m, 4H, octyl CH_2_), 1.31–1.23 (m, 8H, octyl CH_2_); ^13^C NMR (151 MHz, CDCl_3_, δ_C_) δ 206.0, 165.9, 165.8, 165.6, 165.50, 165.45, 165.4, 165.3, 165.1, 165.0, 164.7, 133.3, 133.2, 133.0, 132.8, 132.6, 130.1, 130.0, 129.9, 129.8, 129.74, 129.71, 129.66, 129.6, 129.54, 129.49, 129.4, 129.3, 129.2, 129.04, 128.97, 128.9, 128.71, 128.66, 128.6, 128.5, 128.4, 128.32, 128.29, 128.23, 128.16, 128.1, 128.0, 105.83, 105.79, 105.7, 105.60, 105.59, 105.5, 105.4, 105.3, 105.0, 104.9, 101.1, 101.0, 100.4, 83.20, 83.15, 82.78, 82.75, 82.5, 82.4, 82.2, 82.0, 81.9, 81.8, 81.71, 81.67, 81.6, 81.5, 77.9, 77.82, 77.75, 77.5, 76.6, 76.42, 76.40, 76.3, 76.2, 74.3, 74.2, 74.0, 73.7, 73.58, 73.56, 73.5, 73.4, 73.2, 73.11, 73.08, 72.9, 72.1, 72.03, 72.02, 71.80, 71.76, 70.1, 69.9, 69.7, 69.63, 69.58, 69.4, 67.1, 65.92, 65.86, 65.1, 63.6, 63.2, 62.9, 62.8, 62.7, 62.6, 51.4, 37.9, 29.6, 29.3, 29.1, 28.8, 27.8, 26.7, 26.0; HRMS (ESI) Calcd for (M + Na) C_418_H_353_O_123_N_3_Na: 7409.3550. Found 7409.3800.

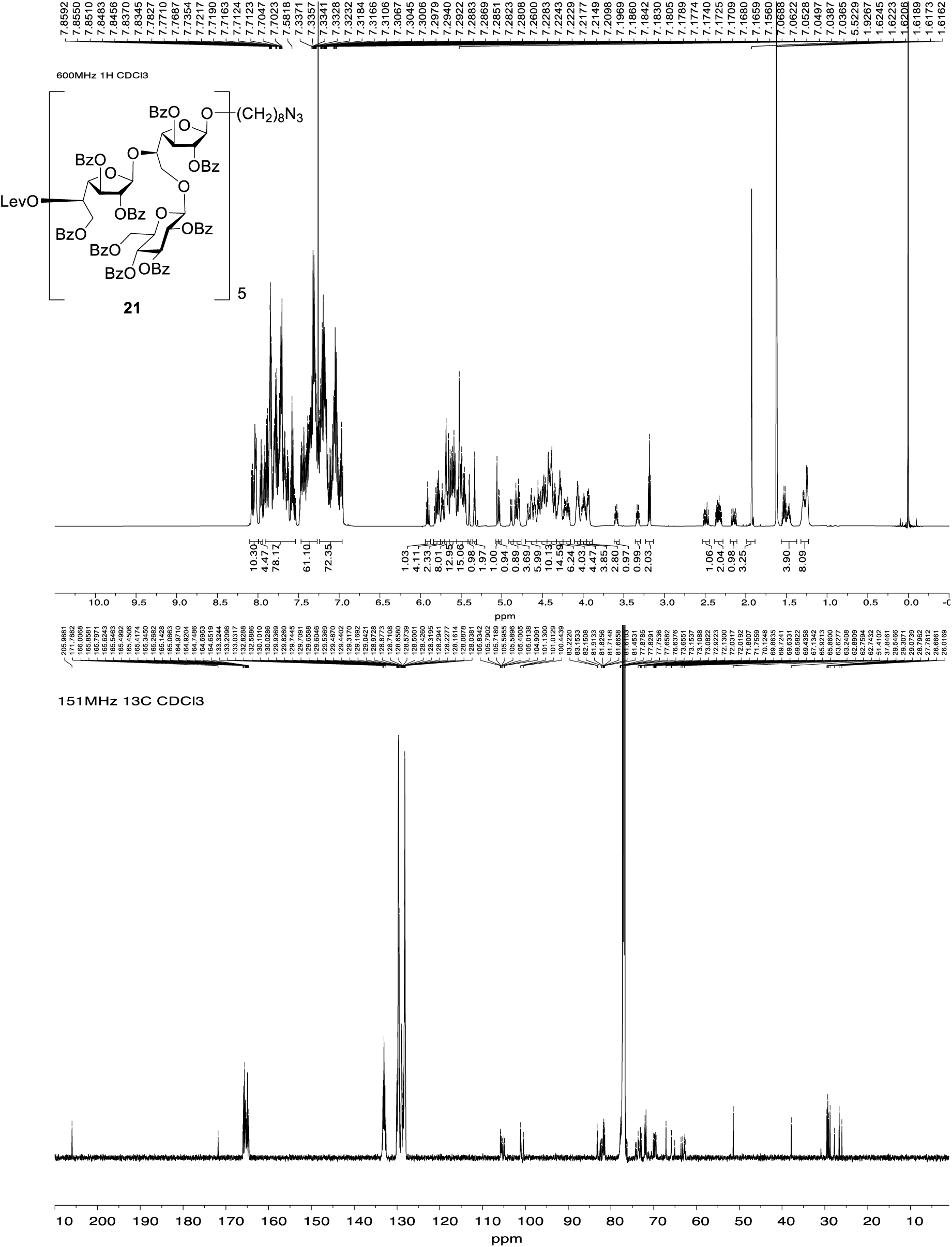

#### 8-Azidooctyl 2,3,6-tri-*O*-benzoyl-β-D-galactofuranosyl-(1→5)-[2,3,4,6-tetra-*O*-benzoyl-β-D-glucopyranosyl-(1→6)]-2,3-di-*O*-benzoyl-β-D-galactofuranosyl-(1→5)-2,3,6-tri-*O*-benzoyl-β-D-galactofuranosyl-(1→5)-[2,3,4,6-tetra-*O*-benzoyl-β-D-glucopyranosyl-(1→6)]-2,3-di-*O*-benzoyl-β-D-galactofuranosyl-(1→5)-2,3,6-tri-*O*-benzoyl-β-D-galactofuranosyl-(1→5)-[2,3,4,6-tetra-*O*-benzoyl-β-D-glucopyranosyl-(1→6)]-2,3-di-*O*-benzoyl-β-D-galactofuranosyl-(1→5)-2,3,6-tri-*O*-benzoyl-β-D-galactofuranosyl-(1→5)-[2,3,4,6-tetra-*O*-benzoyl-β-D-glucopyranosyl-(1→6)]-2,3-di-*O*-benzoyl-β-D-galactofuranosyl-(1→5)-2,3,6-tri-*O*-benzoyl-β-D-galactofuranosyl-(1→5)-[2,3,4,6-tetra-*O*-benzoyl-β-D-glucopyranosyl-(1→6)]-2,3-di-*O*-benzoyl-β-D-galactofuranoside (22)

Pentadecasaccharide **21** (20 mg, 2.8 μmol) and hydrazine acetate (0.3 mg, 3.4 μmol) were dissolved in CH_2_Cl_2_–CH_3_OH (0.3:0.1 mL). After stirring at room temperature for 4 h, the resulting mixture was diluted with CH_2_Cl_2_ (10 mL) and washed with satd NaCl soln. The organic layer was separated and the aqueous layer was extracted with CH_2_Cl_2_ (10 mL × 3). The combined organic layer was dried over Na_2_SO_4_, filtered, the filtrate was concentrated. The residue was purified by chromatography (hexanes– acetone, 1:1.17) to give a compound that was purified again by LH-20 size exclusion chromatography (180 mL, CH_2_Cl_2_–CH_3_OH, 1:3) affording **22** (16 mg, 87%) as a white foam. *R*_f_ 0.38 (hexanes–acetone, 1:1.17); [α]_D_ –8.2 (*c* 1.1, CHCl_3_); ^1^H NMR (600 MHz, CDCl_3_, δ_H_) 8.05–7.99 (m, 10H), 7.98–7.90 (m, 9H), 7.90–7.82 (m, 21H), 7.81–7.76 (m, 12H), 7.75–7.62 (m, 34H), 7.61–7.53 (m, 7H), 7.49–7.27 (m, 60H), 7.24–6.94 (m, 72H), 5.91 (app t, *J* = 9.5 Hz, 1H), 5.85–5.71 (m, 6H), 5.71–5.55 (m, 20H), 5.55–5.40 (m, 16H), 5.33 (d, *J* = 1.6 Hz, 1H), 5.06 (s, 1H), 5.03 (d, *J* = 7.7 Hz, 1H), 4.88 (d, *J* = 7.8 Hz, 1H), 4.85–4.77 (m, 4H), 4.71–4.56 (m, 6H), 4.56–4.33 (m, 23H), 4.33–4.15 (m, 12H), 4.12–3.90 (m, 13H), 3.59 (dt, *J* = 9.6, 6.6 Hz, 1H, OC<u>H_2_</u>CH_2_), 3.33 (dt, *J* = 9.7, 6.3 Hz, 1H, OC<u>H_2_</u>CH_2_), 3.18 (t, *J* = 7.0 Hz, 2H, <u>CH_2_</u>N_3_), 2.72 (d, *J* = 8.5 Hz, 1H, OH), 1.55–1.45 (m, 4H, octyl CH_2_), 1.36–1.28 (m, 8H, octyl CH_2_); ^13^C NMR (151 MHz, CDCl_3_, δ_C_) δ 166.2, 166.0, 165.9, 165.7, 165.51, 165.45, 165.4, 165.3, 165.23, 165.18, 165.15, 165.0, 164.94, 164.90, 164.71, 164.67, 133.32, 133.26, 133.2, 133.0, 132.9, 132.8, 132.7, 132.6, 130.1, 130.04, 130.98, 130.95, 129.8, 129.7, 129.6, 129.5, 129.1, 129.03, 128.97, 128.9, 128.8, 128.7, 128.6, 128.52, 128.47, 128.42, 128.35, 128.3, 128.22, 128.16, 128.10, 128.06, 105.81, 105.77, 105.75, 105.62, 105.60, 105.5, 105.44, 105.41, 105.1, 104.9, 101.2, 101.0, 100.5, 83.24, 83.15, 83.1, 82.8, 82.54, 82.45, 82.23, 82.21, 82.20, 82.01, 81.96, 81.91, 81.86, 81.69, 81.65, 81.5, 77.73, 77.68, 76.5, 74.2, 74.1, 74.0, 73.73, 73.70, 73.62, 73.57, 73.49, 73.45, 73.2, 73.12, 73.07, 73.0, 72.2, 72.1, 71.94, 71.85, 71.8, 71.7, 71.38, 71.35, 71.3, 71.2, 69.9, 69.8, 69.69, 69.65, 69.6, 67.2, 66.3, 65.92, 65.86, 65.1, 63.4, 63.3, 62.9, 62.8, 62.7, 62.8, 51.4, 29.7, 29.3, 29.1, 28.8, 26.7, 26.0; HRMS (ESI) Calcd for (M + Na) C_413_H_347_O_121_N_3_Na:7311.2529. Found 7311.2597.

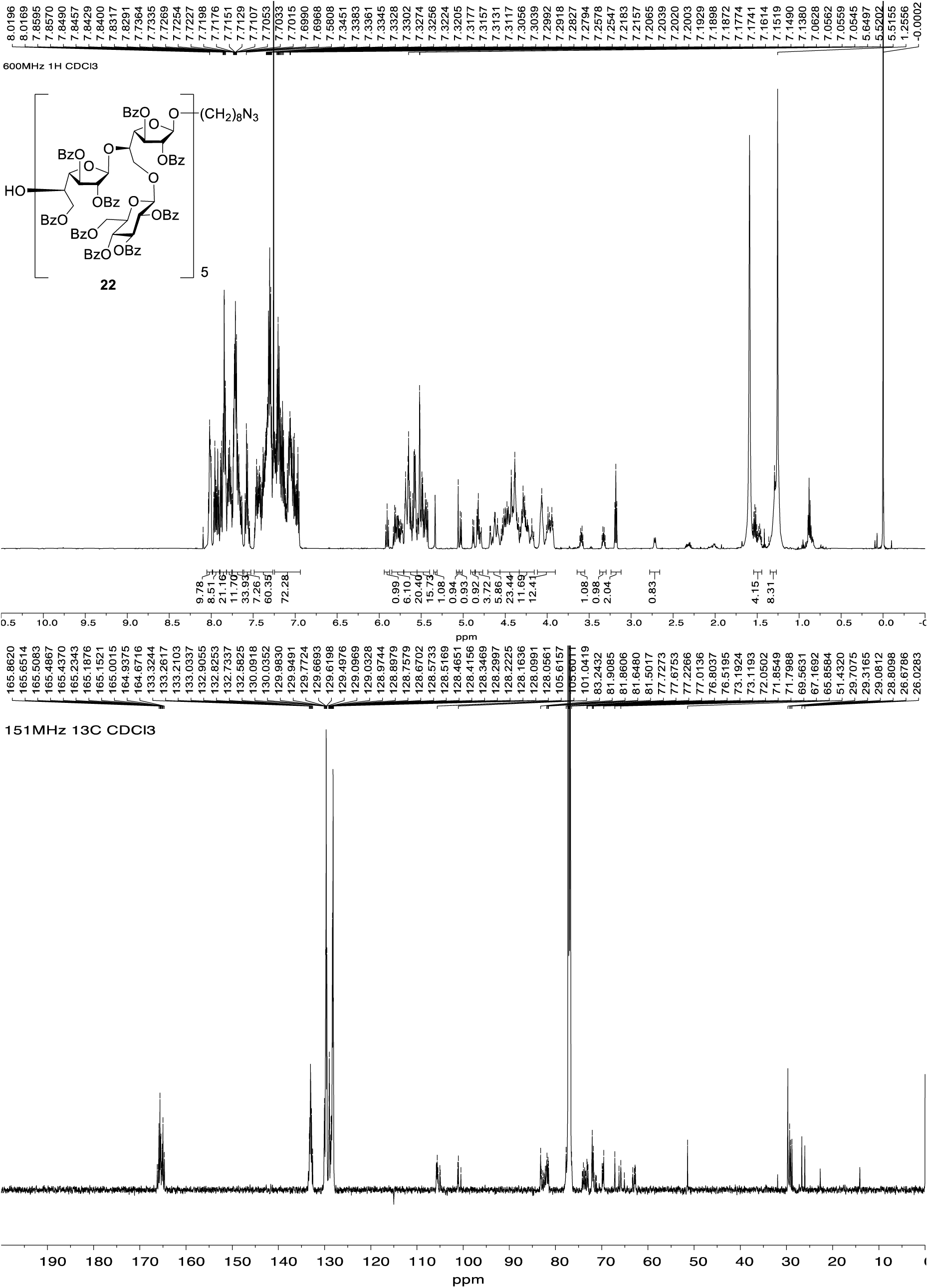

#### 8-Azidooctyl 2,3,6-tri-*O*-benzoyl-5-*O*-levulinoyl-β-D-galactofuranosyl-(1→5)-[2,3,4,6-tetra-*O*-benzoyl-β-D-glucopyranosyl-(1→6)]-2,3-di-*O*-benzoyl-β-D-galactofuranosyl-(1→5)-2,3,6-tri-*O*-benzoyl-β-D-galactofuranosyl-(1→5)-[2,3,4,6-tetra-*O*-benzoyl-β-D-glucopyranosyl-(1→6)]-2,3-di-*O*-benzoyl-β-D-galactofuranosyl-(1→5)-2,3,6-tri-*O*-benzoyl-β-D-galactofuranosyl-(1→5)-[2,3,4,6-tetra-*O*-benzoyl-β-D-glucopyranosyl-(1→6)]-2,3-di-*O*-benzoyl-β-D-galactofuranosyl-(1→5)-2,3,6-tri-*O*-benzoyl-β-D-galactofuranosyl-(1→5)-[2,3,4,6-tetra-*O*-benzoyl-β-D-glucopyranosyl-(1→6)]-2,3-di-*O*-benzoyl-β-D-galactofuranosyl-(1→5)-2,3,6-tri-*O*-benzoyl-β-D-galactofuranosyl-(1→5)-[2,3,4,6-tetra-*O*-benzoyl-β-D-glucopyranosyl-(1→6)]-2,3-di-*O*-benzoyl-β-D-galactofuranosyl-(1→5)-2,3,6-tri-*O*-benzoyl-β-D-galactofuranosyl-(1→5)-[2,3,4,6-tetra-*O*-benzoyl-β-D-glucopyranosyl-(1→6)]-2,3-di-*O*-benzoyl-β-D-galactofuranoside (23)

**A mixture of alcohol** 22 **(16** mg, 2.2 μmol), glycosyl fluoride **24** (16 mg, 11 μmol) and 4 Å molecular sieves (powder, 52 mg) in CH_2_Cl_2_ (0.2 mL) was stirred at room temperature for 30 min and then cooled to 0 °C before bis(cyclopentadienyl)zirconium (IV) dichloride (3.2 mg, 11 μmol) and silver trifluoromethanesulfonate (5.6 mg, 22 μmol) were added successively. After stirring for 4h at rt, triethylamine was added and the resulting mixture was filtered through Celite. The filtrate was diluted with CH_2_Cl_2_ (10 mL) and washed with a satd NaHCO_3_ soln. The organic layer was separated and the aqueous layer was extracted with CH_2_Cl_2_ (10 mL × 3). The combined organic layers were dried over Na_2_SO_4_, filtered and the filtrate was concentrated. The residue was purified by chromatography (hexanes–acetone, 1:1.17) to give a compound that was purified again by LH-20 size exclusion chromatography (180 mL, CH_2_Cl_2_–CH_3_OH, 1:3) to afford **23** (14 mg, 74%) as a white foam. *R*_f_ 0.13 (hexanes–acetone, 1:1.17); [α]_D_ –37.5 (*c* 0.08, CHCl_3_); ^1^H NMR (500 MHz, CDCl_3_, δ_H_) 8.18–7.58 (m, 109H), 7.58–6.95 (m, 161H), 5.94 (app t, *J* = 9.6 Hz, 1H), 5.90–5.46 (m, 51H), 5.43 (s, 1H), 5.37 (d, *J* = 5.7 Hz, 2H), 5.09 (s, 1H), 5.06 (d, *J* = 7.8 Hz, 1H), 4.91 (d, *J* = 7.8 Hz, 1H), 4.90–4.79 (m, 4H), 4.76–4.20 (m, 49H), 4.13–3.90 (m, 15H), 3.72–3.70 (m, 2H, O<u>CH_2_</u>CH_2_), 3.51 (t, *J* = 6.8 Hz, 2H, <u>CH_2_</u>N_3_), 2.60–2.24 (m, 4H, Lev CH_2_), 1.95 (s, 3H, Lev CH_3_), 1.68–1.52 (m, 4H, octyl CH_2_), 1.54–1.35 (m, 8H, octyl CH_2_); ^13^C NMR (126 MHz, CDCl_3_, δ_C_) 206.0, 171.8, 171.2, 166.0, 165.9, 165.6, 165.4, 165.3, 165.2, 165.0, 164.7, 133.2, 133.0, 132.9, 132.6, 130.1, 129.8, 129.7, 129.6, 129.5, 129.1, 128.9, 128.7, 128.7, 128.6, 128.3, 128.2, 128.1, 105.8, 105.7, 105.6, 105.5, 105.0, 101.1, 101.0, 100.5, 99.8, 83.2, 81.8, 81.7, 81.5, 73.2, 73.1, 72.9, 72.5, 72.1, 71.8, 71.8, 71.3, 70.5, 70.2, 69.9, 69.7, 69.6, 69.6, 69.5, 69.5, 67.2, 65.9, 61.9, 60.4, 51.4, 37.9, 29.7, 29.4, 29.3, 29.3; HRMS (ESI) Calcd for (M + Na) C_499_H_419_O_147_N_3_Na: 8832.7598. Found 8832.7477.

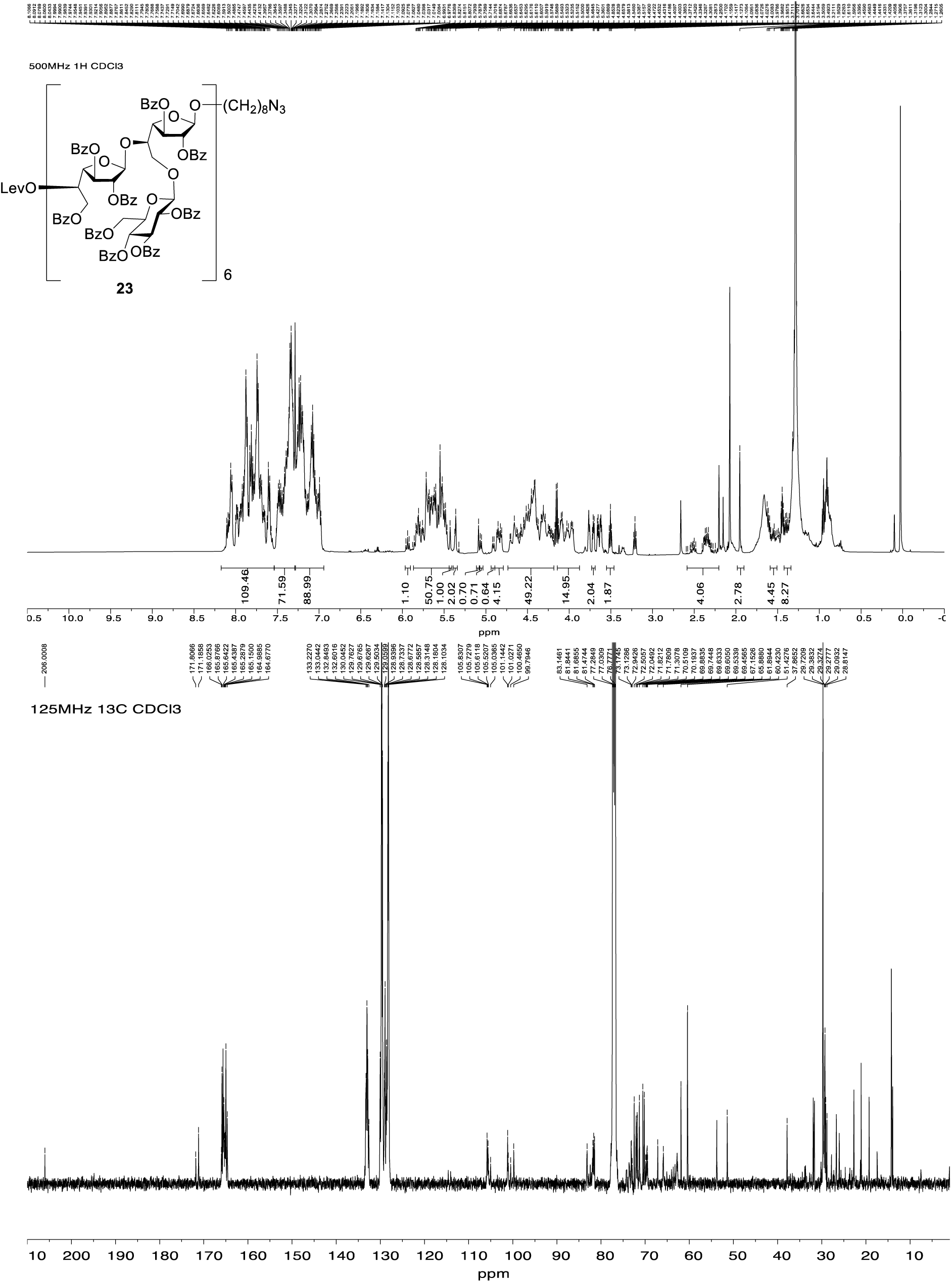

#### 2,3,6-Tri-*O*-benzoyl-5-*O*-levulinoyl-β-D-galactofuranosyl-(1→5)-[2,3,4,6-tetra-*O*-benzoyl-β-D-glucopyranosyl-(1→6)]-2,3-di-*O*-benzoyl-β-D-galactofuranosyl fluoride (24)

To a solution of **12** (0.11 g, 65 μmol) in CH_2_Cl_2_ (1.2 mL) at 0 °C was added *N*,*N*-diethylamino sulfur trifluoride (24 μL, 182 μmol) followed by *N*-bromosuccinimide (33 mg, 185 μmol). The reaction mixture was stirred overnight at room temperature and then CH_3_OH was added. The reaction mixture was diluted with CH_2_Cl_2_ and washed with satd aq NaHCO_3_ soln before the organic layer was separated. The aqueous layer was extracted again with CH_2_Cl_2_ and the combined organic layers were dried over Na_2_SO_4_. The solution was filtered, the filtrate was concentrated and the resulting residue was purified by chromatography (hexanes–acetone, 5:2) to afford glycosyl fluoride **24** (0.1 g, 99%) as a white foam. *R*_f_ 0.13 (hexanes– acetone, 5:2); [α]_D_ +4.4 (*c* 3.2, CHCl_3_); ^1^H NMR (500 MHz, CDCl_3_, δ_H_) 8.13–8.08 (m, 2H, Ar), 7.99–7.91 (m, 8H, Ar), 7.91–7.87 (m, 2H, Ar), 7.82–7.74 (m, 6H, Ar), 7.62–7.53 (m, 2H, Ar), 7.52–7.42 (m, 7H, Ar), 7.42–7.38 (m, 4H, Ar), 7.37–7.29 (m, 10H, Ar), 7.27–7.18 (m, 2H, Ar), 7.15–7.09 (m, 2H, Ar), 5.92 (app t, *J* = 9.6 Hz, 1H, H-3’’), 5.88–5.71 (m, 2H, H-1, H-2), 5.67–5.59 (m, 2H, H-4’’, H-5’), 5.59–5.49 (m, 4H, H-1’, H-2’, H-3, H-2’’), 5.46–5.42 (m, 1H, H-3’), 5.04 (d, *J* = 7.8 Hz, 1H, H-1’’), 4.76 (app t, *J* = 4.8 Hz, 1H, H-4’), 4.69–4.58 (m, 2H, H-4, H-6b’), 4.56–4.78 (m, 2H, H-6b’’, H-5), 4.44–4.33 (m, 2H, H-6a’’, H-6a’), 4.24 (dd, *J* = 11.0, 5.2 Hz, 1H, H-6b), 4.18 (ddd, *J* = 9.9, 5.4, 3.2 Hz, 1H, H-5’’), 4.14–4.05 (m, 1H, H-6a), 2.66–2.36 (m, 4H, Lev CH_2_), 2.01 (s, 3H, Lev CH_3_); ^13^C NMR (126 MHz, CDCl_3_, δ_C_) 206.0 (Lev ketone C=O), 171.9 (Lev ester C=O), 166.0 (Ar), 165.9 (Ar), 165.7 (Ar), 165.43 (Ar), 165.37 (Ar), 165.3 (Ar), 165.14 (Ar), 165.08 (Ar), 164.9 (Ar), 133.7 (Ar), 133.60 (Ar), 133.55 (Ar), 133.4 (Ar), 133.3 (Ar), 133.2 (Ar), 133.1 (Ar), 130.1 (Ar), 129.9 (Ar), 129.8 (Ar), 129.74 (Ar), 129.71 (Ar), 129.66 (Ar), 129.6 (Ar), 129.5 (Ar), 129.1 (Ar), 129.0 (Ar), 128.79 (Ar), 128.76 (Ar), 128.7 (Ar), 128.61 (Ar), 128.55 (Ar), 128.5 (Ar), 128.4 (Ar), 128.3 (Ar), 128.2 (Ar), 128.24 (Ar), 128.21 (Ar), 112.06 (d, *J* = 225.8 Hz, C-1), 104.8 (C-1’), 100.7 (C-1’’), 85.0 (C-4), 81.8 (C-4’), 81.4 (C-2’), 80.9 (d, *J* = 39.3 Hz, C-2), 77.1 (C-5), 75.7 (C-3’’), 73.2 (C-2’’), 72.8 (C-5’’), 72.3 (C-5’), 71.7 (C-4’’), 70.1 (C-, 69.7 (C-3), 67.7(C-3’), 63.2 (C-6’), 63.11 (C-6’’), 37.9 (Lev CH_2_), 29.6 (Lev CH_2_), 27.9 (Lev CH_3_); ^19^F NMR (470 MHz, CDCl_3_, δ_F_) –124.28; HRMS (ESI) Calcd for (M + Na) C_86_H_73_FNaO_26_: 1563.4266. Found 1563.4272.

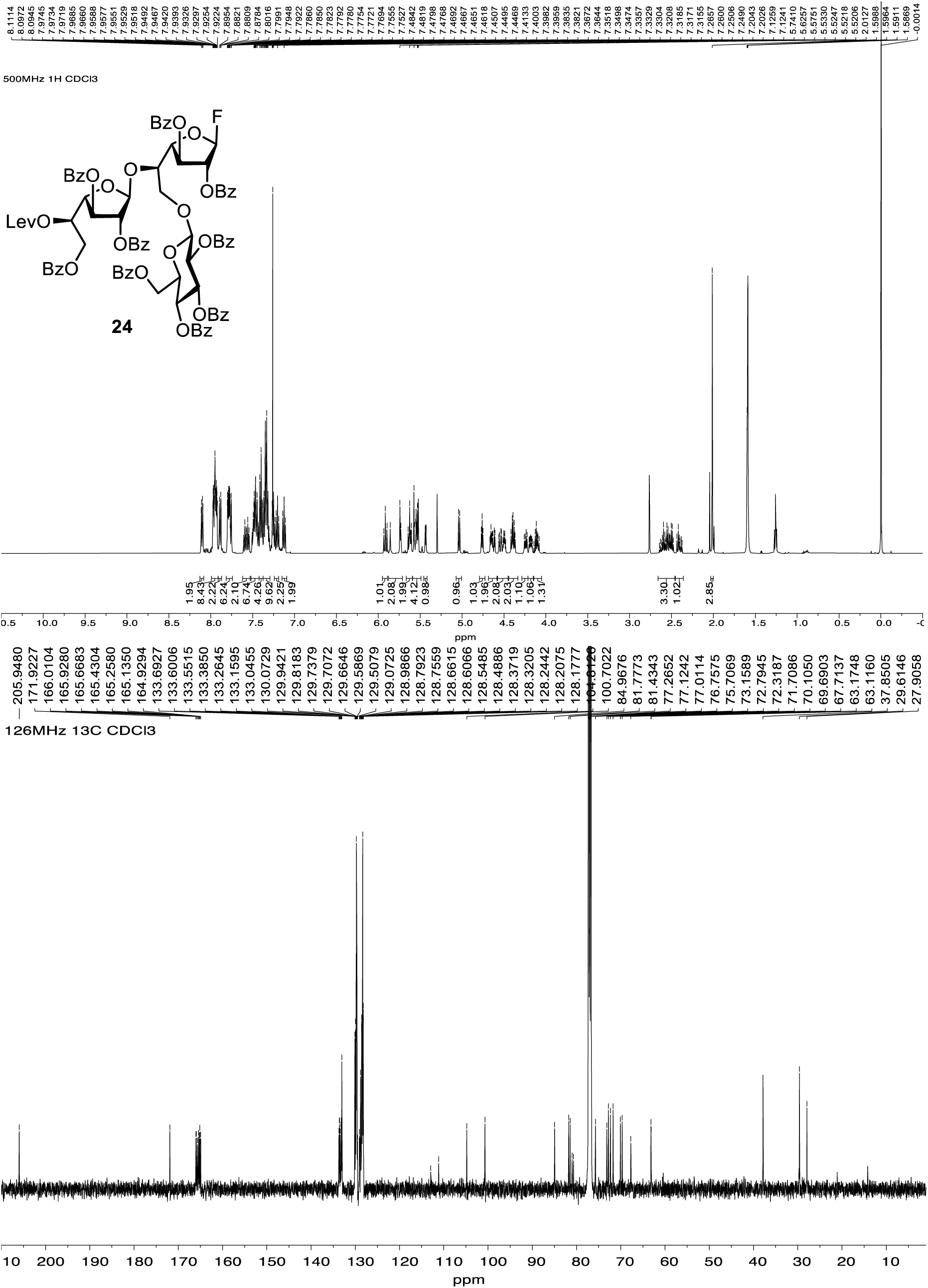

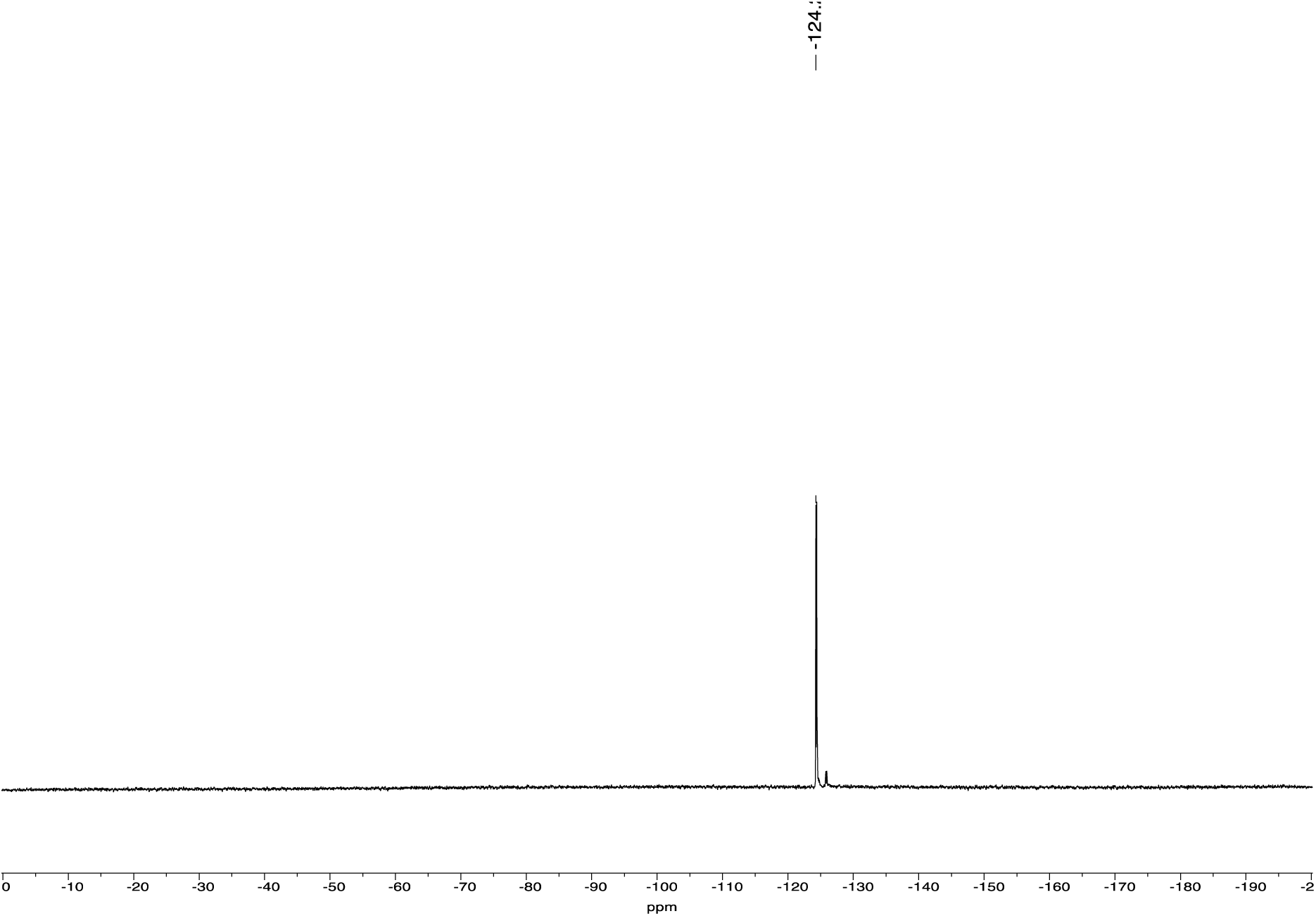

